# Paired-probe chemoproteomic strategy enables global site-resolved quantification of lysine redox states

**DOI:** 10.64898/2026.11.03.764013

**Authors:** Yuan-Fei Zhou, Ling Zhang, D. M. Nirosh Udayanga, Xin Wang, Hasim Bakbak, Hemendra N. Shah, Nima Sharifi, Zhipeng A. Wang

## Abstract

Site-specific quantification of lysine (Lys) redox states remains unexplored, as current approaches identify only a limited number of modification sites without determining their stoichiometry. Allysine, an oxidized form of Lys, provides a model posttranslational modification (PTM) for addressing this challenge by requiring selective enrichment and site-specific stoichiometric quantification. Here, we developed a quantitative chemoproteomic platform for **p**aired **r**edox-state **i**sotopic **s**toichiometry **m**apping of **lys**ine, termed **PRISM-Lys**. We used an Am-alkyne probe for allysine enrichment by reductive amination and developed a complementary Ald-alkyne probe to label Lys, with identical reactions generating the same chemical product. Coupling this paired-probe strategy with a quantitative chemoproteomic workflow enables global site-resolved quantification and stoichiometric analysis of Lys redox states across cells and tissues. Applied to ferroptosis, PRISM-Lys revealed bidirectional, site-selective remodeling of Lys oxidation events associated with ferroptotic progression. Together, PRISM-Lys establishes a paired-chemistry framework for global, site-resolved quantification of Lys redox states, providing a new quantitative perspective for investigating protein oxidation in biological systems.

## Introduction

Posttranslational modifications (PTMs) alter the chemical properties of proteins and thereby regulate their structure, function, and molecular interactions.^1–3^ Proteomics is a powerful approach for identifying PTM sites and monitoring their dynamic changes.^4–6^ However, the analysis of low-abundance PTMs remains challenging for two major reasons. First, modified peptides are often masked by the overwhelming excess of their unmodified counterparts, making selective enrichment essential for sensitive detection.^7, 8^ Second, such enrichment excludes the corresponding unmodified counterparts, limiting access to the information needed to determine site-specific modification stoichiometry, which is a measure that provides insight into PTM dynamics and associated changes in protein function. Accurate determination of PTM stoichiometry therefore requires a strategy that captures and quantifies both modified and unmodified species at the same site simultaneously.^9^ Consequently, current proteomic strategies often face a trade-off: enrichment-based approaches enable profiling of low-abundance PTMs but generally do not provide modification stoichiometry, whereas site-specific stoichiometric analyses have been established mainly for relatively extensively studied PTMs but rarely reported for low-abundance PTMs.^10^

Allysine is an aldehyde-bearing product of lysine (Lys, K) oxidation that can arise through LOX-family-catalyzed oxidative deamination^11^ or nonenzymatic metal-catalyzed oxidation^12^. Beyond its established role in extracellular matrix cross-linking, allysine formation has been implicated in the regulation of chromatin organization and non-histone protein function.^13^ Its accumulation is associated with oxidative stress and has been implicated in diverse pathophysiological conditions, including aging,^14^ fibrosis,^15^ cancer,^16^ and Alzheimer’s disease.^17^ Several site-specific examples illustrate its functional relevance. Allysine at H3K4 contributes to transcriptional repression and heterochromatin compaction and has been linked to epithelial–mesenchymal transition and triple-negative breast cancer.^18, 19^ LOXL2-mediated oxidation of methylated TAF10 regulates TFIID-dependent transcription and the balance between pluripotency and differentiation,^20^ whereas auto-oxidation of LOXL2 K731 suppresses LOXL2 activity and has been implicated in fibrosis and collagen-dependent cancer metastasis.^21^ Despite its emerging biological significance, proteome-wide characterization of allysine remains challenging because of its low abundance and high reactivity toward neighboring amino acid residues. Several strategies, including DNPH labeling ^22, 23^ and capture-and-release approaches,^24, 25^ have been developed to profile allysine modifications. However, the number of identified allysine peptides remains limited and, more importantly, these approaches did not provide allysine stoichiometry. These limitations highlight the need for an integrated strategy that combines sensitive allysine enrichment with site-specific stoichiometric analysis, enabling both broad site coverage and quantitation of allysine stoichiometry.

In this study, we developed **PRISM-Lys**, a quantitative chemoproteomic platform that integrates efficient allysine enrichment with site-specific stoichiometric analysis. PRISM-Lys employs an amine-functionalized alkyne probe (Am-alkyne probe) for allysine enrichment by reductive amination together with a complementary aldehyde-functionalized alkyne probe (Ald-alkyne) for labeling reduced Lys, with identical reactions converging on the same chemical product (Figure 1), which is accomplished by switching the reaction complementarity of the probe and the bait. Coupling this paired chemistry with isotopic quantitative chemoproteomics enables global, site-resolved measurement of allysine stoichiometry across cells and tissues. When we applied PRISM-Lys to ferroptosis, which is an iron-dependent form of regulated cell death characterized by the accumulation of lipid peroxides, it revealed widespread remodeling of Lys redox states and uncovered site-specific oxidation events associated with ferroptotic response. Together, PRISM-Lys establishes an integrated framework for enrichment and stoichiometric quantification of allysine, providing a new quantitative perspective for investigating protein oxidation in biological systems.

**Figure 1.**
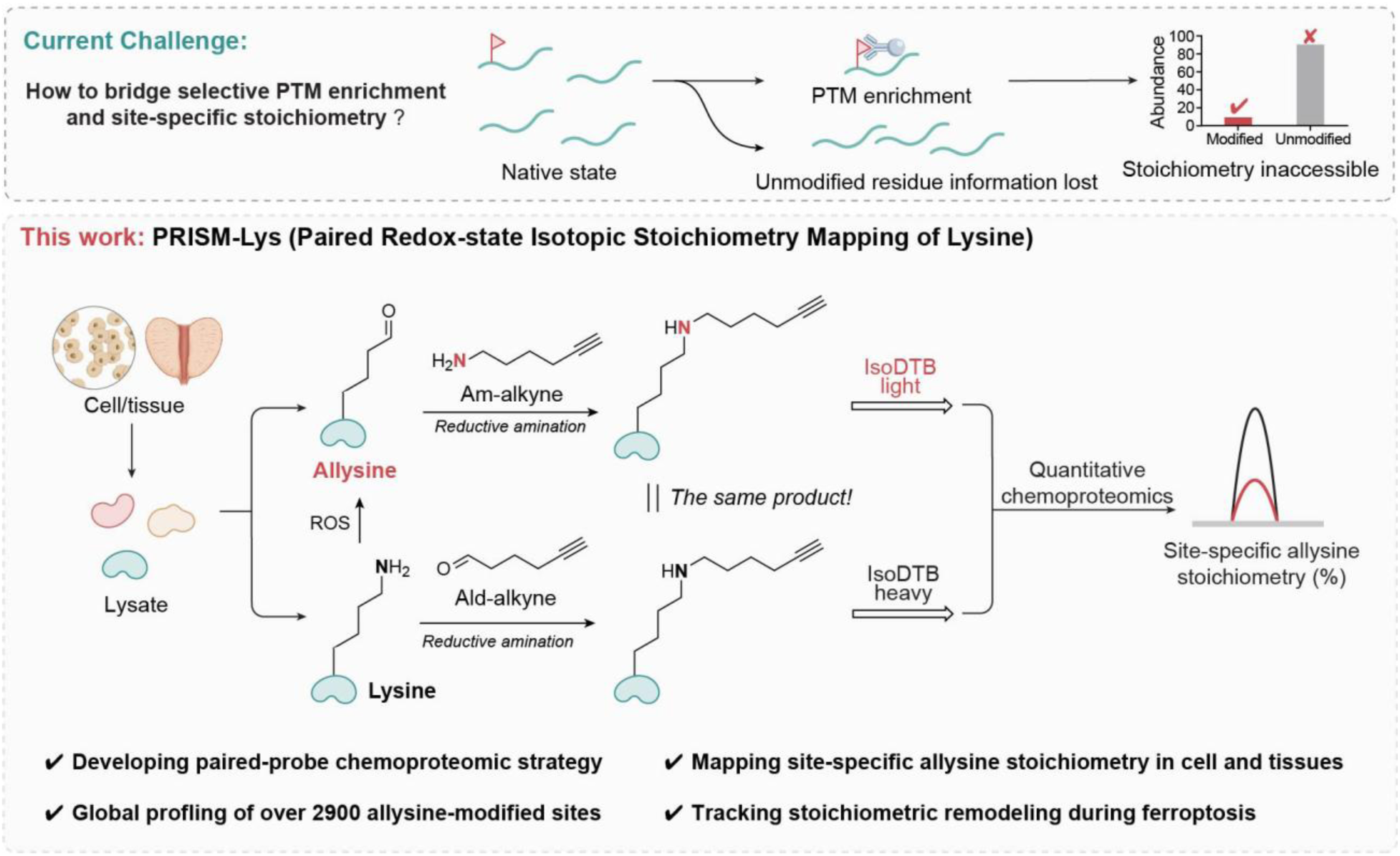
Conceptual framework of PRISM-Lys for site-specific quantification of Lys redox states. Conventional PTM-enrichment strategies selectively capture modified peptides while losing information on the corresponding unmodified residues, precluding direct measurement of site-specific stoichiometry (Top). PRISM-Lys addresses this limitation through complementary reductive-amination reactions that separately label allysine and unmodified Lys. The two states are encoded with light and heavy IsoDTB tags and jointly analyzed by quantitative chemoproteomics, enabling direct measurement of site-specific allysine stoichiometry and global Lys redox-state in cells and tissues and its remodeling during ferroptosis (Bottom).

## Results

### Chemoproteomic study identifies Am-alkyne as a chemoselective probe for allysine enrichment

Bioorthogonal chemical probes provide a powerful platform for interrogating PTMs.^26–30^ These probes typically contain a reactive warhead that selectively targets native PTM sites on endogenous proteins and a bioorthogonal alkyne or azide handle for subsequent click-chemistry conjugation with fluorescent or affinity tags, enabling visualization, enrichment, and proteomic identification of modified proteins and sites.^31, 32^ Previous studies have demonstrated that aldehyde-containing PTMs can be chemically profiled through carbonyl-directed chemistries, including aminooxy ligation^33^ and reductive amination^34^. For example, Wang and colleagues developed an aniline-based probe for HNE-derived protein carbonylation, enabling proteome-wide mapping of HNE-modified proteins and sites.^34^ Separately, reductive amination is a well-established and proteome-compatible reaction in quantitative proteomics, in which formaldehyde reacts with primary amines to form imines that are subsequently reduced by NaBH₃CN.^34, 35^ Together, these established chemistries prompted us to exploit the reactivity between allysine aldehyde and amines. We therefore screened a panel of amine-containing probes (**1**–**4**) (Figure S1a) under reductive amination conditions to identify an efficient probe for allysine capture. In-gel fluorescence analysis following reductive amination and click conjugation with Rho-N₃ did not reveal clear differences among the probes (Figure S1a). We therefore turned to a site-resolved chemoproteomic approach using tandem orthogonal proteolysis-activity-based protein profiling (TOP-ABPP)^36, 37^ to directly evaluate their ability to capture allysine across the proteome (Figure 2a). Specifically, HEK293T cell lysates were labeled with 5 mM amine probes and treated with NaBH₃CN. Following precipitation, the labeled proteomes were conjugated to an acid-cleavable azido-DADPS-biotin tag^38^ via click chemistry (Figure S1b), followed by affinity enrichment, on-bead digestion, and acid-mediated cleavage to release and enrich probe-modified peptides. Successful allysine capture should produce a characteristic adduct on Lys, and with expected mass shifts relative to unmodified Lys of 243.1372 Da for **1**, 223.1685 Da for **2** (Am-alkyne), 181.1215 Da for **3** and 280.1899 Da for **4** (Figure 2b, S1c). First, targeted searches of each chemoproteomic dataset for the expected adduct revealed that Am-alkyne showed marked allysine enrichment, identifying approximately 1805 sites, whereas the other three probes yielded far fewer sites (Figure 2c). Open-search analysis further showed that Am-alkyne produced both the highest number of PTM identifications and the most abundant allysine-derived adduct (Figure 2d, S2a). We manually verified the representative MS2 spectra for the allysine-modified peptides, such as H1.2_K110Allysine (**K**[223.1685]AASGEAKPK) and ENO1_K126Allysine (AGAVE**K**[223.1685]GVPLYR), (Figure S2b) showing high confidence in identification. To further validate allysine enrichment, we first performed the labeling reaction in the absence of NaBH_3_CN and observed no lysine adduct, supporting reductive amination-dependent capture of allysine by Am-alkyne. We then repeated the reductive amination analysis in LNCaP cells by changing DADPS tag to a mixture of light and heavy IsoDTB tag^39, 40^ (Figure S2c), where open-search analysis reproduced the expected allysine-derived mass shifts at 561.3387 Da and 567.3462 Da (Figure S2d, 2e). For example, TUFM_K259Allysine (DLE**K**[IsoDTB tag]PFLLPVEAVYSVPGR) had a matched MS1 pattern (Figure 2f) and MS2 spectra show a shared fragmentation pattern, while the fragment ions containing the IsoDTB-modified allysine exhibit a characteristic 6.01 Da mass shift in the modified b ions between the light and heavy channels (Figure 2f). To improve the robustness and coverage of allysine profiling, we performed additional analyses in HEK293T cells using DADPS tag. Across four HEK293T replicates, we identified 2371 allysine sites, including 731 consistently detected in all four datasets (Figure 2g, top). De-identified benign prostatic hyperplasia (BPH) tissue specimens were collected from patients undergoing holmium laser enucleation of the prostate (HoLEP) and used for allysine enrichment. In human BPH tissues, we identified 1241 allysine sites, of which 794 were reproducibly detected in both datasets (Figure 2g, bottom left). To the best of our knowledge, this method achieves the most comprehensive site-resolved profiles of allysine reported to date, revealing a large number of previously unreported modification sites and proteins (Figure 2g, bottom right). Together, these results establish Am-alkyne as a robust chemoselective reagent for deep, unbiased, site-resolved enrichment of endogenous allysine.

**Figure 2.**
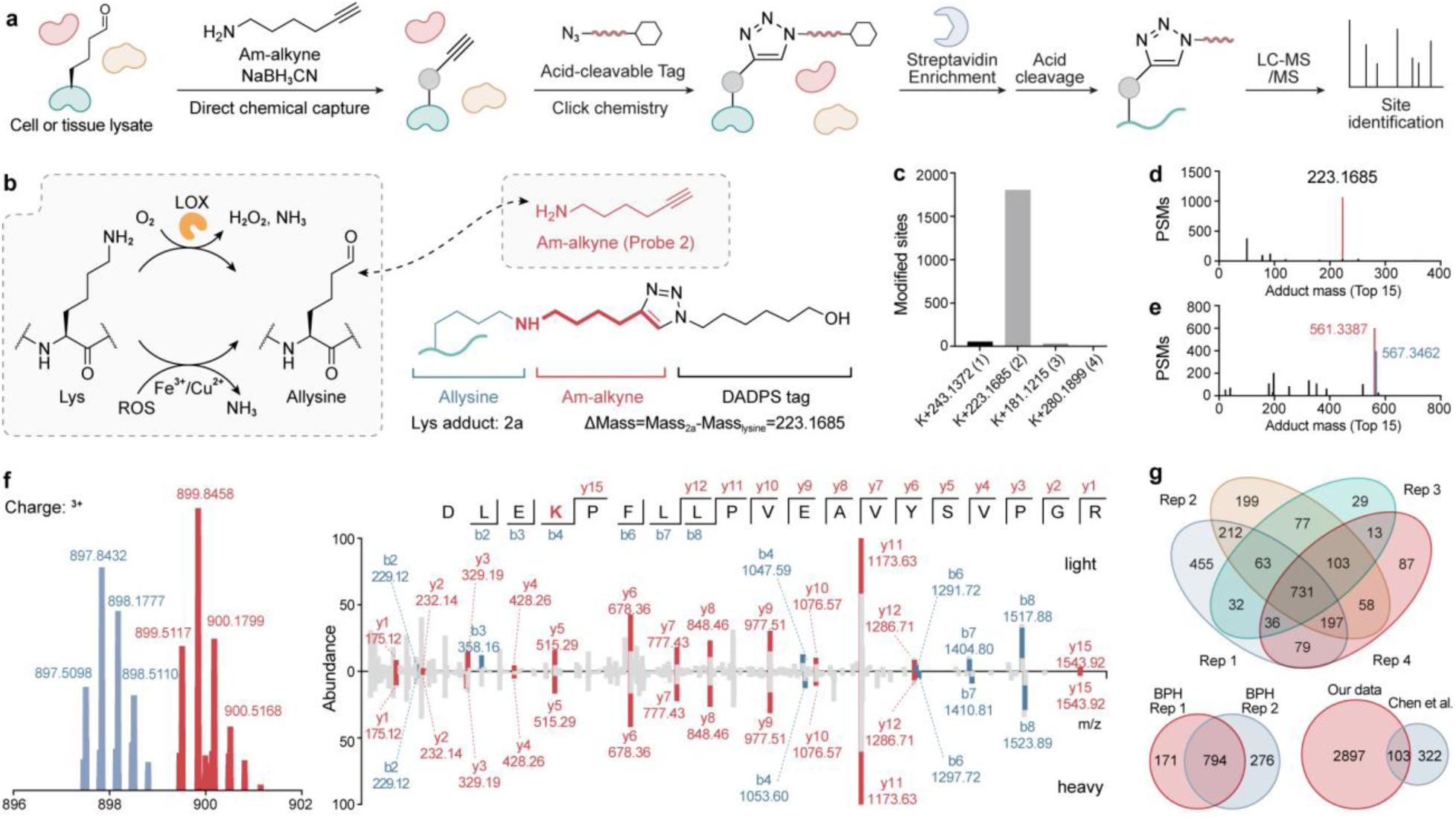
Chemoproteomic characterization identifies Am-alkyne as a chemoselective probe for allysine enrichment. (a) Schematic workflow of TOP-ABPP for chemoproteomic profiling of allysine through reductive amination. (b) Structure of the predicted Lys adduct generated from the reaction of allysine with probe **2** (Am-alkyne). (c) Theoretical mass shifts of the allysine-derived adducts for probes **1**–**4** and the numbers of modification sites identified for each probe. (d) Open-search analysis of Am-alkyne revealed +223.1685 Da as the predominant mass shift, accounting for the largest number of modified PSMs. (e) IsoDTB-based chemoproteomic analysis further identified mass shifts of +561.3387 and +567.3462 Da, corresponding to the expected light- and heavy-IsoDTB-tagged Lys adducts, respectively. (f) Representative MS1 isotopic pattern and MS/MS fragmentation spectrum of TUFM K259Allysine, corresponding to the peptide DLEK[IsoDTB tag]PFLLPVEAVYSVPGR. (g) Venn diagrams showing the overlap of allysine sites identified across four biological replicates (top), the overlap between two replicates in BPH tissue (bottom left), and the overlap between the sites identified in this study and previously reported allysine sites (bottom right).

### Paired-probe chemical design and mass-balanced quantification principle

Determining site-specific PTM stoichiometry, defined as the fraction of a protein population modified at a given residue, remains a major challenge.^9^ This is well illustrated by phosphorylation, where changes in site stoichiometry closely reflect dynamic cellular regulation.^41, 42^ Having established that Am-alkyne selectively captures allysine, we asked whether an analogous probe could be designed to capture Lys in its reduced state. In principle, paired enrichment of allysine and Lys enables global, site-resolved measurement of their relative abundance and thereby defines allysine stoichiometry. To achieve complementary enrichment, the paired probes should rely on the same chemical reaction and generate an identical product. Under these criteria, probe **5** (Ald-alkyne), an aldehyde-containing probe, emerged as an appropriate counterpart to label Lys by reductive amination (Figure 3a). We therefore synthesized Ald-alkyne and evaluated its proteome-wide reactivity using the same reductive amination conditions and TOP-ABPP workflow (Figure S3a). Using the DADPS tag, an open search yielded 14802 peptide-spectrum matches (PSMs) corresponding to the expected 223.1685-Da mass shift, and a subsequent closed search identified 12024 modified Lys sites (Figure 3b, c), demonstrating broad proteome-wide labeling of unmodified Lys residues by Ald-alkyne. To further validate Lys labeling through reductive amination, we replaced NaBH₃CN with NaBD₃CN, which is expected to introduce an additional 1.0063 Da, resulting in a mass shift of 224.1748 Da (Figure S3b). Consistent with this prediction, an open search yielded 16126 PSMs at a mass shift of 224.1747 Da, further supporting the proposed labeling chemistry. Comparison with the allysine dataset revealed 2197 overlapping Lys sites (Figure 3d), highlighting the potential to quantify both the allysine and Lys states at these positions. Furthermore, close searches identified only 101 sites with mass shifts assigned to two Ald-alkyne-derived adducts at 303.2310 and 446.3369 Da (Figure S3c) and shared 51 common sites with allysine (Figure 3d), suggesting that these adducts make a negligible contribution to Lys labeling. We further reasoned that replacing the DADPS tag with the IsoDTB system would enable direct stoichiometry measurement by quantifying its light and heavy adducts at +561.3387 and +567.3462 Da, respectively. Specifically, reductive amination of allysine with Am-alkyne and of Lys with Ald-alkyne under exactly the same condition generates exactly the same chemical adduct (Figure 3e), while subsequent conjugation with light and heavy IsoDTB produces chemically equivalent isotopologues that differ only in mass. Consequently, the two forms are expected to co-elute during LC while remaining distinguishable at the MS1 level, thereby enabling direct quantification of their relative abundance. This chemical convergence design therefore provides the basis for site-resolved quantification of the Lys/allysine pair.

**Figure 3.**
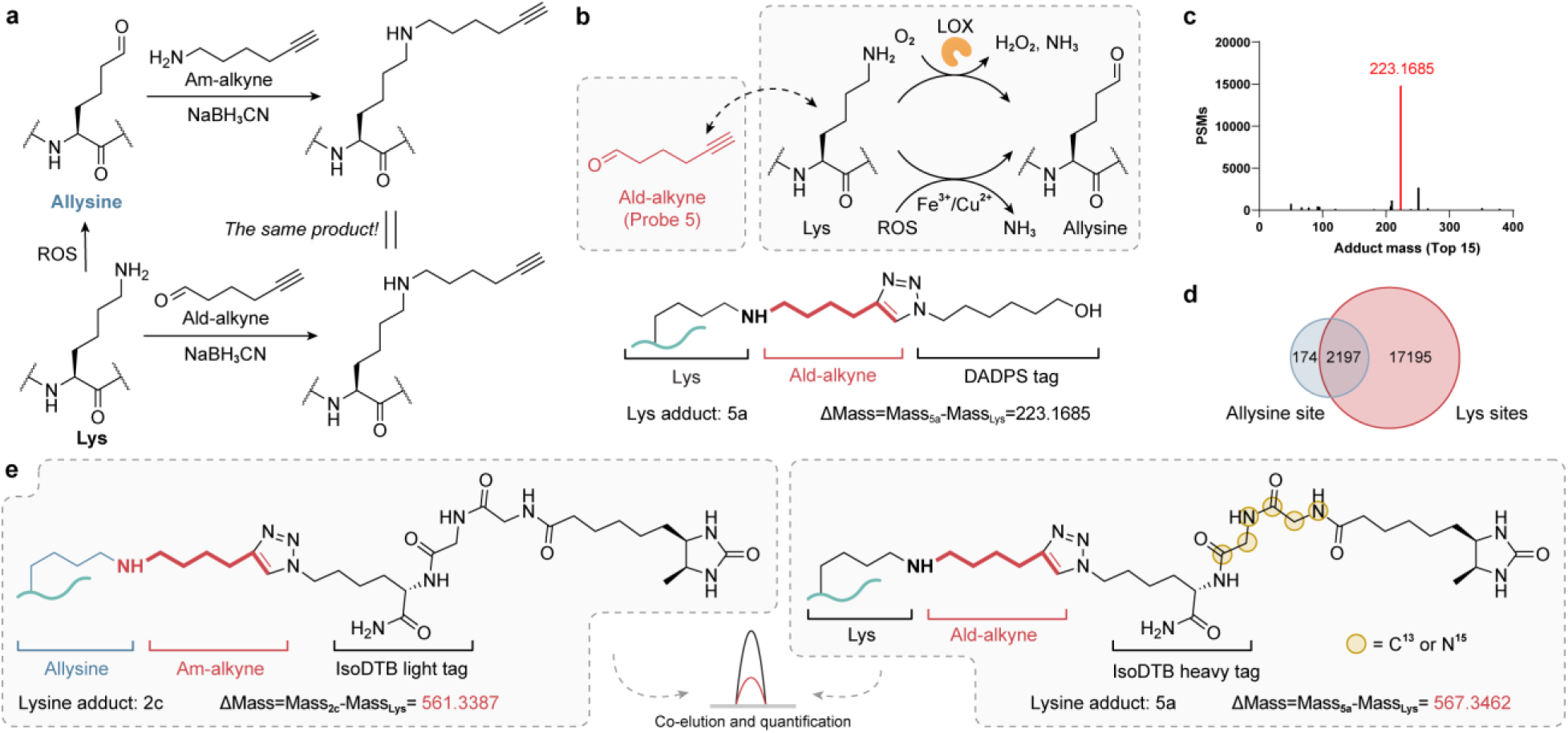
Design of a complementary probe for paired profiling of Lys redox states. (a) Schematic illustration of the paired chemical labeling strategy for Lys redox states. Allysine is captured with amine Am-alkyne, whereas reduced Lys is labeled with aldehyde probe **5** (Ald-alkyne) through reductive amination. (b) Structure of the Lys-derived adduct generated by Ald-alkyne following DADPS tagging, with a theoretical mass shift of +223.1685 Da relative to unmodified Lys. (c) Open-search analysis of Ald-alkyne-labeled proteomic samples identified +223.1685 Da as the predominant mass shift among the most abundant modified PSMs. (d) Venn diagram showing the overlap between Lys sites identified with Ald-alkyne (two replicates) and allysine sites (four replicates) identified with Am-alkyne in HEK293T cells. (e) Structures of the light-IsoDTB-tagged allysine adduct derived from Am-alkyne and the heavy-IsoDTB-tagged Lys adduct derived from Ald-alkyne, corresponding to theoretical mass shifts of +561.3387 and +567.3462 Da, respectively. The paired products are designed to co-elute during LC–MS analysis, enabling direct quantification of the two Lys redox states.

### Paired chemical tagging enables quantitative analysis of allysine in model peptides and proteins

Encouraged by the above chemical design, we next asked whether the paired products could co-elute and accurately determine predefined Lys/allysine ratios in controlled model systems. We first synthesized two Lys-containing peptides, **6a** and **7a**, using solid-phase peptide synthesis (SPPS) (Figure S4a). Following the reported tertiary amine coupling by oxidation (TACO) chemistry,^43, 44^ the corresponding allysine-containing peptides, **6b** and **7b**, were subsequently prepared (Figure S4b). To evaluate the quantitative accuracy of the paired-probe strategy, **6a** and **6b** (Figure 4a) were prepared at defined input amounts, reacted separately with Ald-alkyne and Am-alkyne, respectively, and conjugated with heavy and light IsoDTB tags, and then mixed, followed by LC-MS/MS detection. Doubly charged adducts (Figure S4c) indicated the heavy- and light-labeled products exhibited matching retention times and co-eluted (Figure 4b), supporting our hypothesis in paired-probe chemical design. Measured stoichiometry showed a strong linear relationship with expected values, with modest underestimation. Comparable performance was observed for the **7a**/**7b** peptide pair (Figure 4c, 4d). The two peptide pairs yielded R^2^ values of 0.9932 and 0.9950, with corresponding slopes of 0.9486 and 0.9144, respectively, supporting the quantitative performance of the paired-probe strategy in model peptides.

**Figure 4.**
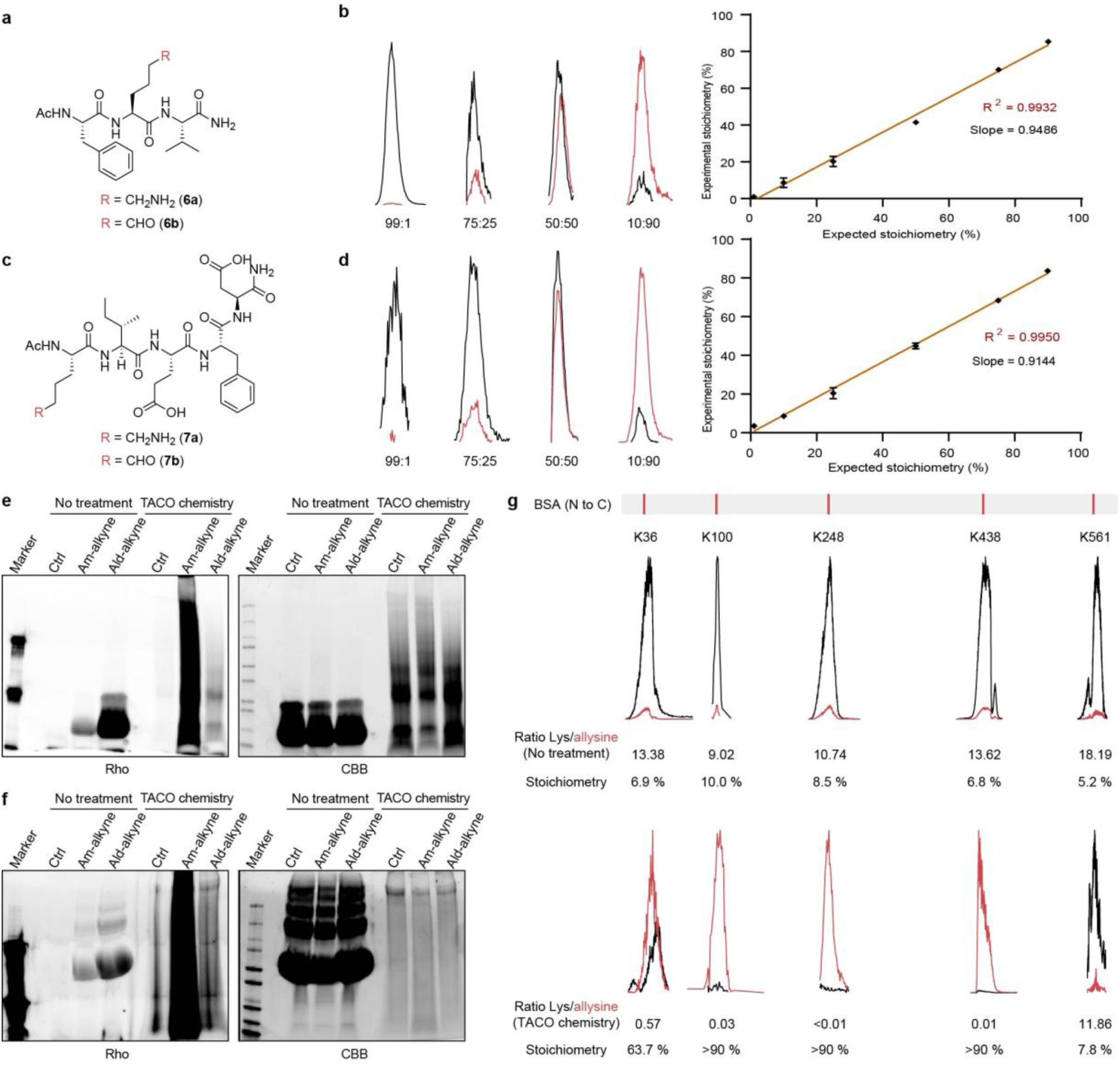
Validation of the paired-probe strategy using model peptides and proteins. (a, c) Structures of the Lys- and allysine-containing model peptide pairs **6a**/**6b** and **7a**/**7b**, respectively. (b, d) Representative extracted ion chromatograms showing co-elution of the paired labeling products at the indicated Lys-to-allysine input ratios, and linear regression of measured versus expected allysine stoichiometry. (e, f) In-gel fluorescence (Rho) and Coomassie Brilliant Blue (CBB) staining of model proteins labeled with amine- or aldehyde-functionalized alkyne probes, with or without prior TACO treatment. Ctrl, control. (g) Representative site-resolved quantification of Lys and allysine in BSA before and after TACO treatment. The positions of five representative Lys residues are indicated along the protein sequence. Black and red traces correspond to Lys- and allysine-derived products, respectively. Allysine stoichiometry was calculated as 100/(1 + R), where R is the measured Lys-to- allysine ratio.

We next evaluated the paired-probe strategy at the protein level using bovine serum albumin (BSA) and lysozyme as model proteins. In their native states, both proteins exhibited substantially stronger labeling with Ald-alkyne than with Am-alkyne (Figure 4e), consistent with the expected predominance of the reduced Lys state. In contrast, following TACO-mediated oxidation, which increases the abundance and stoichiometry of allysine, labeling with Am-alkyne became markedly stronger than that with Ald-alkyne, as expected (Figure 4f). These results demonstrate that the two probes can report shifts in the relative abundance of unmodified Lys and oxidized allysine states at the protein level. TACO-treated proteins also exhibited pronounced band smearing, consistent with increased molecular heterogeneity following extensive protein oxidation. ^43, 44^ Finally, we applied the paired probes to site-resolved quantification of native and TACO-treated BSA (Figure S5a, S5b). Several Lys residues, including K36, K100, K248, K438, and K561, were predominantly present in the reduced Lys state under native conditions (Figure 4g, S5c, S5d). Following TACO-mediated oxidation, K36, K100, K248, and K438 shifted markedly toward the allysine state, with allysine becoming the predominant species, whereas K561 showed only a modest increase in allysine stoichiometry (Figures 4g, S5c, and S5d). Together, these results establish that the Am-alkyne and Ald-alkyne pair enables site-resolved stoichiometry of Lys oxidation and accurately captures changes at individual residues.

### PRISM-Lys for global, site-resolved quantification of Lys redox states across cells and tissues

Given the quantitative accuracy of the Am-alkyne and Ald-alkyne pair demonstrated in both peptide and protein model systems, we next extended this paired-probe strategy to proteome-wide analysis. We termed this strategy **PRISM-Lys** (Paired Redox-state Isotopic Stoichiometry Mapping of Lys), which enables global, site-resolved quantification of allysine stoichiometry. Specifically, the same proteome sample was first divided into two equal aliquots (Figure 5a). One aliquot was labeled with Am-alkyne to capture the oxidized allysine state, with the other labeled with Ald-alkyne to capture the unmodified Lys state. The two samples were then independently subjected to click conjugation with light and heavy IsoDTB tags, respectively. Following click reaction, the samples were combined and processed together through affinity enrichment, proteolytic digestion, and acid/acetonitrile-mediated release from the beads. The resulting isotopically encoded peptides were subsequently analyzed by LC-MS/MS, enabling direct measurement of the relative abundance of the allysine and Lys states at individual Lys sites and thereby calculation of site-specific allysine stoichiometry.

**Figure 5.**
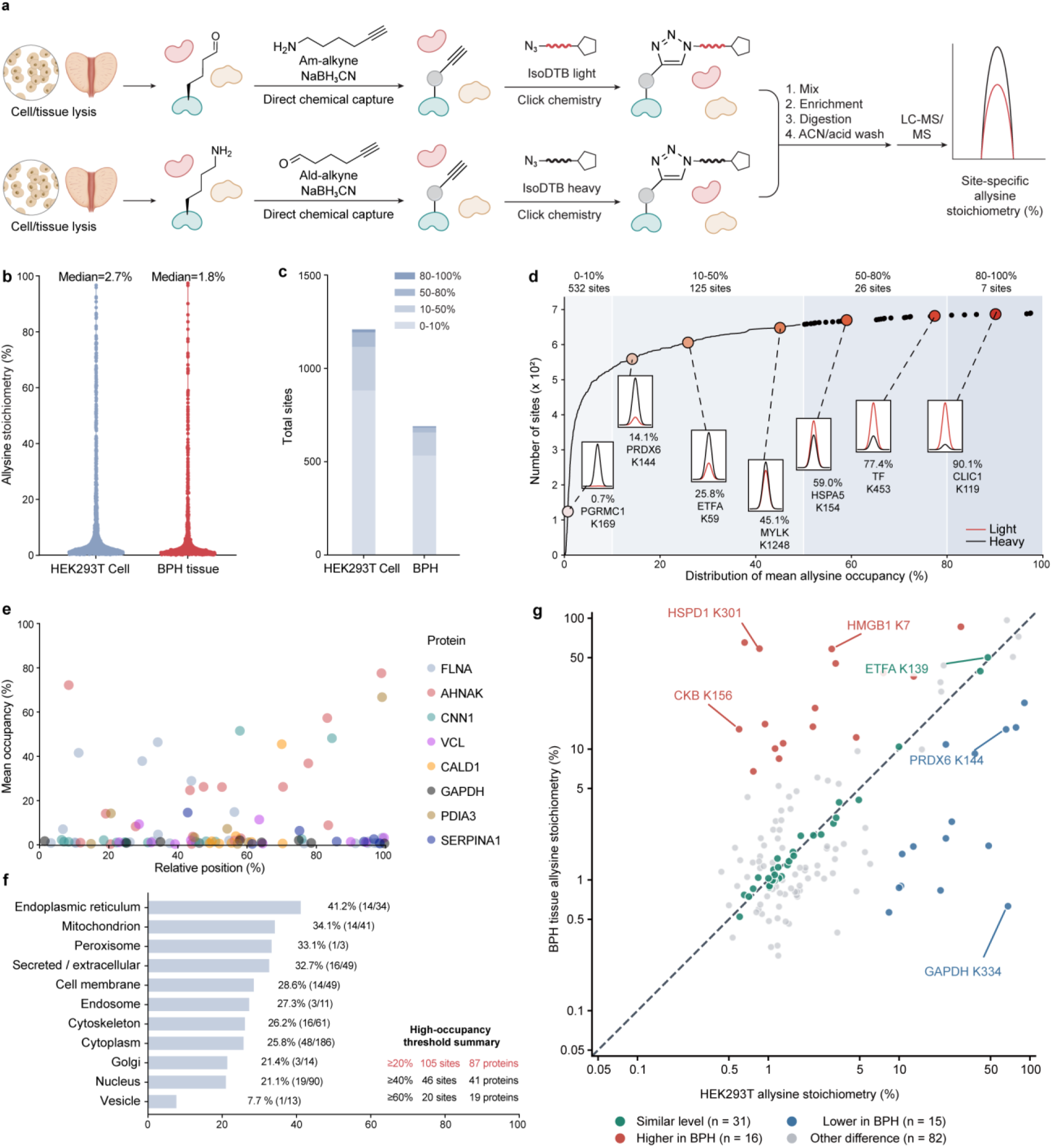
PRISM-Lys enables global site-resolved quantification of allysine stoichiometry. (a) Schematic workflow of PRISM-Lys for site-specific quantification of allysine stoichiometry. Allysine is selectively labeled with Am-alkyne by reductive amination and conjugated to the light IsoDTB tag, whereas reduced Lys is labeled with Ald-alkyne and conjugated to the heavy IsoDTB tag. The light- and heavy-labeled samples are subsequently combined, enriched, digested, washed with ACN/acid, and analyzed by LC–MS/MS. Site-specific allysine stoichiometry is calculated from the heavy-to-light ratio as stoichiometry = 1/[1 + 2^(log₂ H/L)]. (b) Distribution of site-specific allysine occupancies quantified in HEK293T cells and benign prostatic hyperplasia (BPH) tissues. Median occupancies were 2.7% and 1.8%, respectively. (c) Distribution of quantified allysine sites across four stoichiometry ranges (0–10%, 10–50%, 50–80%, and 80–100%) in HEK293T cells and BPH tissues. (d) Cumulative distribution of mean allysine occupancies in BPH tissues. Background shading indicates different stoichiometry intervals, with 532, 125, 26, and 7 sites falling within the 0–10%, 10–50%, 50–80%, and 80–100% ranges, respectively. Insets show representative light/heavy MS1 isotope pairs for sites spanning different stoichiometry levels. (e) Distribution of mean allysine stoichiometry as a function of relative Lys position within representative proteins containing high-stoichiometry sites. Selected high-stoichiometry Lys residues are indicated. (f) Subcellular distribution of proteins containing high-stoichiometry allysine sites. Bars indicate the proportion and corresponding numbers of proteins harboring high-stoichiometry sites within each subcellular compartment. The numbers of sites and proteins exceeding stoichiometry thresholds of 20%, 40%, and 60% are summarized on the right. (g) Scatter plot of mean allysine stoichiometry for 144 shared Lys sites quantified in at least two of three replicates in each sample type. Sites with similar mean stoichiometry are shown in green (BPH/HEK293T ratio of 0.80-1.25 and an absolute difference of ≤5 percentage points; 31 sites). Sites with higher or lower stoichiometry in BPH tissues are shown in red (ratio ≥2 and an increase of ≥5 percentage points; 16 sites) or blue (ratio ≤0.5 and a decrease of ≥5 percentage points; 15 sites), respectively. The remaining 82 sites are shown in gray.

Application of PRISM-Lys to both cultured cells and BPH tissue revealed a predominantly low-stoichiometry but highly heterogeneous landscape of Lys oxidation. Across three biological replicates in HEK293T cells, 1209 allysine sites were reproducibly quantified in at least two replicates, with strong pairwise agreement in log_2_ H/L ratios (Figure S6a). Similarly, measurements across BPH tissue replicates showed good reproducibility and strong pairwise agreement (r = 0.82-0.86) in site-specific log_2_ H/L ratios (Figure S6b). Among the quantified sites, the median allysine stoichiometry was 2.70% in cells and 1.83% in BPH tissue, with most sites exhibiting relatively low modification stoichiometry (Figure 5b). Consistently, 72.9% and 77.1% of quantified sites in cells and BPH tissue, respectively, displayed allysine occupancies below 10%, whereas only a small fraction reached high-stoichiometry states over 80% (Figure 5c). Nevertheless, a distinct subset of sites exhibited intermediate-to-high allysine stoichiometry, indicating that Lys oxidation can accumulate to substantial levels at selected residues. To further visualize the dynamic range of PRISM-Lys measurements, representative sites spanning nearly the entire stoichiometry range were examined by their paired light/heavy MS signals. These included PGRMC1 K169 (0.7%), PRDX6 K144 (14.1%), ETFA K59 (25.8%), MYLK K1248 (45.1%), HSPA5 K154 (59.0%), TF K453 (77.4%), and CLIC1 K119 (90.1%) (Figure 5d). We next examined the distribution of allysine stoichiometry within individual proteins, as multiple Lys residues within the same protein can undergo oxidation to form allysine, potentially exhibiting distinct modification stoichiometries. Mapping allysine stoichiometry across individual proteins revealed striking site-level heterogeneity (Figure 5e). Most Lys residues remained minimally oxidized, whereas a small subset formed pronounced oxidation hotspots. For instance, PDIA3 K500 exhibited 66.7% stoichiometry, while most other PDIA3 sites remained below 10%; similarly, CNN1 K172 and K251 reached 51.5% and 48.2%, respectively, compared with ∼1–3% stoichiometry at most other CNN1 sites. These data demonstrate that Lys oxidation is strongly concentrated at selected residues rather than uniformly distributed across individual proteins.

Furthermore, we asked whether high-stoichiometry allysine events showed preferential association with specific subcellular compartments (Figure 5f). Overall, 87 proteins contained at least one high-stoichiometry allysine site (defined as ≥20% stoichiometry), whereas 41 and 19 proteins contained sites exceeding 40% and 60% stoichiometry, respectively. Proteins localized to the endoplasmic reticulum showed the highest fraction harboring high-stoichiometry allysine sites (41.2%, 14/34), followed by mitochondrial (34.1%, 14/41), peroxisomal (33.3%, 1/3), secreted/extracellular (32.7%, 16/49), and cell-membrane proteins (28.6%, 14/49). Comparison of 144 allysine sites quantified in both HEK293T cells and benign prostatic hyperplasia (BPH) tissues revealed shared and context-dependent patterns of lysine oxidation. Among these sites, 31 exhibited similar stoichiometries, whereas 16 showed higher and 15 showed lower stoichiometries in BPH tissues (Figure 5g). Representative sites with comparable oxidation levels included ETFA K139 (47.62% versus 50.21%), HSPA8 K512 (9.97% versus 10.42%), and MDH2 K241 (1.56% versus 1.53%) in HEK293T cells and BPH tissues, respectively (Figure S7). In contrast, CKB K156 exhibited substantially higher stoichiometry in BPH tissues (14.20%) than in HEK293T cells (0.60%), whereas GAPDH K334 and PRDX6 K144 showed lower stoichiometries in BPH tissues, decreasing from 67.92% to 0.63% and from 65.69% to 14.15%, respectively. Together, PRISM-Lys reveals predominantly low-stoichiometry Lys oxidation with pronounced site- and compartment-specific heterogeneity across cells and tissues, including highly occupied allysine sites on proteins associated with membrane- bound organelles and extracellular compartments.

### PRISM-Lys reveals ferroptosis-responsive remodeling of Lys redox states

Ferroptosis is driven by iron-dependent oxidative reactions,^45^ including Fenton chemistry, which generates reactive radical species and amplifies cellular oxidative stress. A defining feature of this process is the peroxidation of PUFA-containing phospholipids, leading to membrane damage and ultimately cell death.^46, 47^ Given this highly oxidative and iron-rich environment associated with ferroptosis, we asked whether protein Lys oxidation is also dynamically remodeled during this process. To address this question, we applied PRISM-Lys to a pharmacologically controlled ferroptosis-rescue system (Figure 6a). RSL3 (150 nM) treatment induced a pronounced increase in lipid peroxidation in HT1080 Cells, whereas co-treatment with ferrostatin-1 (Fer) or liproxstatin-1 (Lip) effectively suppressed this response, confirming the establishment of a reversible ferroptotic state (Figure 6b). Consistent with this oxidative shift, proteome-wide labeling with the allysine-directed Am-alkyne was enhanced following RSL3 treatment and partially restored upon Lip rescue, whereas labeling of the reduced Lys state with Ald- alkyne showed comparatively modest global changes (Figure 6c, S8a). We therefore used PRISM-Lys to quantify site-specific allysine stoichiometry across control, RSL3-treated, and RSL3/Lip-rescued cells.

**Figure 6.**
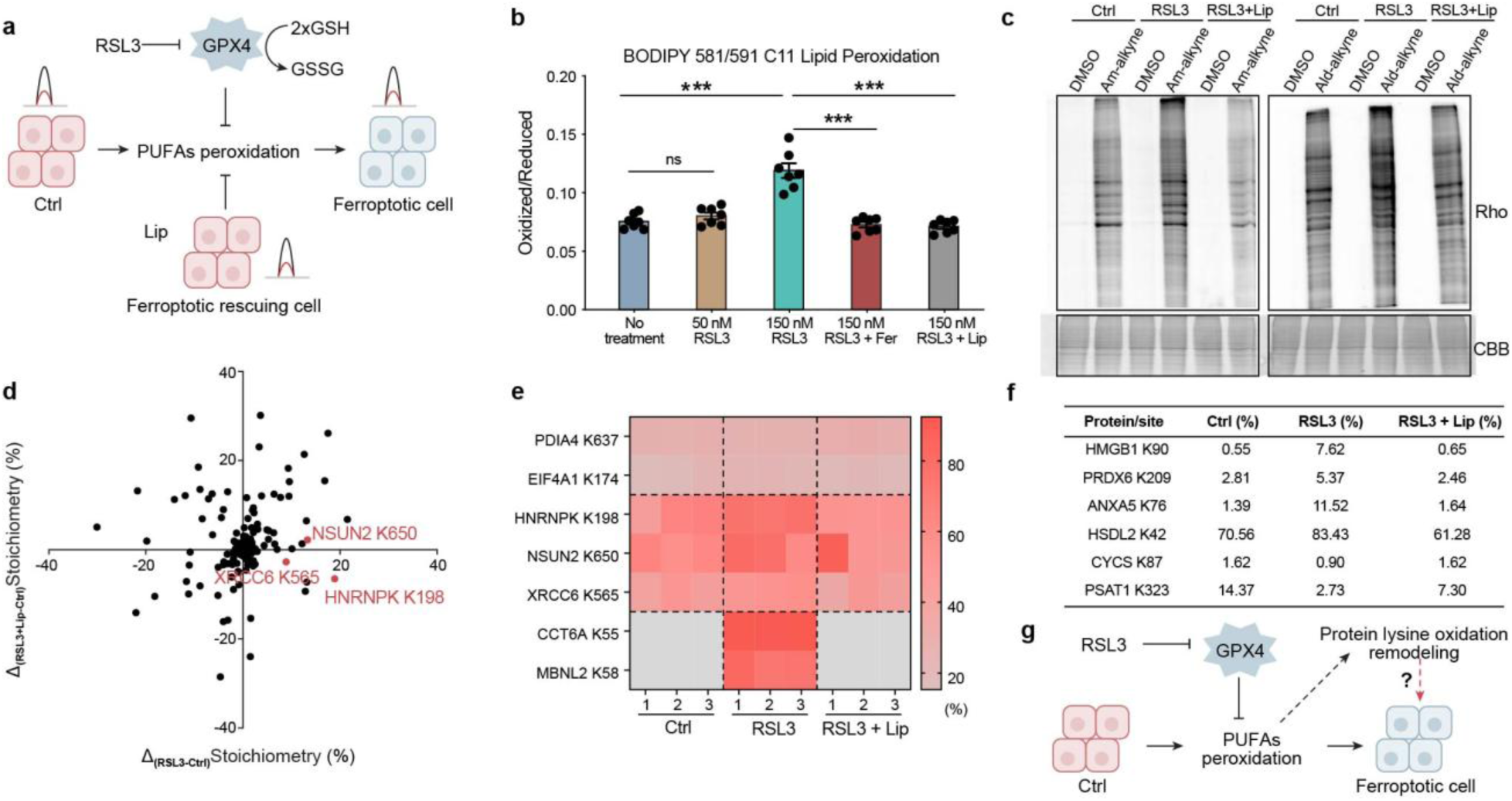
PRISM-Lys reveals site-specific remodeling of Lys oxidation during ferroptosis and its rescue. (a) Schematic illustration of the ferroptosis model. RSL3 inhibits GPX4, promoting PUFA lipid peroxidation and ferroptotic cell death, whereas liproxstatin-1 (Lip) suppresses lipid peroxidation and rescues ferroptosis. (b) C11-BODIPY 581/591 analysis of lipid peroxidation under the indicated treatments. HT1080 Cells were treated with RSL3 at the indicated concentrations, with Fer or Lip used as ferroptosis inhibitors. Data are presented as the oxidized/reduced C11-BODIPY fluorescence ratio. (c) In-gel fluorescence analysis of Am-alkyne labeling in control, RSL3-treated, and RSL3 + Lip-treated HT1080 Cells. DMSO-treated samples served as labeling controls, and Coomassie Brilliant Blue (CBB) staining indicates protein loading. (d) Site-level comparison of changes in allysine stoichiometry induced by RSL3 and following Lip rescue. The x-axis represents the stoichiometry change between RSL3-treated and control cells, whereas the y-axis represents the stoichiometry change between RSL3 + Lip-treated and control cells. Representative ferroptosis-responsive sites, including NSUN2 K650, HNRNPK K198, and XRCC6 K565, are highlighted. (e) Heatmap showing allysine occupancies of representative ferroptosis-responsive sites across three biological replicates of control, RSL3, and RSL3 + Lip conditions. Gray indicates sites not detected (ND). (f) Representative ferroptosis-related proteins exhibiting distinct stoichiometry responses to RSL3 treatment and Lip rescue. Mean site occupancies under control, RSL3, and RSL3 + Lip conditions are shown. (g) Proposed model illustrating ferroptosis-associated remodeling of protein Lys oxidation. RSL3-mediated GPX4 inhibition promotes lipid peroxidation and is accompanied by widespread changes in site-specific Lys oxidation, suggesting a potential protein oxidative footprint associated with ferroptotic stress.

Rather than producing a uniform increase in Lys oxidation, RSL3 induced a heterogeneous remodeling of site-specific allysine stoichiometry. Among sites showing RSL3-induced increases, HNRNPK_K198, NSUN2_K650, and XRCC6_K565 increased from 57.2% to 76.1%, 59.2% to 72.5%, and 43.3% to 52.3%, respectively (Figure S8b), and shifted back toward their basal occupancies following Lip rescue (50.7%, 61.4%, and 40.7%, respectively) (Figure 6d). Conversely, a subset of sites showed decreased stoichiometry upon RSL3 treatment and similarly moved toward control levels after Lip treatment, indicating that Lys oxidation remodeling during ferroptosis is not uniformly directional. Notably, CCT6A_K55 and MBNL2_K58 were undetectable in all control replicates but reproducibly detected across all three RSL3-treated replicates (Figure 6e), and became undetectable again following Lip rescue, defining a distinct class of RSL3-emergent allysine events. Together, these findings indicate that ferroptotic stress reshapes the Lys oxidation landscape in a site-selective manner rather than simply increasing global protein oxidation. Notably, several responsive sites occurred on proteins previously implicated in ferroptosis-associated processes, including HMGB1,^48^ PRDX6,^49^ ANXA5,^50^ HSDL2,^51^ CYCS,^52^ and PSAT1^53^. For example, HMGB1_K90, PRDX6_K209, ANXA5_K76, and HSDL2_K42 showed increased allysine stoichiometry following RSL3 treatment and substantial reversal after Lip rescue (Figure 6f), whereas CYCS_K87 and PSAT1_K323 displayed the opposite pattern, with reduced stoichiometry under RSL3 treatment followed by recovery upon rescue. These bidirectional changes reveal site-specific remodeling of Lys oxidation across functionally diverse proteins during ferroptosis, rather than a uniform increase in oxidation (Figure 6g). Collectively, PRISM-Lys reveals a previously unknown, site-resolved protein oxidation landscape associated with ferroptotic stress and raises the possibility that selected Lys oxidation events may represent not only molecular footprints of ferroptosis but also potential modulators of ferroptotic signaling and cellular response.

## Discussion

Allysine is a chemically distinct product of Lys oxidation and has long been recognized as a marker of protein oxidative damage.^54^ More recently, isotope-enabled chemical proteomics identified hundreds of endogenous allysine sites in cultured cells and tissues, highlighting the prevalence and potential biological relevance of this modification.^24^ Nevertheless, the low abundance and reactive aldehyde functionality of allysine continue to complicate proteome-wide analysis. Here, we combined reductive amination using Am-alkyne probes with a TOP-ABPP workflow to capture allysine. Replacing the DADPS tag with IsoDTB tags provided further evidence supporting the proposed adduct structure. Unlike cysteine residues, Lys side chains are relatively stable under aqueous buffer conditions from an organic chemistry point of view, whereas artifactual allysine formation generally requires metal-catalyzed oxidation with high concentrations of H_2_O_2_. Although CuSO_4_ was used for click chemistry, it was introduced only after Am-alkyne labeling and therefore could not generate the allysine species captured by the probe, supporting the endogenous origin of the identified allysine sites in this study. Importantly, 103 identified allysine sites in our study matched previously reported sites (Figure 2g), providing orthogonal cross-validation of the method’s reliability.

Importantly, identifying an oxidized Lys residue does not reveal the extent of modification.^55^ A site present at 1% stoichiometry and one present at 80% stoichiometry may have very different biological consequences.^56^ Several strategies have therefore been developed to quantify PTM stoichiometry, including phosphatase-assisted phosphorylation measurements,^57^ isotope-encoded chemical completion for Lys acetylation,^58^ isotopically balanced quantification of ubiquitination,^59^ differential redox labeling such as OxICAT,^60^ and protein-level approaches such as STO-MS.^61^ A central analytical challenge in stoichiometry measurements is to enable accurate comparison between modified and unmodified states despite their potentially different chemical and physicochemical properties. Carroll and colleagues developed complementary chemical probes for site-resolved measurement of sulfenylation stoichiometry.^10^ PRISM-Lys employs complementary probes to independently capture oxidized allysine and reduced Lys, converting them into chemically equivalent products distinguished only by isotopic encoding. This convergence, achieved through reductive amination, minimizes state-dependent differences in chromatographic behavior and mass-spectrometric response, enabling direct calculation of site-specific allysine stoichiometry. This strategy could also be integrated with SILAC^62^, TMT^63^, or isotopically labeled probes for quantitative analysis. PRISM-Lys provides a proof of concept for PTM stoichiometry measurement through paired-probe chemical convergence. This strategy may extend to modifications that interconvert functional groups with complementary reactivity, provided that suitable probes convert the modified and unmodified states into a common product. Potential applications include N-terminal pyruvoyl or N-α-ketoacyl modification,^64, 65^ although the corresponding probe chemistry and quantitative performance require validation. Thus, paired-probe chemical convergence offers a design principle for extending site-specific stoichiometry measurements to other chemically tractable PTMs.

It should be noted that the allysine stoichiometry measured here reflects the relative abundance of allysine and unmodified Lys captured by the paired probes, without accounting for other Lys PTMs such as acetylation, methylation, or ubiquitylation. Thus, the calculated values represent the stoichiometric partitioning between allysine and unmodified Lys rather than the absolute fraction of all possible chemical states at a given Lys site. Nevertheless, validation of the paired probe using standard peptides and model proteins provides strong evidence for quantitative accuracy. A more rigorous future validation would involve site-specific incorporation of allysine into proteins through amber-codon suppression^66, 67^, followed by protein-level linearity measurements across defined allysine stoichiometry. Looking ahead, integrating site-specific stoichiometry with protein semisynthesis^68, 69^, could help establish how Lys oxidation at defined sites affects protein structure and function.

To our knowledge, allysine has never before been detected in human prostatic tissues. However, ROS can be produced as a consequence of androgen receptor stimulation, as well as other mechanisms, and thus play a probable role in prostatic diseases^70, 71^. Furthermore, in other contexts, antioxidants may also surprisingly promote prostatic diseases^72, 73^. The stoichiometric quantitative of allysine by PRISM-Lys as we show here thus has the potential to resolve these discrepancies by both elucidating fundamental mechanisms downstream of ROS in prostatic pathologies and for use as a quantitative biomarker. Beyond defining the landscape of allysine stoichiometry across cells and tissues, PRISM-Lys enables site-resolved quantification of allysine changes in response to biological perturbations. We demonstrated this capability in a ferroptosis model, identifying site-specific changing upon RSL3 treatment and rescue by Lip-1. These findings support broader applications of PRISM-Lys to identify redox-sensitive Lys sites and associated pathways across oxidative stress, metabolic perturbations, and disease contexts. In future studies, integrating PRISM-Lys with functional analyses could help uncover previously unrecognized aspects of redox biology.

## Acknowledgements

We would like to thank Emanuel Petricoin and Weidong Zhou from George Mason University for proteomics analysis. We also want to thank Yi Zheng for illustrations (www.yizhengillustration.com) and Zhipeng Meng for kindly providing cell lines. This research was supported by Prostate Cancer Foundation Young Investigator Award (25YOUN18), an Elsa U. Pardee Award (Grant #1290), Women’s Cancer Association (WCA) Award and startup funds from Desai Sethi Urology Institute & Sylvester Comprehensive Cancer Center, University of Miami to Zhipeng A. Wang. Xin Wang was supported by the National Science Foundation grant (2414925).

## Author contributions

Y.F.Z. conceived the study, performed experiments, analyzed data, and wrote the manuscript. L.Z. performed experiments, analyzed data and revised the manuscript. D.M.N.U. provided experimental help and scientific discussion. X.W. contributed to quantitative analyses of peptides and proteins. H.B. and H.N.S. provided BPH tissue samples, scientific advice and edited the manuscript. N.S. provided scientific consultation and revised the manuscript. Z.A.W. contributed to study conception, manuscript writing, figure illustration, and funding.

## Materials availability

The raw MS data have been deposited to the ProteomeXchange Consortium (https://proteomecentral.proteomexchange.org) via the iProX partner repository with the data set identifier PXD086375 (iProx identifier IPX0020316000). All other data are available in the main text or the supplementary materials.

## References

1. C. T. Walsh, S. Garneau-Tsodikova and G. J. Gatto, Jr., Protein posttranslational modifications: the chemistry of proteome diversifications, Angewandte Chemie International Edition, 2005, 44, 7342–7372.

2. Y. L. Deribe, T. Pawson and I. Dikic, Post-translational modifications in signal integration, Nature Structural & Molecular Biology, 2010, 17, 666–672.

3. Y.-F. Zhou, L. Zhang, D. M. N. Udayanga, N. Sharifi and Z. A. Wang, A framework for understanding post-translational modifications of metabolic origin, Trends in Biochemical Sciences, 2026, DOI: 10.1016/j.tibs.2026.08.006.

4. E. A. Kisty, J. A. Falco and E. Weerapana, Redox proteomics combined with proximity labeling enables monitoring of localized cysteine oxidation in cells, Cell Chemical Biology, 2023, 30, 321–336.e6.

5. Q. Wang, Z. Liu, Y. Wang, Y. Liu, Y. Chen, S. Zhang, W. Zeng, D. Li, F. Yang, Z. He, W. Xiao, C. Liu and C. Wang, Quantitative chemoproteomics reveals dopamine’s protective modification of Tau, Nature Chemical Biology, 2025, 21, 1341–1350.

6. J. Gao, R. Liu, K. Huang, Z. Li, X. Sheng, K. Chakraborty, C. Han, D. Zhang, L. Becker and Y. Zhao, Dynamic investigation of hypoxia-induced L-lactylation, Proceedings of the National Academy of Sciences of the United States of America, 2025, 122, e2404899122.

7. S. Lai, Y. Chen, F. Yang, W. Xiao, Y. Liu and C. Wang, Quantitative Site-Specific Chemoproteomic Profiling of Protein Lipoylation, Journal of the American Chemical Society, 2022, 144, 10320–10329.

8. L. Yan, M. Zheng, M. Fan, R. Yao, K. Zou, S. Feng and M. Wu, A Chemoselective Enrichment Strategy for In-Depth Coverage of the Methyllysine Proteome, Angewandte Chemie International Edition, 2024, 63, e202408564.

9. G. Prus, A. Hoegl, B. T. Weinert and C. Choudhary, Analysis and Interpretation of Protein Post-Translational Modification Site Stoichiometry, Trends in Biochemical Sciences, 2019, 44, 943–960.

10. Y. Shi, L. Fu, J. Yang and K. S. Carroll, Wittig reagents for chemoselective sulfenic acid ligation enables global site stoichiometry analysis and redox-controlled mitochondrial targeting, Nature Chemistry, 2021, 13, 1140–1150.

11. H. M. Kagan, M. A. Williams, S. D. Calaman and E. M. Berkowitz, Histone H1 is a substrate for lysyl oxidase and contains endogenous sodium borotritide-reducible residues, Biochemical and Biophysical Research Communications, 1983, 115, 186–192.

12. J. R. Requena, C. C. Chao, R. L. Levine and E. R. Stadtman, Glutamic and aminoadipic semialdehydes are the main carbonyl products of metal-catalyzed oxidation of proteins, Proceedings of the National Academy of Sciences of the United States of America, 2001, 98, 69–74.

13. G. Serra-Bardenys and S. Peiró, Enzymatic lysine oxidation as a posttranslational modification, The FEBS Journal, 2022, 289, 8020–8031.

14. X. Fan, J. Zhang, M. Theves, C. Strauch, I. Nemet, X. Liu, J. Qian, F. J. Giblin and V. M. Monnier, Mechanism of lysine oxidation in human lens crystallins during aging and in diabetes, Journal of Biological Chemistry, 2009, 284, 34618–34627.

15. H. Ma, I. Y. Zhou, Y. I. Chen, N. J. Rotile, I. Ay, E. A. Akam, H. Wang, R. S. Knipe, L. P. Hariri, C. Zhang, M. Drummond, P. Pantazopoulos, B. F. Moon, A. T. Boice, S. E. Zygmont, J. Weigand-Whittier, M. Sojoodi, R. A. Gonzalez-Villalobos, M. K. Hansen, K. K. Tanabe and P. Caravan, Tailored Chemical Reactivity Probes for Systemic Imaging of Aldehydes in Fibroproliferative Diseases, Journal of the American Chemical Society, 2023, 145, 20825–20836.

16. F. Mahjour, V. Dambal, N. Shrestha, V. Singh, V. Noonan, A. Kantarci and P. C. Trackman, Mechanism for oral tumor cell lysyl oxidase like-2 in cancer development: synergy with PDGF-AB, Oncogenesis, 2019, 8, 34.

17. F. Dakterzada, M. Jové, J. L. Cantero, R. Pamplona and G. Piñoll-Ripoll, Plasma and cerebrospinal fluid nonenzymatic protein damage is sustained in Alzheimer’s disease, Redox Biology, 2023, 64, 102772.

18. N. Herranz, N. Dave, A. Millanes-Romero, L. Pascual-Reguant, L. Morey, V. M. Díaz, V. Lórenz-Fonfría, R. Gutierrez-Gallego, C. Jerónimo, A. Iturbide, L. Di Croce, A. García de Herreros and S. Peiró, Lysyl oxidase-like 2 (LOXL2) oxidizes trimethylated lysine 4 in histone H3, The FEBS Journal, 2016, 283, 4263–4273.

19. J. P. Cebrià-Costa, L. Pascual-Reguant, A. Gonzalez-Perez, G. Serra-Bardenys, J. Querol, M. Cosín, G. Verde, R. A. Cigliano, W. Sanseverino, S. Segura-Bayona, A. Iturbide, D. Andreu, P. Nuciforo, C. Bernado-Morales, V. Rodilla, J. Arribas, J. Yelamos, A. G. de Herreros, T. H. Stracker and S. Peiró, LOXL2-mediated H3K4 oxidation reduces chromatin accessibility in triple-negative breast cancer cells, Oncogene, 2020, 39, 79–121.

20. A. Iturbide, L. Pascual-Reguant, L. Fargas, J. P. Cebrià, B. Alsina, A. García de Herreros and S. Peiró, LOXL2 Oxidizes Methylated TAF10 and Controls TFIID-Dependent Genes during Neural Progenitor Differentiation, Molecular Cell, 2015, 58, 755–766.

21. Y. Wei, T. J. Kim, D. H. Peng, D. Duan, D. L. Gibbons, M. Yamauchi, J. R. Jackson, C. J. Le Saux, C. Calhoun, J. Peters, R. Derynck, B. J. Backes and H. A. Chapman, Fibroblast-specific inhibition of TGF-β1 signaling attenuates lung and tumor fibrosis, The Journal of Clinical Investigation, 2017, 127, 3675–3688.

22. A. R. L. Chang, J. A. Stanley, M. C. Robinson, J. W. Sher, Z. Li, Y. A. Chan, A. R. Omdahl, R. Wattiez, A Godzik and S. Matallana-Surget, Protein structure, amino acid composition and sequence determine proteome vulnerability to oxidation-induced damage, The EMBO Journal, 2020, 39, e104523.

23. M. Estévez, P. Padilla, L. Carvalho, L. Martín, A. Carrapiso and J. Delgado, Malondialdehyde interferes with the formation and detection of primary carbonyls in oxidized proteins, Redox Biology, 2019, 26, 101277.

24. Y. C. Sin, N. Hosny, A. B. Batti Angulski, D. H. Kim, J. M. Metzger and Y. Chen, Isotopic Labeling-Enabled Chemical Proteomics Analysis Revealed Structural and Functional Features of Allysine Modifications in Mammalian Cells and Tissues, Analytical Chemistry, 2025, 97, 9944–9952.

25. Q. Li, Z. Zhang, A. Huang, Z. Zhang, X. Yue, P. Yin, X. Chen, M. Ye and H. Qin, Antibody-Free Profiling of Lysine Dimethylation via Chemoselective Oxidation-Coupled Reversible Enrichment, Analytical Chemistry, 2026, 98, 17199–17208.

26. M. L. Matthews, L. He, B. D. Horning, E. J. Olson, B. E. Correia, J. R. Yates, P. E. Dawson and B. F. Cravatt, Chemoproteomic profiling and discovery of protein electrophiles in human cells, Nature Chemistry, 2017, 9, 234–243.

27. Y.-F. Zhou, L. Zhang, Z. L. Niu, X. Wang, A. Storper, R. Hunt, Y. Zhao, N. Sharifi and Z. A. Wang, Chemical Proteomics Identifies Ketogenesis-Mediated Cysteine Modifications Regulating Redox Function, Angewandte Chemie International Edition, 2026, 65, e19830.

28. B. Ma, D. Liu, W. Xiao and C. Wang, Cross-Species Profiling of Itaconate-Mediated Modifications by a Universal Probe, Journal of the American Chemical Society, 2026, DOI: 10.1021/jacs.6c10764.

29. C. A. Ocasio, M. P. Baggelaar, J. Sipthorp, A. Losada de la Lastra, M. Tavares, J. Volarić, C. Soudy, E. M. Storck, J. W. Houghton, S. A. Palma-Duran, J. I. MacRae, G. Tomić, L. Carr, J. Downward, U. S. Eggert and E. W. Tate, A palmitoyl transferase chemical–genetic system to map ZDHHC-specific S-acylation, Nature Biotechnology, 2024, 42, 1548–1558.

30. Y.-F. Zhou, L. Zhang, Z. L. Niu and Z. A. Wang, Targeting the Reactive Proteome: Recent Advances in Activity-Based Protein Profiling and Probe Design, Biomolecules, 2025, 15, 1699.

31. Y. Liu, M. P. Patricelli and B. F. Cravatt, Activity-based protein profiling: the serine hydrolases, Proceedings of the National Academy of Sciences of the United States of America, 1999, 96, 14694–14699.

32. Y.-F. Zhou, S. Yuan, B. Ma, J. Gao and C. Wang, Chemical proteomic profiling of lysine crotonylation using minimalist bioorthogonal probes in mammalian cells, Chemical Science, 2025, 16, 2843–2849.

33. Y. Chen, Y. Cong, B. Quan, T. Lan, X. Chu, Z. Ye, X. Hou and C. Wang, Chemoproteomic profiling of targets of lipid-derived electrophiles by bioorthogonal aminooxy probe, Redox Biology, 2017, 12, 712–718.

34. Y. Chen, Y. Liu, T. Lan, W. Qin, Y. Zhu, K. Qin, J. Gao, H. Wang, X. Hou, N. Chen, J. P. Friedmann Angeli, M. Conrad and C. Wang, Quantitative Profiling of Protein Carbonylations in Ferroptosis by an Aniline-Derived Probe, Journal of the American Chemical Society, 2018, 140, 4712–4720.

35. F. Yang, J. Gao, J. Che, G. Jia and C. Wang, A Dimethyl-Labeling-Based Strategy for Site-Specifically Quantitative Chemical Proteomics, Analytical Chemistry, 2018, 90, 9576–9582.

36. E. Weerapana, A. E. Speers and B. F. Cravatt, Tandem orthogonal proteolysis-activity-based protein profiling (TOP-ABPP)—a general method for mapping sites of probe modification in proteomes, Nature Protocols, 2007, 2, 1414–1425.

37. A. E. Speers and B. F. Cravatt, A Tandem Orthogonal Proteolysis Strategy for High-Content Chemical Proteomics, Journal of the American Chemical Society, 2005, 127, 10018–10019.

38. J. Szychowski, A. Mahdavi, J. J. L. Hodas, J. D. Bagert, J. T. Ngo, P. Landgraf, D. C. Dieterich, E. M. Schuman and D. A. Tirrell, Cleavable Biotin Probes for Labeling of Biomolecules via Azide−Alkyne Cycloaddition, Journal of the American Chemical Society, 2010, 132, 18351–18360.

39. P. R. A. Zanon, L. Lewald and S. M. Hacker, Isotopically Labeled Desthiobiotin Azide (isoDTB) Tags Enable Global Profiling of the Bacterial Cysteinome, Angewandte Chemie International Edition, 2020, 59, 2829–2836.

40. P. R. A. Zanon, F. Yu, P. Z. Musacchio, L. Lewald, M. Zollo, K. Krauskopf, D. Mrdović, P. Raunft, T. E. Maher, M. Cigler, C. J. Chang, K. Lang, F. D. Toste, A. I. Nesvizhskii and S. M. Hacker, Profiling the proteome-wide selectivity of diverse electrophiles, Nature Chemistry, 2025, 17, 1712–1721.

41. N. R. Barthélemy, B. Saef, Y. Li, B. A. Gordon, Y. He, K. Horie, E. Stomrud, G. Salvadó, S. Janelidze, C. Sato, V. Ovod, R. L. Henson, A. M. Fagan, T. L. S. Benzinger, C. Xiong, J. C. Morris, O. Hansson, R. J. Bateman and S. E. Schindler, CSF tau phosphorylation occupancies at T217 and T205 represent improved biomarkers of amyloid and tau pathology in Alzheimer’s disease, Nature Aging, 2023, 3, 391–401.

42. J. V. Olsen, M. Vermeulen, A. Santamaria, C. Kumar, M. L. Miller, L. J. Jensen, F. Gnad, J. Cox, T. S. Jensen, E. A. Nigg, S. Brunak and M. Mann, Quantitative phosphoproteomics reveals widespread full phosphorylation site occupancy during mitosis, Science Signaling, 2010, 3, ra3.

43. B. Emenike, S. Shahin and M. Raj, Bioinspired Synthesis of Allysine for Late-Stage Functionalization of Peptides, Angewandte Chemie International Edition, 2024, 63, e202403215.

44. B. Emenike, P. Czabala, J. Farhi, J. Swaminathan, E. V. Anslyn, J. Spangle and M. Raj, Tertiary Amine Coupling by Oxidation for Selective Labeling of Dimethyl Lysine Post-Translational Modifications, Journal of the American Chemical Society, 2024, 146, 10621–10631.

45. X. Jiang, B. R. Stockwell and M. Conrad, Ferroptosis: mechanisms, biology and role in disease, Nature Reviews Molecular Cell Biology, 2021, 22, 266–282.

46. S. J. Dixon, K. M. Lemberg, M. R. Lamprecht, R. Skouta, E. M. Zaitsev, C. E. Gleason, D. N. Patel, A. J. Bauer, A. M. Cantley, W. S. Yang, B. Morrison III and B. R. Stockwell, Ferroptosis: An Iron-Dependent Form of Nonapoptotic Cell Death, Cell, 2012, 149, 1060–1072.

47. W. S. Yang, R. SriRamaratnam, M. E. Welsch, K. Shimada, R. Skouta, V. S. Viswanathan, J. H. Cheah, P. A. Clemons, A. F. Shamji, C. B. Clish, L. M. Brown, A. W. Girotti, V. W. Cornish, S. L. Schreiber and B. R. Stockwell, Regulation of Ferroptotic Cancer Cell Death by GPX4, Cell, 2014, 156, 317–331.

48. Q. Wen, J. Liu, R. Kang, B. Zhou and D. Tang, The release and activity of HMGB1 in ferroptosis, Biochemical and Biophysical Research Communications, 2019, 510, 278–283.

49. J. Ito, T. Nakamura, T. Toyama, D. Chen, C. Berndt, G. Poschmann, A. S. D. Mourão, S. Doll, M. Suzuki, W. Zhang, J. Zheng, D. Trümbach, N. Yamada, K. Ono, M. Yazaki, Y. Kawai, M. Arisawa, Y. Ohsaki, H. Shirakawa, A. Wahida, B. Proneth, Y. Saito, K. Nakagawa, E. Mishima and M. Conrad, PRDX6 dictates ferroptosis sensitivity by directing cellular selenium utilization, Molecular Cell, 2024, 84, 4629–4644.e9.

50. Y. Gao, H. Zhang, J. Wang, F. Li, X. Li, T. Li, C. Wang, L. Li, R. Peng, L. Liu, W. Cui, S. Zhang and J. Zhang, Annexin A5 ameliorates traumatic brain injury-induced neuroinflammation and neuronal ferroptosis by modulating the NF-ĸB/HMGB1 and Nrf2/HO-1 pathways, International Immunopharmacology, 2023, 114, 109619.

51. S. Ma, Y. Ma, F. Qi, J. Lei, F. Chen, W. Sun, D. Wang, S. Zhou, Z. Liu, Z. Lu and D. Zhang, HSDL2 knockdown promotes the progression of cholangiocarcinoma by inhibiting ferroptosis through the P53/SLC7A11 axis, World Journal of Surgical Oncology, 2023, 21, 293.

52. X. Song, Z. Zhou, J. Liu, J. Li, C. Yu, H. J. Zeh, D. J. Klionsky, B. R. Stockwell, J. Wang, R. Kang, G. Kroemer and D. Tang, Cytosolic cytochrome c represses ferroptosis, Cell Metabolism, 2025, 37, 1326–1343.e10.

53. P. Zheng, Z. Hu, Y. Shen, L. Gu, Y. Ouyang, Y. Duan, G. Ji, B. Dong, Y. Lin, T. Wen, Q. Tian, Y. Hou, Q. Zhou, X. Sun, X. Chen, K. L. Wang, S. Luo, S. Wu, Y. Sun, M. Li, L. Xiao, Q. Wu, Y. Meng, G. Liu, Z. Wang, X. Bai, S. Duan, Y. Ding, Y. Bi, Y. Wang, G. Li, X. Liu, Z. Lu, X. Wu, Z. Tang and D. Xu, PSAT1 impairs ferroptosis and reduces immunotherapy efficacy via GPX4 hydroxylation, Nature Chemical Biology, 2025, 21, 1420–1432.

54. R. Kehm, T. Baldensperger, J. Raupbach and A. Höhn, Protein oxidation - Formation mechanisms, detection and relevance as biomarkers in human diseases, Redox Biology, 2021, 42, 101901.

55. K. S. Carroll and J. Yang, Defining and refining the cysteine redoxome through sulfur chemical biology, Nature Chemical Biology, 2026, 22, 525–539.

56. G. Prus, S. Satpathy, B. T. Weinert, T. Narita and C. Choudhary, Global, site-resolved analysis of ubiquitylation occupancy and turnover rate reveals systems properties, Cell, 2024, 187, 2875–2892.e21.

57. R. Wu, W. Haas, N. Dephoure, E. L. Huttlin, B. Zhai, M. E. Sowa and S. P. Gygi, A large-scale method to measure absolute protein phosphorylation stoichiometries, Nature Methods, 2011, 8, 677–683.

58. J. Baeza, J. A. Dowell, M. J. Smallegan, J. Fan, D. Amador-Noguez, Z. Khan and J. M. Denu, Stoichiometry of Site-specific Lysine Acetylation in an Entire Proteome, Journal of Biological Chemistry, 2014, 289, 21326–21338.

59. Y. Li, J. Evers, A. Luo, L. Erber, Z. Postler and Y. Chen, A Quantitative Chemical Proteomics Approach for Site-specific Stoichiometry Analysis of Ubiquitination, Angewandte Chemie International Edition, 2019, 58, 537–541.

60. L. I. Leichert, F. Gehrke, H. V. Gudiseva, T. Blackwell, M. Ilbert, A. K. Walker, J. R. Strahler, P. C. Andrews and U. Jakob, Quantifying changes in the thiol redox proteome upon oxidative stress in vivo, Proceedings of the National Academy of Sciences of the United States of America, 2008, 105, 8197–8202.

61. Y. Chen, B. Quan, Y. Li, Y. Liu, W. Qin and C. Wang, Quantitative profiling of PTM stoichiometry by resolvable mass tags, RSC Chemical Biology, 2022, 3, 1320–1324.

62. S.-E. Ong, B. Blagoev, I. Kratchmarova, D. B. Kristensen, H. Steen, A. Pandey and M. Mann, Stable Isotope Labeling by Amino Acids in Cell Culture, SILAC, as a Simple and Accurate Approach to Expression Proteomics, Molecular & Cellular Proteomics, 2002, 1, 376–386.

63. A. Thompson, J. Schäfer, K. Kuhn, S. Kienle, J. Schwarz, G. Schmidt, T. Neumann and C. Hamon, Tandem Mass Tags: A Novel Quantification Strategy for Comparative Analysis of Complex Protein Mixtures by MS/MS, Analytical Chemistry, 2003, 75, 1895–1904.

64. C. Tian, K. Liu, R. Sun, L. Fu and J. Yang, Chemoproteomics Reveals Unexpected Lysine/Arginine-Specific Cleavage of Peptide Chains as a Potential Protein Degradation Machinery, Analytical Chemistry, 2018, 90, 794–800.

65. V. L. Davidson, Protein-Derived Cofactors Revisited: Empowering Amino Acid Residues with New Functions, Biochemistry, 2018, 57, 3115–3125.

66. Z. A. Wang, Y. Zeng, Y. Kurra, X. Wang, J. M. Tharp, E. C. Vatansever, W. W. Hsu, S. Dai, X. Fang and W. R. Liu, A Genetically Encoded Allysine for the Synthesis of Proteins with Site-Specific Lysine Dimethylation, Angewandte Chemie International Edition, 2017, 56, 212–216.

67. Z. A. Wang, Y. Kurra, X. Wang, Y. Zeng, Y.-J. Lee, V. Sharma, H. Lin, S. Y. Dai and W. R. Liu, A Versatile Approach for Site-Specific Lysine Acylation in Proteins, Angewandte Chemie International Edition, 2017, 56, 1643–1647.

68. Z. A. Wang, J. W. Markert, S. D. Whedon, M. Yapa Abeywardana, K. Lee, H. Jiang, C. Suarez, H. Lin, L. Farnung and P. A. Cole, Structural Basis of Sirtuin 6-Catalyzed Nucleosome Deacetylation, Journal of the American Chemical Society, 2023, 145, 6811–6822.

69. Z. A. Wang, S. D. Whedon, M. Wu, S. Wang, E. A. Brown, A. Anmangandla, L. Regan, K. Lee, J. Du, J. Y. Hong, L. Fairall, T. Kay, H. Lin, Y. Zhao, J. W. R. Schwabe and P. A. Cole, Histone H2B Deacylation Selectivity: Exploring Chromatin’s Dark Matter with an Engineered Sortase, Journal of the American Chemical Society, 2022, 144, 3360–3364.

70. M. O. Ripple, W. F. Henry, R. P. Rago and G. Wilding, Prooxidant-antioxidant shift induced by androgen treatment of human prostate carcinoma cells, Journal of the National Cancer Institute, 1997, 89, 40–48.

71. N. Sharifi, E. M. Hurt, S. B. Thomas and W. L. Farrar, Effects of Manganese Superoxide Dismutase Silencing on Androgen Receptor Function and Gene Regulation: Implications for Castration-Resistant Prostate Cancer, Clinical Cancer Research, 2008, 14, 6073–6080.

72. E. A. Klein, I. M. Thompson, C. M. Tangen, J. J. Crowley, M. S. Lucia, P. J. Goodman, L. M. Minasian, L. G. Ford, H. L. Parnes, J. M. Gaziano, D. D. Karp, M. M. Lieber, P. J. Walther, L. Klotz, J. K. Parsons, J. L. Chin, A. K. Darke, S. M. Lippman, G. E. Goodman, F. L. Meyskens and L. H. Baker, Vitamin E and the Risk of Prostate Cancer: The Selenium and Vitamin E Cancer Prevention Trial (SELECT), JAMA (Journal of the American Medical Association), 2011, 306, 1549–1556.

73. M. M. Swamynathan, S. Kuang, K. E. Watrud, M. R. Doherty, C. Gineste, G. Mathew, G. Q. Gong, H. Cox, E. Cheng, D. Reiss, J. Kendall, D. Ghosh, C. R. Reczek, X. Zhao, T. Herzka, S. Špokaitė, A. N. Dessus, S. T. Kim, O. Klingbeil, J. Liu, D. G. Nowak, H. Alsudani, T.-L. Wee, Y. Park, F. Minicozzi, K. Rivera, A. S. Almeida, K. Chang, R. P. Chakrabarty, J. E. Wilkinson, P. A. Gimotty, S. D. Diermeier, M. Egeblad, C. R. Vakoc, J. W. Locasale, N. S. Chandel, T. Janowitz, J. B. Hicks, M. Wigler, D. J. Pappin, R. L. Williams, P. Cifani, D. A. Tuveson, J. Laporte and L. C. Trotman, Dietary pro-oxidant therapy by a vitamin K precursor targets PI 3-kinase VPS34 function, Science, 2024, 386, eadk9167.

