## Supplementary Information for "Paired-probe chemoproteomic strategy enables global site-resolved quantification of lysine redox states"

#### Electronic Supplementary Information (ESI)

#### Table of contents

1. Supplementary tables ........................................................ S3
2. Supplementary figures ........................................................ S4
3. Organic synthesis materials and methods ........................................................ S13
4. Quantification of model peptides and protein ........................................................ S17
5. Biological experiments materials and methods ........................................................ S19
6. Chemical proteomics study ........................................................ S20
7. Characterization of the compounds ........................................................ S25
8. References ........................................................ S39

#### 1. Supplementary tables S1-S6

Table S1: open search results for probe 1 to 4 (6 sheets)

Table S2: Am-alkyne sites in HEK293T cell and BPH tissue (7 sheets)

Table S3: open search of Ald-alkyne and its sites (6 sheets)

Table S4: quantification datasets of model peptides and protein (4 sheets)

Table S5: stiochiometry in HEK293T cell and BPH tissues (11 sheets)

Table S6: stoichiometry in HT1080 cells ctrl or RSL3 treatment or rescue (11 sheets)

#### 2. Supplementary figures

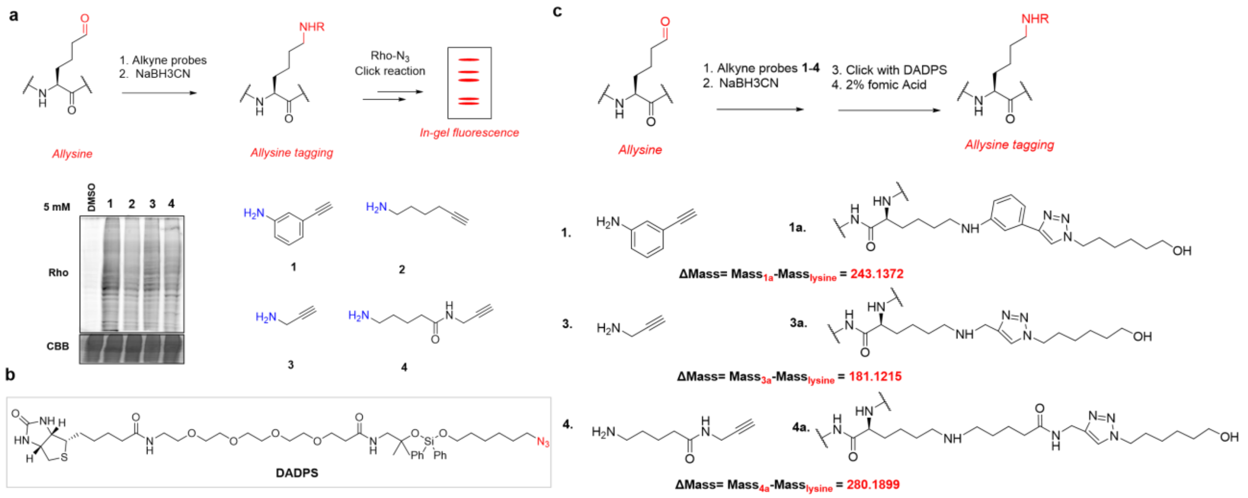


**Figure S1.** (a) In-gel fluorescence analysis of alkyne-functionalized amine probes in HEK293T cell lysates. Coomassie Brilliant Blue (CBB). (b) Structure of DADPS tag. (c) Theoretically predicted probe-derived adducts and their corresponding mass calculations.

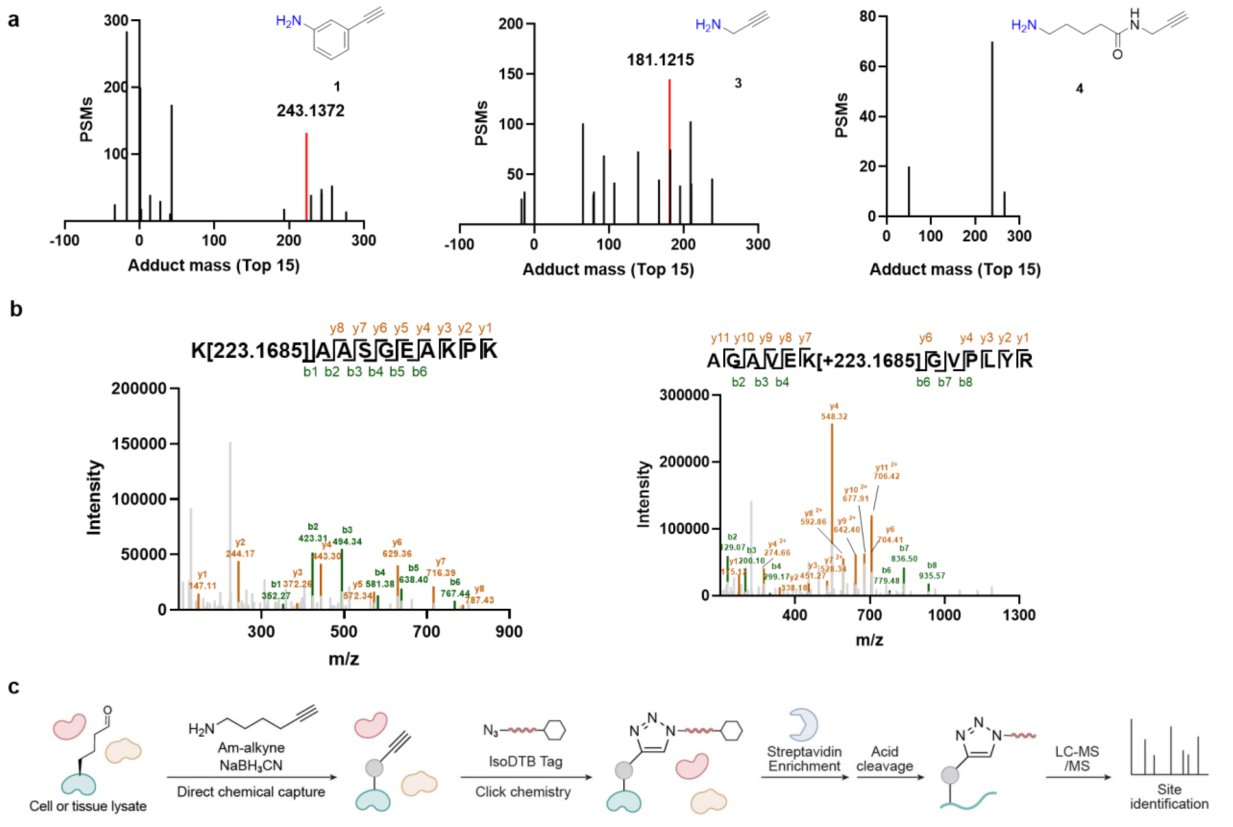


**Figure S2.** (a) Open-search results for compounds **1**, **3**, and **4**, showing the top 15 most abundant mass shifts. (b) Representative MS/MS fragmentation spectra of H1.2_K110Allysine (K[223.1685]AASGEAKPK) and ENO1_K126Allysine (AGAVEK[223.1685]GVPLYR). (c) Validation of allysine identification using IsoDTB tags.

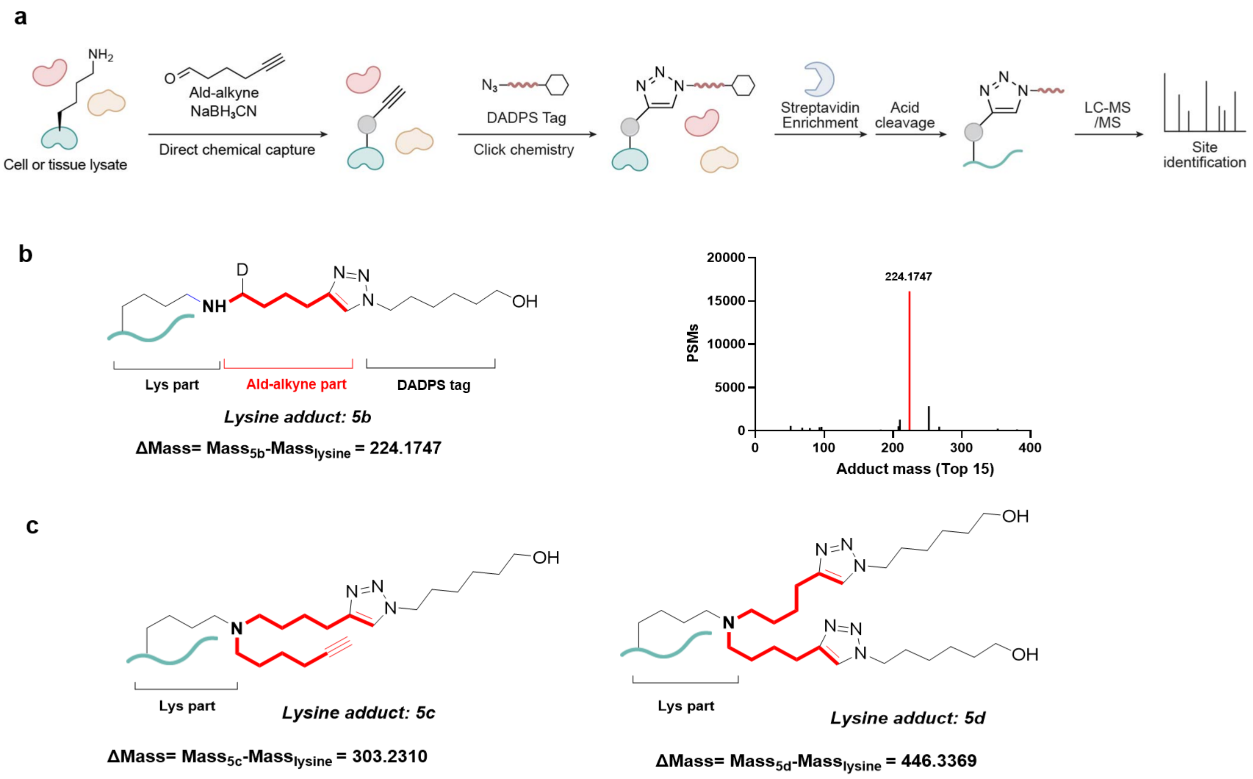


**Figure S3.** (a) TOP-ABPP analysis of the Lys reactivity of **5** (Ald-alkyne). (b) open search yielded a mass shift of 224.1747 Da Lys adducts with $NaBD_3CN$. (c) Proposed adducts bearing two Ald-alkyne-derived substituents at 303.2310 and 446.3369 Da, respectively.

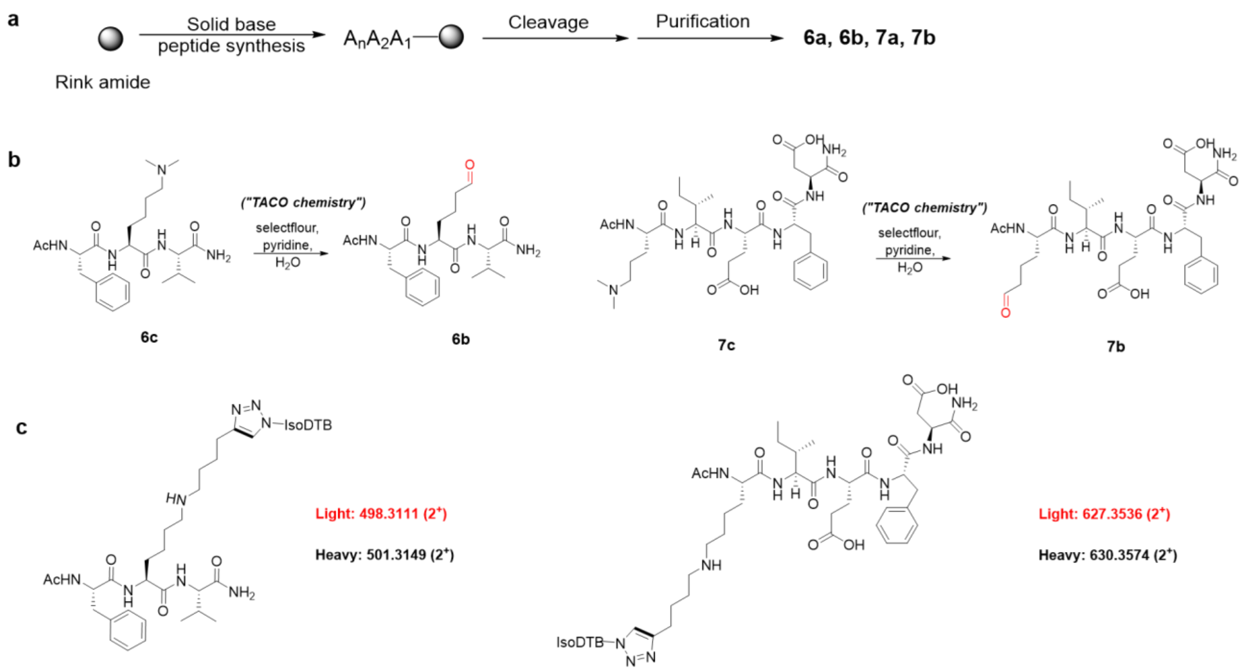


**Figure S4.** (a) SPPS workflow for the synthesis of peptides **6a**, **6b**, **7a**, and **7b**. (b) Synthesis of the corresponding allysine-containing peptides **6b** and **7b** from **6c** and **7c**, respectively. (c) Corresponding doubly charged (2+) light and heavy adducts derived from model peptides.

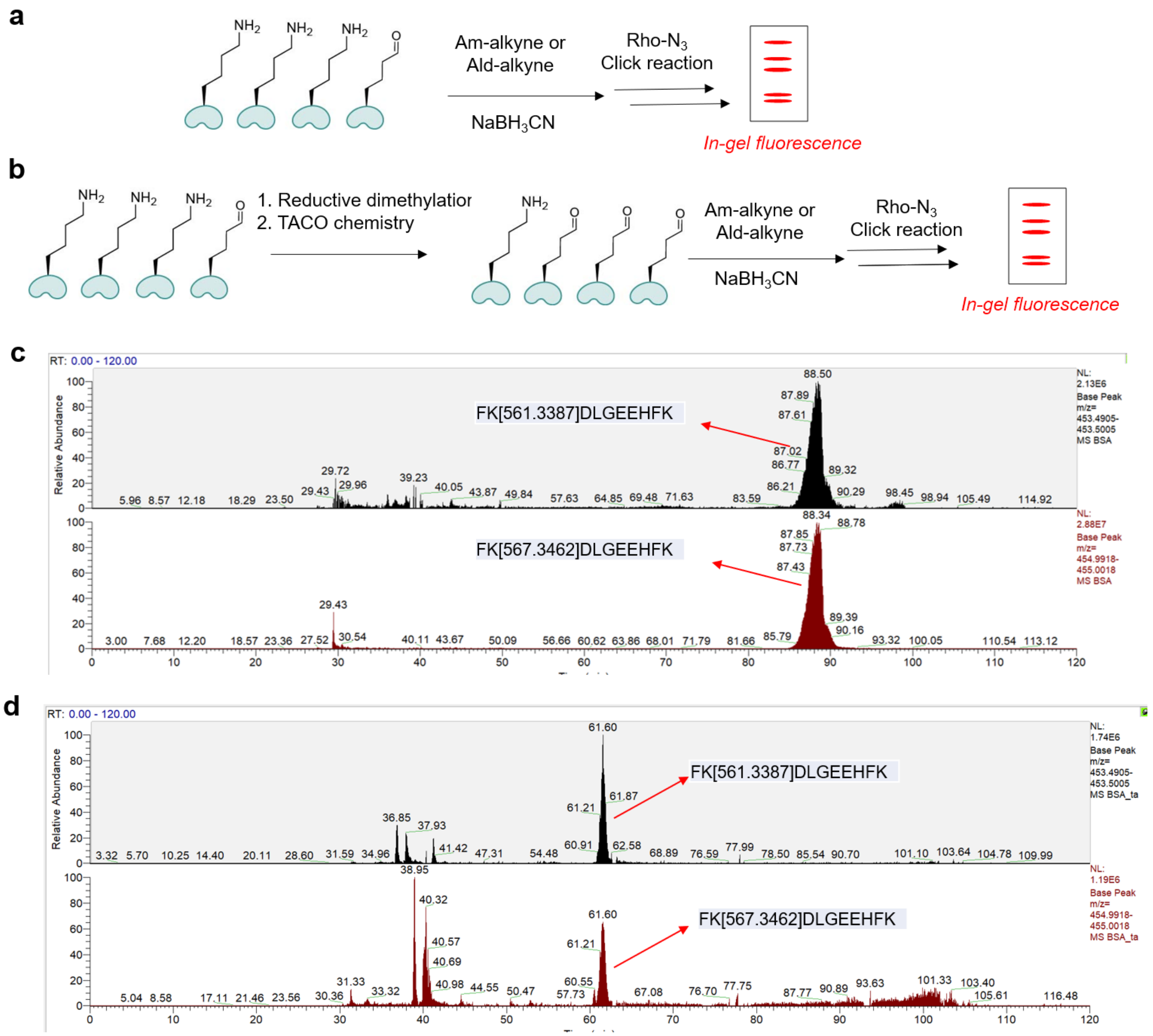


**Figure S5.** (a) BSA and lysozyme were used as model proteins for in-gel fluorescence labeling with Am-alkyne and Ald-alkyne. (b) BSA and lysozyme were first subjected to reductive dimethylation and TACO chemistry, followed by labeling with Am-alkyne and Ald-alkyne and analysis by in-gel fluorescence. (c, d) Representative MS1 analysis of the BSA-derived allysine-containing peptide FK(allysine)DLGEEHFK with or without TACO treatment.

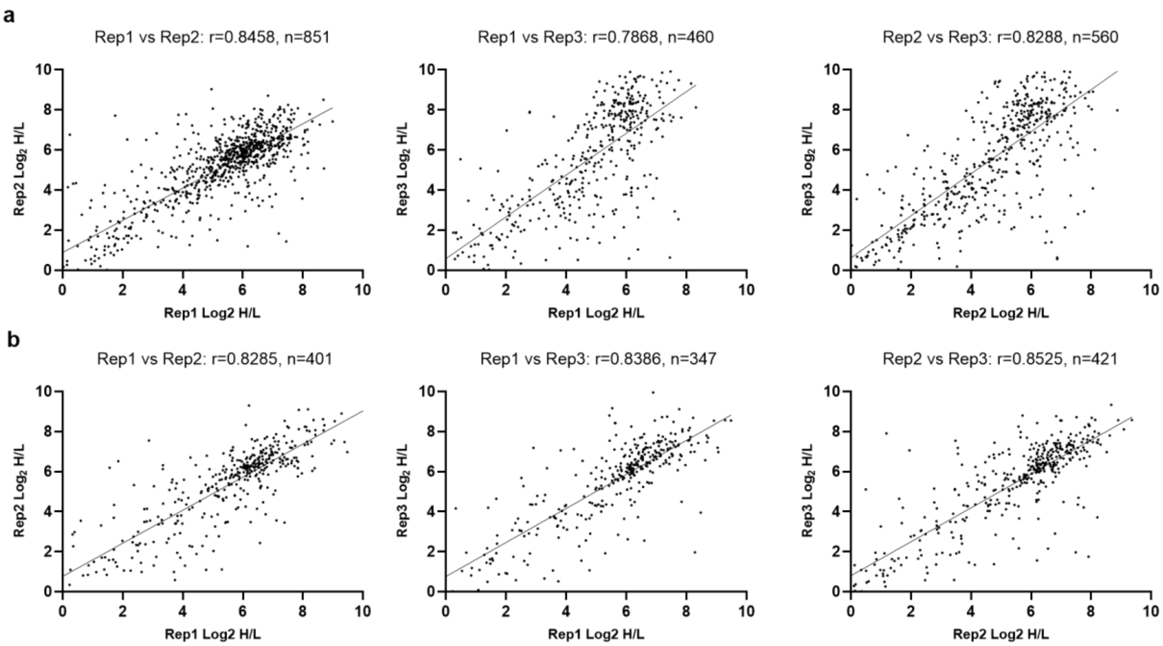


**Figure S6.** Pairwise reproducibility of site-level $Log_2$ H/L quantification in cell and BPH tissue samples. Pearson correlation analysis of $Log_2$ H/L ratios between three biological replicates in cell (a) and BPH tissue (b) samples. The Pearson correlation coefficient (r) and number of commonly quantified sites (n) are indicated for each comparison.

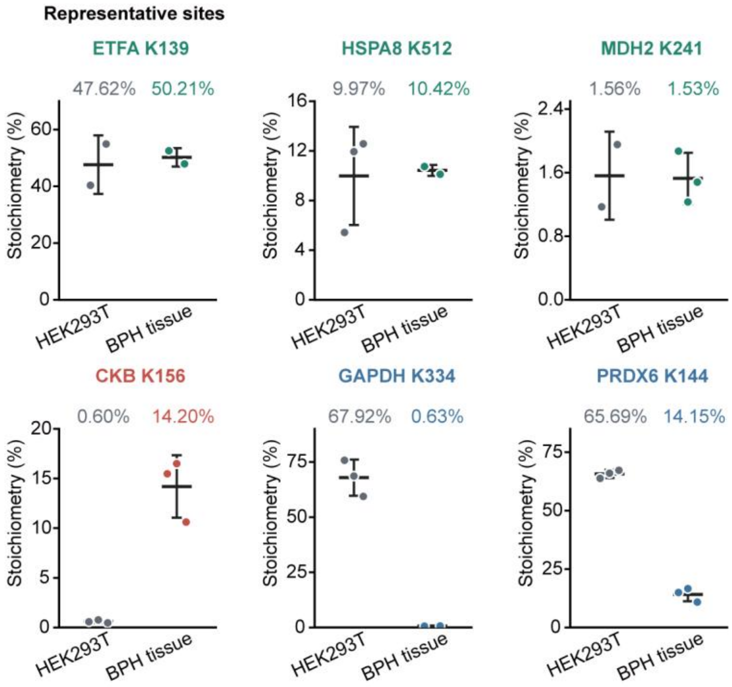


**Figure S7. Comparison of site-specific allysine stoichiometry between HEK293T cells and BPH tissues.** Replicate-level stoichiometries of representative sites with similar levels (ETFA K139, HSPA8 K512, and MDH2 K241), higher levels (CKB K156), or lower levels (GAPDH K334 and PRDX6 K144) in BPH tissues. Points represent individual quantified replicates (n = 2–3 per group), and horizontal lines and error bars indicate mean ± SD. Values above each group indicate mean stoichiometry. Each subplot uses an independent linear y-axis. Missing values were excluded without imputation.

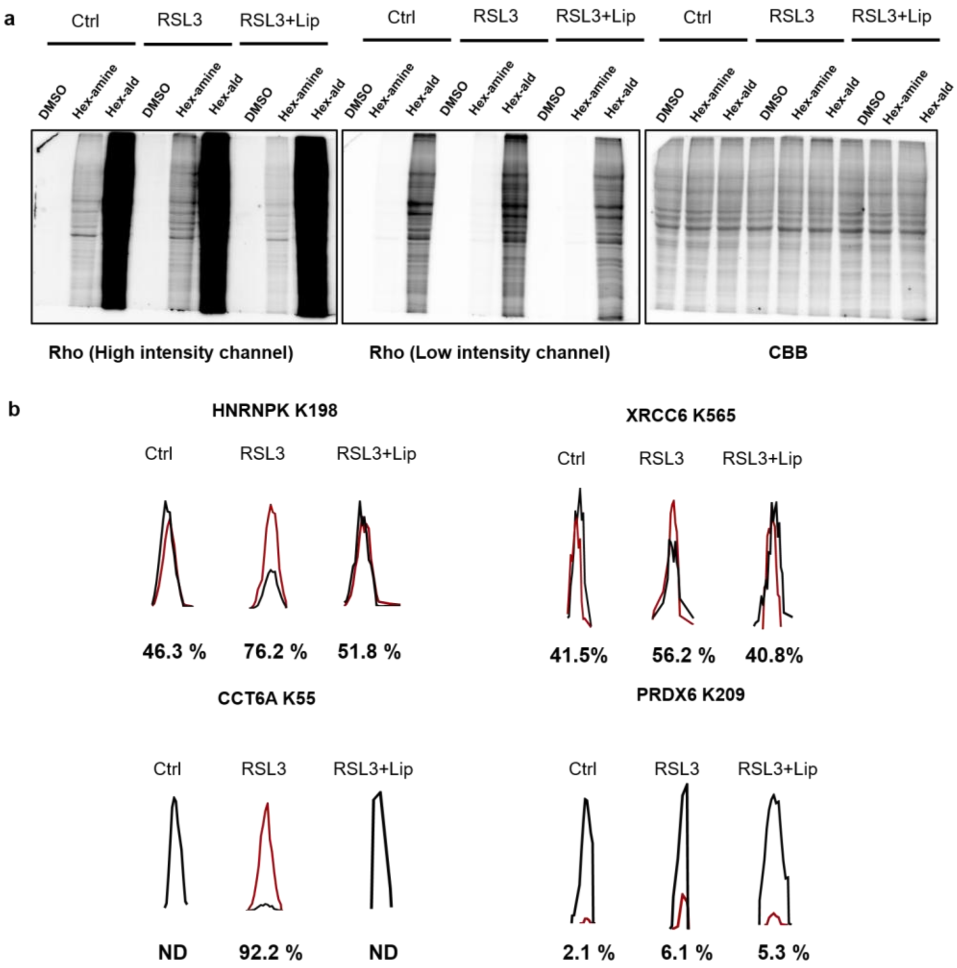


**Figure S8.** (a) In-gel fluorescence analysis of proteins labeled with Am-alkyne and Ald-alkyne in control (Ctrl), RSL3-treated, and RSL3 plus liproxstatin-1-treated (RSL3+Lip) samples. DMSO served as the labeling control. Rhodamine (Rho) fluorescence is shown at high and low intensity settings. CBB staining shows protein loading. (b) Representative extracted ion chromatograms of paired isotope-labeled peptides corresponding to HNRNPK K198, XRCC6 K565, CCT6A K55, and PRDX6 K209. The percentages below the chromatograms indicate allysine stoichiometry at each site. ND, allysine stoichiometry could not be determined.

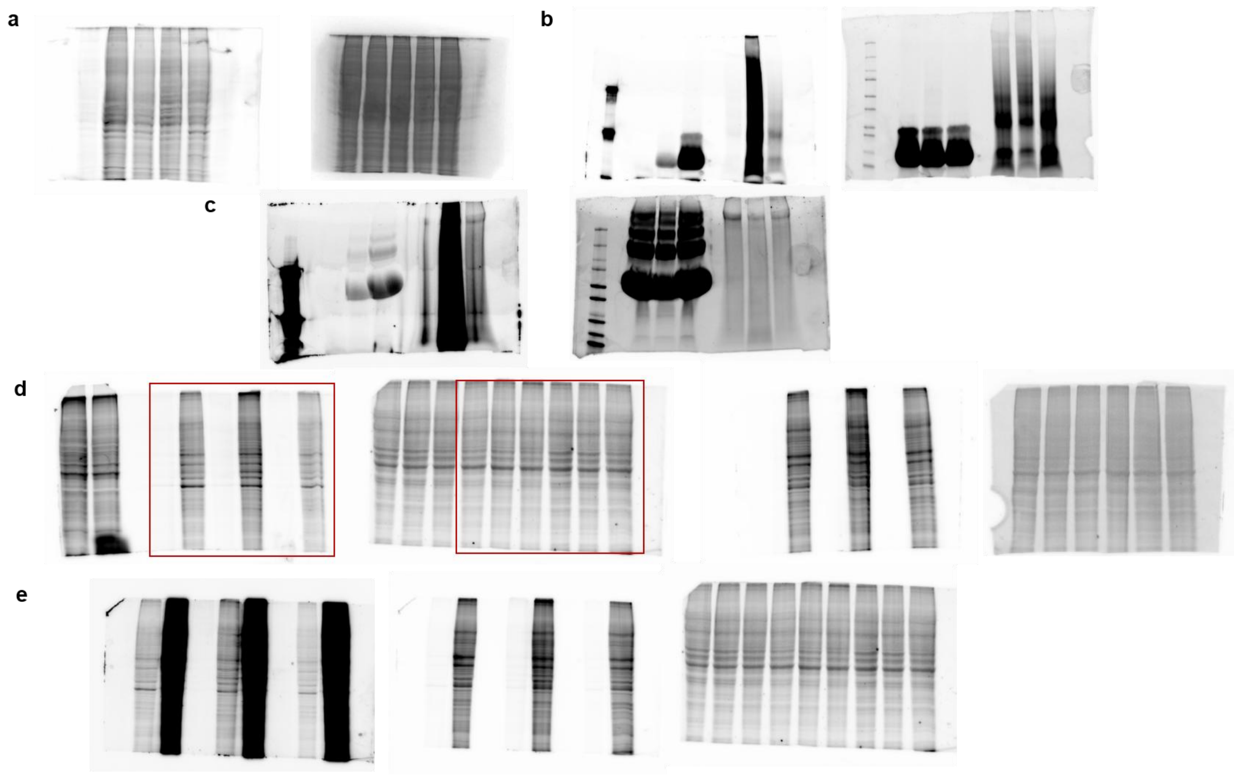


**Figure S9. Uncropped blot for figure S1a (a), figure 4e (b), figure 4f (c), figure 6c (d), figure S7a (e) in this manuscript.**

#### 3. Organic synthesis materials and methods

##### 3.1 General Information

All reactions were carried out under an argon atmosphere with dry solvents under anhydrous conditions, unless otherwise noted. Reagents were purchased at the highest commercial quality and used without further purification, unless otherwise stated. Solvents for chromatography were used as supplied by Merck chemicals. Reactions were monitored by thin layer chromatography (TLC) carried out on 0.2 mm Merck gel plates (60F-254) using UV light as visualizing agent and aqueous ammonium cerium nitrate/ammonium molybdate as developing agent. Merck silica gel (60, particle size 0.040-0.063mm) was used for flash column chromatography. NMR spectra were recorded on Bruker Advance 500 or 800 MHz. The spectra were calibrated by using residual undeuterated solvents (for $^{1}$H NMR) and deuterated solvents (for $^{13}$C NMR) as internal references: chloroform ($\delta_H$ = 7.26 ppm) and $CDCl_3$ ($\delta_C$ = 77.16 ppm) and DMSO-$d_6$ ($\delta$H = 2.50 ppm; $\delta$C = 39.52 ppm). The following abbreviations are used to designate multiplicities: s = singlet, d = doublet, t = triplet, q = quartet, m = multiplet, quint = quintet, br = broad. High-resolution mass spectra (HR-MS) were recorded on Bruker impact II Q-TOF MS. Probes **1**, **2** (Am-alkyne) and **3** are commercial sources and probe **2** (Am-alkyne) is confirmed by NMR (Please see characterization of the compounds part). Probes **4** and **5** (Ald-alkyne) and model peptides **6a**, **6b**, **6c** and **7a**, **7b**, **7c** are synthesized in lab and confirmed by NMR and HR-MS (Please see characterization of the compounds part).

##### 3.2 Synthesis of Ald-alkyne (5) and probe 4

**Synthesis of Ald-alkyne (5)**

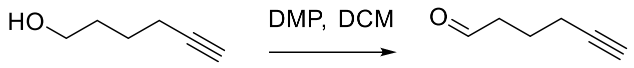


**5-Hexyn-1-ol** **Ald-alkyne (5)**

The compound **1** (5-hexyne-1-o1, 0.1 M, 9.8 g), were dissolved in 100 mL of dichloromethane (DCM) and then added Dess-Martin periodinane (DMP, 0.1M, 42.4 g) stirred at room temperature for 3 hours. The crude product was added $NaHCO_3$ to convert free acetic acid (from DMP) to sodium acetate and obtained Ald-alkyne by redistillation as a white oil (5 g, 52 % yield). $^{1}$H NMR (500 MHz, Chloroform-*d*) $\delta$ 9.80 (s, 1H), 2.61 (t, $J$ = 7.2 Hz, 2H), 2.37 - 2.17 (m, 2H), 2.10 - 1.97 (m, 1H), 1.86 (q, $J$ = 7.1 Hz, 2H). $^{13}$C NMR (126 MHz, $CDCl_3$) δ 201.7, 83.2, 69.4, 42.5, 20.8, 17.7. HR-ESI (m/z): $[M + H]^+$ calcd for $C_6H_9O$ 97.0646, found 97.0653.

**Synthesis of probe 4**

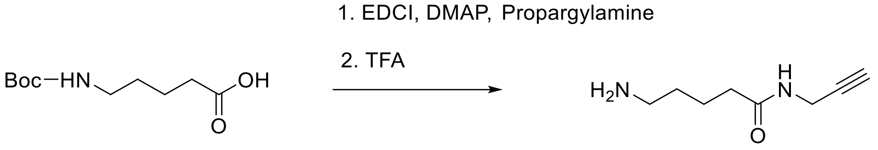


**Boc-5-Ava-OH**

**4**

Compound boc-5-Ava-OH (2.17 g, 10 mmol), EDCI (1.97 g, 10 mmol), DMAP (2.44 g, 20 mmol) was added propargylamine (0.55 g, 10 mmol) in DCM (10 mL) at room temperature. The reaction mixture was stirred for 2 hours. Then the crude mixture was quenched with water and extracted with ethyl acetate. The combined organic extracts were concentrated under reduced pressure to afford crude intermediates. The intermediates were treated with 3 mL of 50% TFA in DCM and stirred for 3 h. The mixture was then concentrated under reduced pressure, and the residue was suspended in saturated aqueous $NaHCO_3$ and extracted with ethyl acetate three times. The combined organic extracts were concentrated under reduced pressure to afford the crude product. The crude product was purified by flash column chromatography (DCM: acetone = 5:1, v/v) and further concentrated under reduced pressure to yield probe 4 as a white oil (1.25 g, 78 % yield). $^1$H NMR (800 MHz, Methanol-$d_4$) $\delta$ 3.95 (d, $J$ = 3.3 Hz, 2H), 2.96 (t, $J$ = 6.4 Hz, 2H), 2.54 (d, $J$ = 3.0 Hz, 1H), 2.29 (d, $J$ = 6.4 Hz, 2H), 1.83 - 1.57 (m, 4H). $^{13}$C NMR (201 MHz, MeOD) $\delta$ 173.8, 79.2, 70.8, 39.0, 34.4, 28.1, 26.6, 22.0. HR-ESI (m/z): [M + H]$^+$ calcd for $C_8H_{15}N_2O$ 155.1179, found 155.1281.

##### 3.3 Solid-phase peptide synthesis

##### 3.3.1 Automated solid-phase peptide synthesis

Automated Fmoc-based solid-phase peptide synthesis was performed on a CEM Liberty Blue 2.0 synthesizer at 0.1 mmol scale using DMF as solvent. The resin was pre-swelled in DMF. Each cycle involved Fmoc deprotection with 10% piperidine in DMF (4 mL) at 90 °C for 1 min, followed by DMF washes (4 × 4 mL). Coupling was carried out with 5.0 equiv. Fmoc-amino acid, 10.0 equiv. DIC, and 5.0 equiv. Oxyma under microwave heating at 90 °C for 2 min. The dimethyl Lyswas used to synthesize dimethyl containing Lys. The process was repeated for each residue in the sequence.

##### 3.3.2 Acid Cleavage Procedures

Each peptide was cleaved from resin using a freshly prepared cocktail of TFA/TIS/$H_2O$/DODt (92.5:2.5:2.5:2.5, v/v), with 10 mL prepared per cleavage. Approximately 5 mL was added to the peptide-bound resin and shaken vigorously at room temperature for 2.5 hours. The mixture was collected, and the resin was rinsed with an additional 3 mL of the same cocktail. The combined flow-through (FT) was partially evaporated by airflow, then diluted to 50 mL with cold (-20 °C) diethyl ether. After vortexing and centrifugation (20,000 g, 15 min, 4 °C), the peptide pellet was washed once more with cold ether and centrifuged again. The resulting pellet was air-dried under a KimWipe in a fume hood for ~2 hours and stored at -20 °C.

**3.3.3 Tertiary amine coupling by Oxidation (TACO) chemistry**

To 10 mg of unprotected dimethyllysine peptide dissolved in 1000 μL of 10 mM sodium phosphate buffer (NaP, pH 7.0), was added pyridine (5 eq) and selectfluor (2 eq). The reaction mixture was stirred for 1 h and subsequently injected into the HPLC for isolation the aldehyde peptide product.

**3.3.4 Confirmation of final products**

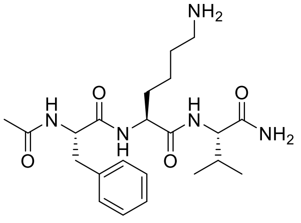

**6a**

**Compound 6a:** $^1$H NMR (800 MHz, DMSO-d$_6$) $\delta$ 8.18 (d, $J$ = 7.9 Hz, 1H), 8.06 (dd, $J$ = 8.1, 4.7 Hz, 1H), 7.69 (d, $J$ = 20.7 Hz, 3H), 7.57 – 7.53 (m, 1H), 7.36 – 7.33 (m, 1H), 7.19 (q, $J$ = 4.9, 4.3 Hz, 4H), 7.11 (tt, $J$ = 6.2, 2.4 Hz, 1H), 4.43 (ddd, $J$ = 10.3, 8.2, 4.0 Hz, 1H), 4.22 (td, $J$ = 8.5, 5.1 Hz, 1H), 4.05 (dd, $J$ = 8.9, 6.3 Hz, 1H), 2.93 (dd, $J$ = 13.9, 4.0 Hz, 1H), 2.70 (d, $J$ = 6.7 Hz, 2H), 2.67 - 2.63 (m, 1H), 1.96 - 1.83 (m, 1H), 1.68 (s, 3H), 1.64 - 1.61 (m, 1H), 1.48 (ddt, $J$ = 14.7, 7.3, 4.4 Hz, 3H), 1.31 - 1.20 (m, 2H), 0.79 (d, $J$ = 6.8 Hz, 3H), 0.77 (d, $J$ = 6.8 Hz, 3H). $^{13}$C NMR (201 MHz, DMSO) $\delta$ 173.3, 172.0, 171.7, 169.8, 138.5, 129.6, 128.5, 126.7, 57.7, 54.5, 53.0, 39.2, 37.9, 31.5, 31.1, 27.0, 22.9, 22.7, 19.8, 18.3. HR-ESI (m/z): $[M + H]^+$ calcd for $C_{22}H_{36}N_5O_4$ 434.2762, found 434.2760.

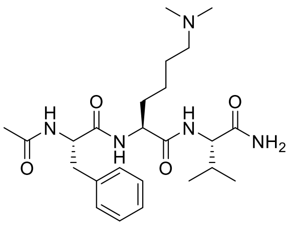

**6b**

**Compound 6b:** $^1$H NMR (800 MHz, DMSO-$d_6$) $\delta$ 8.26 (d, $J$ = 8.0 Hz, 1H), 8.12 (d, $J$ = 8.2 Hz, 1H), 7.63 (d, $J$ = 8.9 Hz, 1H), 7.42 - 7.40 (m, 1H), 7.29 – 7.25 (m, 4H), 7.19 (tt, $J$ = 5.4, 2.4 Hz, 1H), 7.09 - 7.04 (m, 1H), 4.52 (ddd, $J$ = 10.3, 8.2, 4.0 Hz, 1H), 4.35 – 4.28 (m, 1H), 4.13 (dd, $J$ = 8.9, 6.3 Hz, 1H), 3.00 (dd, $J$ = 14.0, 3.9 Hz, 1H), 2.85 (t, $J$ = 7.8 Hz, 2H), 2.73 (dd, $J$ = 14.0, 10.4 Hz, 1H), 2.64 (s, 6H), 1.96 (q, $J$ = 6.7 Hz, 1H), 1.72 (tt, $J$ = 7.2, 3.0 Hz, 1H), 1.57 (dd, $J$ = 9.8, 5.8 Hz, 3H), 1.30 (m, 2H), 0.87 (d, $J$ = 6.8 Hz, 3H),

0.84 (d, *J* = 6.8 Hz, 3H). $^{13}C$ NMR (201 MHz, DMSO) *δ* 173.2, 172.0, 171.6, 169.7, 138.5, 129.6, 128.5, 126.7, 57.7, 57.5, 54.4, 52.9, 43.2, 37.9, 31.7, 31.1, 22.9, 22.9, 19.8, 18.3. HR-ESI (m/z): $[M + H]^+$ calcd for $C_{24}H_{40}N_5O_4$ 462.3075, found 462.3065.

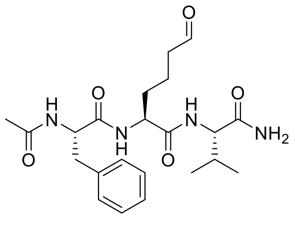
6c

**Compound 6c:** $^1H$ NMR (800 MHz, DMSO-*d*6) *δ* 9.58 (s, 1H), 8.12 (d, *J* = 7.9 Hz, 1H), 8.03 (d, *J* = 8.3 Hz, 1H), 7.56 (d, *J* = 9.0 Hz, 1H), 7.29 (s, 1H), 7.21 – 7.16 (m, 6H), 7.11 (dt, *J* = 5.0, 2.3 Hz, 1H), 6.97 (s, 1H), 4.50 - 4.40 (m, 1H), 4.22 (d, *J* = 5.4 Hz, 1H), 4.04 (dd, *J* = 8.9, 6.3 Hz, 1H), 2.92 (dd, *J* = 13.9, 4.0 Hz, 1H), 2.64 (dd, *J* = 13.9, 10.2 Hz, 1H), 2.37 (s, 1H), 1.90 - 1.84 (m, 1H), 1.70 - 1.68 (m, 3H), 1.60 (s, 1H), 1.47 (m, 2H), 1.22 (m, 1H), 0.78 (d, *J* = 6.8 Hz, 3H), 0.77 (s, 3H). $^{13}C$ NMR (201 MHz, DMSO) δ 203.7, 173.2, 172.0, 171.6, 169.6, 138.5, 129.6, 128.5, 126.7, 57.7, 54.3, 53.0, 43.1, 37.9, 31.6, 31.1, 22.9, 19.8, 18.5, 18.3. $[M + H]^+$ calcd for $C_{22}H_{33}N_4O_5$ 433.2446, found 433.2464.

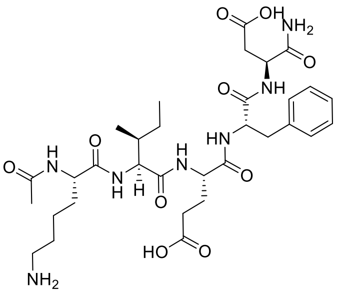
7a

**Compound 7a:** $^1H$ NMR (800 MHz, DMSO-*d*6) *δ* 8.13 (d, *J* = 7.9 Hz, 1H), 7.95 (d, *J* = 8.1 Hz, 1H), 7.91 (s, 1H), 7.87 (s, 1H), 7.74 (d, *J* = 8.3 Hz, 1H), 7.16 (d, *J* = 7.0 Hz, 4H), 7.11 (d, *J* = 6.9 Hz, 1H), 7.02 (s, 1H), 6.90 (s, 1H), 4.41 (q, *J* = 7.4 Hz, 1H), 4.36 (q, *J* = 7.1 Hz, 1H), 4.21 (q, *J* = 7.6 Hz, 1H), 4.15 (q, *J* = 7.4 Hz, 1H), 4.05 (t, *J* = 7.9 Hz, 1H), 2.95 (dd, *J* = 14.2, 4.9 Hz, 1H), 2.73 (dd, *J* = 14.0, 9.0 Hz, 1H), 2.68 (t, *J* = 7.7 Hz, 2H), 2.57 - 2.52 (m, 1H), 2.08 (m, 2H), 1.76 - 1.72 (m, 1H), 1.62 (h, *J* = 7.3, 6.6 Hz, 2H), 1.55 (h, *J* = 6.3 Hz, 1H), 1.46 - 1.38 (m, 3H), 1.32 (t, *J* = 8.7 Hz, 1H), 1.22 (m, 2H), 0.96 (m, 1H), 0.72 (t, *J* = 7.5 Hz, 3H), 0.69 (d, *J* = 6.7 Hz, 3H). $^{13}C$ NMR (201 MHz, DMSO) *δ* 174.6, 173.0, 172.8, 172.5, 172.2, 171.3, 171.1, 169.8, 138.0, 129.6, 128.5, 126.8, 57.3, 54.3, 52.6, 52.3, 50.1, 40.9, 39.2, 37.7, 36.8, 31.7, 30.6, 28.0, 27.1, 24.8, 23.0, 22.8, 15.8, 11.5. $[M + H]^+$ calcd for $C_{32}H_{50}N_7O_{10}$ 692.3614, found 692.3603.

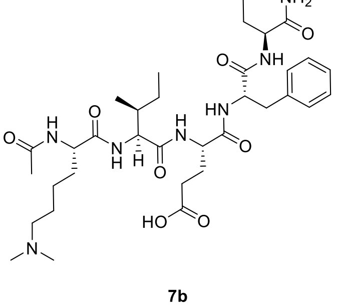
7b

**Compound 7b:** $^1H$ NMR (800 MHz, DMSO-*d*6) *δ* 8.14 (d, *J* = 7.9 Hz, 1H), 7.96 (d, *J* = 7.9 Hz, 1H), 7.93 (d, *J* = 7.7 Hz, 1H), 7.83 (d, *J* = 8.5 Hz, 1H), 7.81 (d, *J* = 7.8 Hz, 1H), 7.16 (d, *J* = 4.5 Hz, 4H), 7.10 (q, *J* = 4.3 Hz, 1H), 7.01 (s, 1H), 6.89 (s, 1H), 2.96 (dd, *J* = 14.0, 4.9 Hz, 1H), 2.83 - 2.79 (m, 2H), 2.73 (dd, *J* = 14.0, 9.2 Hz, 1H), 2.59 (s, 6H), 2.52 (dd, *J* = 16.3, 5.9 Hz, 1H), 2.44 - 2.40 (m, 3H), 2.13 - 2.01 (m, 2H), 1.77 (m, 4H), 1.66 - 1.61 (m, 2H), 1.59 -1.54 (m, 1H), 1.49

(p, $J$ = 7.9 Hz, 2H), 1.42 (dq, $J$ = 10.0, 5.4, 4.2 Hz, 1H), 1.32 (ddd, $J$ = 13.5, 7.3, 3.5 Hz, 1H), 1.18 (dq, $J$ = 16.2, 8.3 Hz, 2H), 1.01 - 0.94 (m, 1H), 0.72 (t, $J$ = 7.4 Hz, 3H), 0.69 (d, $J$ = 6.8 Hz, 3H). $^{13}$C NMR (201 MHz, DMSO) $\delta$ 174.7, 172.9, 172.7, 172.2, 171.4, 171.3, 171.1, 169.7, 138.0, 129.6, 128.5, 126.7, 57.4, 57.1, 54.4, 52.6, 52.2, 50.1, 42.8, 37.6, 36.9, 36.7, 31.9, 30.6, 28.0, 24.8, 24.1, 22.9, 15.8, 11.5. [M + H]$^+$ calcd for $C_{34}H_{54}N_7O_{10}$ 720.3927, found 720.3879.

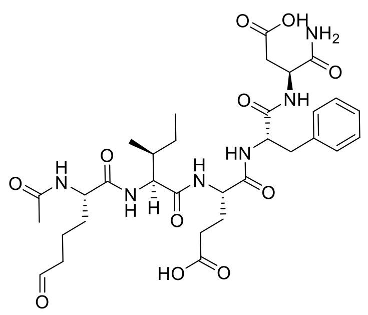


**7c**

**Compound 7c:** $^1$H NMR (800 MHz, DMSO-d$_6$) $\delta$ 9.55 (s, 1H), 8.15 (d, $J$ = 8.0 Hz, 1H), 7.93 (d, $J$ = 8.0 Hz, 1H), 7.89 - 7.84 (m, 2H), 7.79 (d, $J$ = 8.5 Hz, 1H), 7.15 (m, 4H), 7.10 (d, $J$ = 3.9 Hz, 1H), 7.02 (s, 1H), 6.89 (s, 1H), 4.41 - 4.39 (m, 1H), 4.37 (s, 1H), 4.23 – 4.19 (m, 1H), 4.15 (dt, $J$ = 8.0, 4.0 Hz, 1H), 4.05 (d, $J$ = 7.9 Hz, 1H), 2.93 (d, $J$ = 5.2 Hz, 1H), 2.74 (dd, $J$ = 9.0, 2.9 Hz, 1H), 2.56 (t, $J$ = 8.0 Hz, 2H), 2.40 (d, $J$ = 6.8 Hz, 1H), 2.33 (dt, $J$ = 7.9, 3.7 Hz, 1H), 2.06 (d, $J$ = 5.5 Hz, 2H), 1.74 (d, $J$ = 6.6 Hz, 1H), 1.60 (dd, $J$ = 8.7, 4.3 Hz, 2H), 1.53 (d, $J$ = 5.0 Hz, 1H), 1.46 (s, 1H), 1.44 - 1.40 (m, 1H), 1.32 (m, 2H), 0.97 - 0.94 (m, 1H), 0.71 (d, $J$ = 7.5 Hz, 3H), 0.68 (d, $J$ = 7.0 Hz, 3H). $^{13}$C NMR (201 MHz, DMSO) $\delta$ 203.7, 172.7, 172.3, 172.1, 171.4, 171.3, 171.1, 168.7, 137.9, 129.6, 128.5, 126.7, 57.3, 54.3, 52.7, 52.3, 50.0, 43.1, 40.5, 38.0, 37.8, 36.7, 36.4, 31.8, 30.5, 27.9, 24.8, 24.6, 22.9, 18.5, 15.8, 11.5. [M + H]$^+$ calcd for $C_{32}H_{47}N_6O_{11}$ 691.3341, found 691.3297.

#### 4. Quantification datasets of model peptides and protein

##### 4.1 Quantification of model peptides

Lys- and allysine-containing model peptides were separately prepared at paired concentrations of 1980/20, 1800/200, 1500/500, 1000/1000, 500/1500, and 200/1800 μM, respectively. For labeling of the Lys-containing peptide, each reaction contained peptide solution (1 μL), Ald-alkyne (1 μL, 5 mM), PBS (47 μL, pH 5.0), and $NaBH_3CN$ (1 μL, 0.1 M), giving a total reaction volume of 50 μL. The allysine-containing peptide was labeled separately under the same conditions using Am-alkyne (1 μL, 5 mM) instead of Ald-alkyne. The reactions were incubated at 37$^o$C for 1h. Following labeling, the Lys-derived samples were conjugated with heavy IsoDTB tags, whereas the allysine-derived samples were conjugated with light IsoDTB tags. Each click reaction was supplemented with the corresponding IsoDTB tag solution (2.5 μL, 20 mM) and click reaction mixture (7.5 μL), followed by incubation at 37$^o$C for 1h. After the click reactions, the corresponding heavy- and light-labeled samples were combined and desalted using C18 tips before analysis by LC-MS/MS analysis. LC-MS/MS analysis was performed using a Thermo Scientific Orbitrap Exploris 240 mass spectrometer. Mobile phase A consisted of 0.1% formic acid in water, and mobile phase B consisted of 80% acetonitrile containing 0.1% formic acid. Peptides were separated using a 60 min gradient: 4-5% B over 0.5 min, 5-28% B over 44 min, 28-42% B over 4 min, and 42-55% B over 4 min, followed by a wash at 99% B during the final 7.5 min. Full MS scans were acquired over an m/z range of 375-1475 at a resolution of 120,000, with an automatic gain control (AGC) target of $3 \times 10^6$. Data-dependent MS/MS acquisition was performed using a 3 s cycle time and a dynamic exclusion duration of 45 s. MS/MS spectra were acquired at a resolution of 30,000, with an AGC target of $2 \times 10^5$ and an HCD collision energy of 26%. The expected allysine stoichiometry was calculated as the molar amount of the allysine-containing peptide divided by the combined molar amounts of the Lys- and allysine-containing peptides. Measured allysine stoichiometry was calculated from the corresponding light and heavy signals as follows: Allysine stoichiometry (%) = L/(L + H) × 100 %, where L and H represent the signals of the light-labeled allysine-derived peptide and the heavy-labeled Lys-derived peptide, respectively. Calibration curves were generated by plotting the measured occupancies against the expected occupancies and fitting the data by linear regression.

##### 4.2 In-gel fluorescence analysis lysozyme and BSA with or without TACO chemistry

Native lysozyme and BSA (5 mg/mL, 1 mL) were dissolved in PBS containing 1.2% SDS (pH 5.0) and incubated with 5 mM probe and 100 mM NaBH3CN at 37 °C for 1 h. Proteins were then precipitated with methanol/chloroform, and the resulting pellets were washed with cold methanol and resuspended in PBS containing 1.2% SDS. Copper-catalyzed azide-alkyne cycloaddition (CuAAC) was performed by adding azido-rhodamine (100 μM), freshly prepared tris(2-carboxyethyl)phosphine hydrochloride (TCEP, 1 mM), tris[(1-benzyl-1H-1,2,3-triazol-4-yl)methyl]amine (TBTA, 0.1 mM), and $CuSO_4$ (1 mM). The reaction mixtures were incubated at 37 °C for 1 h in the dark with gentle agitation and then for in-gel fluorescence. For TACO treatment, native lysozyme and BSA (5 mg/mL, 1 mL) were incubated with 10 μL of 37% formaldehyde and 100 mM NaBH3CN at 37 °C for 1 h. Proteins were then precipitated with methanol/chloroform and redissolved in 10 mM NaP buffer (pH 7.0), followed by the addition of pyridine (10 μL) and Selectfluor (2 mg). The reaction mixtures were stirred for 1 h, after which the proteins were precipitated and subjected to probe labeling and CuAAC as described above.

##### 4.3 Proteomic analysis Lys state of BSA with or without TACO chemistry

Native lysozyme and BSA (5 mg/mL, 1 mL) were dissolved in PBS containing 1.2% SDS (pH 5.0) and incubated with 5 mM Am-alkyne or Ald-alkyne in the presence of 100 mM NaBH3CN at 37 °C for 1 h. Proteins were then precipitated with methanol/chloroform, and the resulting pellets were washed with cold methanol and resuspended in PBS containing 1.2% SDS. CuAAC was performed by adding light or heavy IsoDTB tags to the Am-alkyne- or Ald-alkyne-labeled samples, respectively, followed by the click reagent mixture. The reaction mixtures were incubated at 37 °C for 1 h in the dark with gentle agitation. Following CuAAC, the light- and heavy-labeled samples were combined and subjected to methanol/chloroform precipitation, followed by proteomic analysis as described in Section 6. Chemical proteomics study. For TACO treatment, native lysozyme and BSA (5 mg/mL, 1 mL) were incubated with 10 µL of 37 % formaldehyde and 100 mM NaBH3CN at 37 °C for 1 h. Proteins were then precipitated with methanol/chloroform and redissolved in 10 mM NaP buffer (pH 7.0), followed by the addition of pyridine (10 µL) and Selectfluor (2 mg). The reaction mixtures were stirred for 1 h, after which the proteins were precipitated and subjected to probe labeling, CuAAC, and proteomic analysis as described above.

**5. Biological experiments materials and methods**

**5.1 Cell culture**

HEK293T (CRL-3216) cells were obtained from the ATCC. HT1080 (CCL-121) cells were kind gifts from Dr. Zhipeng Meng, respectively. HEK293T and HT1080 cells were cultured in DMEM (Gibco, Invitrogen), supplemented with 10% fetal bovine serum (FBS) and 100 units/ml penicillin/streptomycin at 37°C in a humidified incubator under 5% $CO_2$. All cell culture reagents were obtained from ThermoFisher, except for sera, FBS, which were obtained from GeminiBio. Cell lines were tested negative for mycoplasma and authenticated by short tandem repeat (STR) profiling.

**5.2 Benign prostatic hyperplasia (BPH)tissue collection**

De-identified BPH tissue specimens were collected from patients undergoing holmium laser enucleation of the prostate (HoLEP) at the University of Miami. Fresh tissue specimens were dissected into smaller portions and immediately flash-frozen for subsequent proteomic analysis. Specimens from patients with an incidental diagnosis of prostate cancer were excluded from the study. The study was approved by the University of Miami Institutional Review Board (IRB #20250162), and written informed consent was obtained from all participants prior to specimen collection. All study procedures were conducted in accordance with institutional ethical guidelines.

##### 5.3 In-gel fluorescence analysis of probe 1-4

For each probe, 200 μg of cell lysate in 0.1% Triton X-100/PBS was precipitated using methanol/chloroform and washed with cold methanol. The resulting protein pellet was resuspended in 1.2% SDS/PBS (pH 5.0) and incubated with 5 mM probe and 100 mM NaBH3CN for 1 h at 37 °C. Following incubation, the reaction mixture was precipitated using methanol/chloroform, and the pellet was washed with cold methanol. The pellet was then resuspended in 1.2% SDS/PBS, and CuAAC was performed by adding 100 μM azido-rhodamine, 1 mM freshly prepared tris(2-carboxyethyl)phosphine hydrochloride (TCEP), 0.1 mM TBTA, and 1 mM $CuSO_4$. The reaction was incubated for 1 h at 37 °C in the dark with gentle agitation. Reactions were terminated by adding 4× Laemmli sample buffer, followed by heating at 95 °C for 10 min. Samples were separated by 10% SDS-PAGE, and fluorescently labeled proteins were visualized using a ChemiDoc MP Imaging System (Bio-Rad) equipped with a rhodamine filter. After fluorescence imaging, the gel was stained with Coomassie Brilliant Blue to visualize total protein.

##### 5.4 Lipid peroxidation assay

The RSL3-induced lipid peroxidation was measured by using the BODIPY 581/591 C11 probe (Thermo Fisher Scientific). Briefly, HT1080 cells plated in 10cm dishes, twenty-four hours after plating, the cells were treated with vehicle control or 150 nM RSL3 or 150 nM RSL3 with 300 nM Liproxstatin-1. Three hours after treatment, the cells were harvested by trypsinization, resuspended in PBS, and incubated with 10 μM BODIPY C11 at 37°C for 30 minutes. The cell suspension was then loaded into 96-well black-walled plates and measured the fluorescence signal using a microplate reader. Lipid peroxidation was determined by calculating the ratio of oxidized (Ex/Em = 485:535 nm) to reduced (Ex/Em = 560/600 nm) C11 fluorescence signal.

#### 6. Chemical proteomics study

##### 6.1 Chemical proteomics of probe 1-4 (with DADPS tag)

For each probe, 5mg of 293T cell lysate in 0.1% Triton X-100/PBS was precipitated using methanol/chloroform and washed with cold methanol. The resulting protein pellet was resuspended in 1.2% SDS/PBS (pH 5.0) and incubated with 5 mM probe and 100 mM NaBH3CN for 1 h at 37 °C. Following incubation, the reaction mixture was precipitated using methanol/chloroform, and the pellet was washed with cold methanol. The pellet was then resuspended in 1.2% SDS/PBS and then clicked with acid cleavable azido-DADPS-biotin in the presence of TCEP, TBTA, and $CuSO_4$ as described above. Methanol precipitated and washed protein pellets were again resuspended in 1.2% SDS/PBS buffer. Then pre-washed streptavidin agarose beads (~50 µL slurry; ThermoFisher) were added to each sample. The protein and beads mixtures were incubated at room temperature on a rotator for 4 h. The beads were then washed 3 times with PBS and then 3 times with dd$H_2O$. The beads were then added with 6 M urea/PBS and then incubated with 10 mM dithiothreitol for 30 minutes, followed by treatment with 20 mM iodoacetamide for another 30 minutes. After that, the beads were washed with PBS and digested with 2 µg of trypsin in PBS buffer at 37 °C overnight. The beads were again washed 6 times with PBS and dd$H_2O$. Finally, the beads were resuspended with 2% formic acid in dd$H_2O$ to cleave the DADPS linker. The elution was repeated twice with 0.1% formic acid in dd$H_2O$ and the supernatants were pooled and dried by SpeedVac for LC-MS/MS analysis.

##### 6.2 Chemical proteomics of probes with IsoDTB tag

For probe 2 (Am-alkyne), the same procedure was performed except the lysate was clicked with isoDTB light tag in the presence of TCEP, TBTA, and $CuSO_4$ as described above. Methanol precipitated and washed protein pellets were dissolved in 300 µl urea (8 M in 0.1 M triethylammonium bicarbonate (TEAB)) by sonification. Then, 900 µl TEAB (0.1 M in $H_2O$) was added, and the solution was added to 1.2 ml of pre-washed streptavidin agarose beads (~50 µL slurry; ThermoFisher) in 0.2% NP40 in PBS. The samples were rotated for 1 h at room temperature on a rotator for 1 h. The beads were then washed 3 times with PBS and then 3 times with dd$H_2O$. The beads were resuspended in 600 µl 8 M urea in 0.1 M TEAB, with 10 mM dithiothreitol for 30 minutes, followed by treatment with 20 mM iodoacetamide for another 30 minutes. After that, the beads were washed with 0.1 M TEAB and digested with 2 µg of trypsin in 2 M Urea in 0.1 M TEAB buffer at 37 °C overnight. The beads were washed with 3 times with 0.1% NP40/PBS, 3 times with PBS and 3 times with dd$H_2O$. Peptides were eluted into Protein LoBind tubes with 1 × 400 µl and 2 × 200 µl 0.1% formic acid in 50% acetonitrile. The supernatants were pooled and dried by SpeedVac for LC-MS/MS analysis.

##### 6.3 Chemical proteomics of PRISM-lys platform

To quantify the relative abundance of allysine and unmodified Lys, the PRISM-lys platform was applied to 293T cell and benign prostatic hyperplasia (BPH) tissue lysates. Briefly, 5 mg of lysate in 0.1% Triton X-100/PBS was precipitated using methanol/chloroform and washed with cold methanol. The resulting protein pellet was resuspended in 1.2% SDS/PBS (pH 5.0) and incubated with either 5 mM Am-alkyne or 5 mM Ald-alkyne in the presence of 100 mM NaBH3CN for 1 h at 37 °C. Following incubation, both reaction mixtures were precipitated using methanol/chloroform, and the resulting pellets were washed with cold methanol. The pellets were then resuspended in 1.2% SDS/PBS. The Am-alkyne-labeled sample was conjugated to the isoDTB light tag, whereas the Ald-alkyne-labeled sample was conjugated to the isoDTB heavy tag via click chemistry in the presence of TCEP, TBTA, and $CuSO_4$, as described above. Following labeling, the light- and heavy-labeled samples were combined, precipitated using methanol/chloroform, and washed with cold methanol. The combined protein pellet was processed as described above in **4.1** for subsequent LC-MS/MS analysis.

##### 6.4 LC-MS/MS analysis

LC-MS/MS was performed on an Exploris 480 Orbitrap mass spectrometer (Thermo Fisher Scientific) coupled with a Vanquish Neo UHPLC. Mobile phase A was 0.1% formic acid in $H_2O$, and mobile phase B was 0.1% formic acid, 80% acetonitrile in $H_2O$. Flow rate was 10 μL/min for loading and 0.3 μL/min for eluting. Under the positive-ion mode, full-scan mass spectra were acquired over the m/z range from 350 to 1600 using the Orbitrap mass analyzer with mass resolution of 120,000. MS/MS fragmentation was performed in a data dependent mode, of which the most intense ions in a 3-second cycle was selected for MS/MS analysis a resolution of 30,000 using collision mode of HCD. Other important mass parameters: isolation window, 1.6 m/z units; default charge, 2+; normalized collision energy, 26%; dynamic exclusion, 45.0 s.

##### 6.5 Data analysis

###### 6.5.1 open and close search

Search of probe modified sites was performed by using FragPipe 24.0 with MSFragger (version 4.4.1), Philosopher (version v5.1.3), and IonQuant (version 1.11.20).[1-3] Precursor mass tolerance was set -20 to 20 ppm. Missed cleavages were allowed up to 2. Peptide length was set 7 to 50, and peptide mass range was set 500 to 5000. Cysteine carboxyamidomethylation (+57.02146 Da) was set as fixed modification. 15.9949 Da is set as variable modification on methionine. Peptides were required to achieve a peptide false-positive rate below 1%. Subsequently, a closed search of allysine sites was performed using the same software and search parameters, with Lys modification (+223.1685 Da) specified as a variable modification to identify probe-modified peptides. For IsoDTB tag search, a closed search of allysine sites was performed using the same software and search parameters, with Lys modification +561.3387 Da and +567.3462 Da specified as a variable modification to identify probe-modified peptides.

###### 6.5.2 quantitative search of IsoDTB-labelled peptides

Database searching and isotope-label-based quantification. Raw LC-MS/MS data were processed using FragPipe (24.0) with the built-in isoDTB-ABPP workflow adapted for Lys labeling. Database searches were performed using MSFragger (4.4.1) against the human database. The precursor mass tolerance was set to −20 to +20 ppm, and the fragment mass tolerance was set to 20 ppm, with mass calibration and parameter optimization enabled. Precursor isotope errors of 0, +1, and +2 were allowed. Trypsin specificity was defined as cleavage C-terminal to Lys or arginine, except when followed by proline, with up to two missed cleavages permitted. N-terminal methionine clipping was enabled. Peptides containing 6–50 amino acids and having masses of 500-5,000 Da were considered. Cysteine carboxyamidomethylation (+57.02146 Da) was set as fixed modification. Variable modifications included methionine oxidation (+15.9949 Da), protein N-terminal acetylation (+42.0106 Da), and the light (+561.3387 Da) or heavy (+567.3462 Da) probe-IsoDTB adduct on Lys. A maximum of three variable modifications per peptide was allowed, with up to two methionine oxidations, one protein N-terminal acetylation, and one occurrence of each isotope-labeled Lys modification per peptide. MSBooster was enabled to generate retention-time and fragment-intensity predictions using the DIA-NN models, and peptide-spectrum matches were rescored using Percolator. Protein inference was performed using ProteinProphet, followed by sequential filtering in Philosopher with the protein-level false discovery rate set to 1%. MS1-based isotope-label quantification was performed using IonQuant, with the light and heavy channels defined as K561.3387 and K567.3462, respectively. Feature detection and peak tracing used an m/z tolerance of 10 ppm and a retention-time tolerance of 0.4 min, requiring at least one MS1 scan and two isotopic peaks per feature.

The mass spectrometry proteomics data have been deposited to the ProteomeXchange Consortium (http://proteomecentral.proteomexchange.org) via the iProX[4-5] partner repository with the dataset identifier PXD086375 (iProx identifier IPX0020316000).

##### 6.5.3 Statistical analyses

Statistical analyses were performed using statistical software package GraphPad Prism (v. 11). Data are presented as means ± SEM. Differences between two samples were assessed for statistical significance using a two-tailed, unpaired Student's t-test with Welch's correction for unequal variances. The correlation between two replicates was evaluated using Pearson's correlation coefficient (r) to assess the strength of the linear relationship. The results with P values $< 0.05$ were considered statistically significant; ns, not significant; *$p < 0.05$; **$p < 0.01$; ***$p < 0.001$; ****$p < 0.0001$.

### 7. Characterization of the compounds

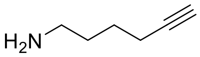


**Am-alkyne (2)**

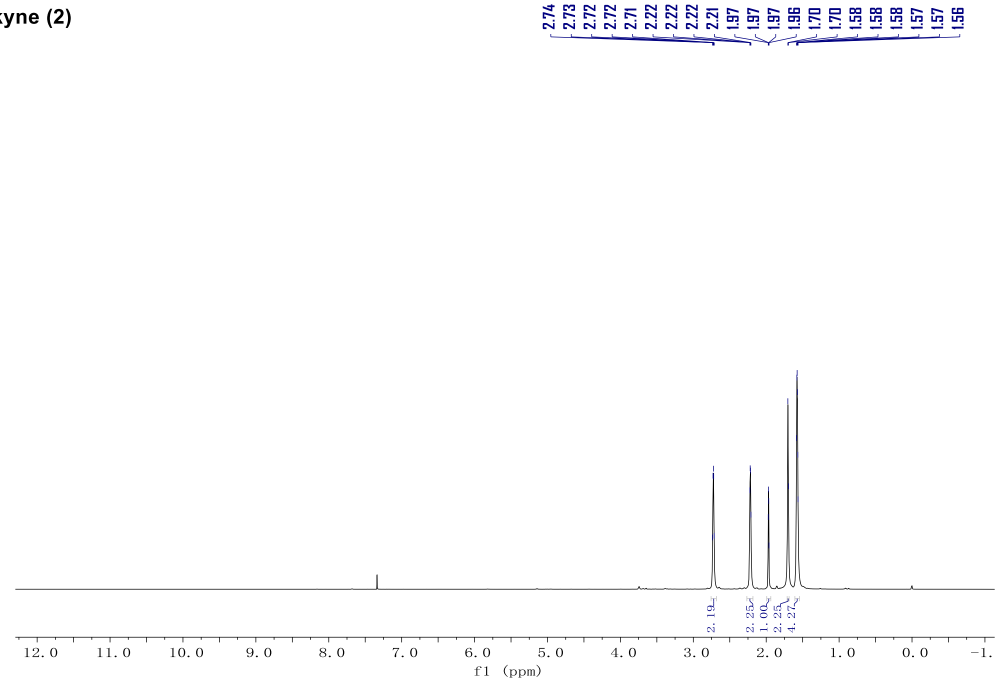


$^{1}$H NMR (500 MHz) spectrum of Am-alkyne in $CDCl_3$.

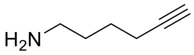


**Am-alkyne (2)**

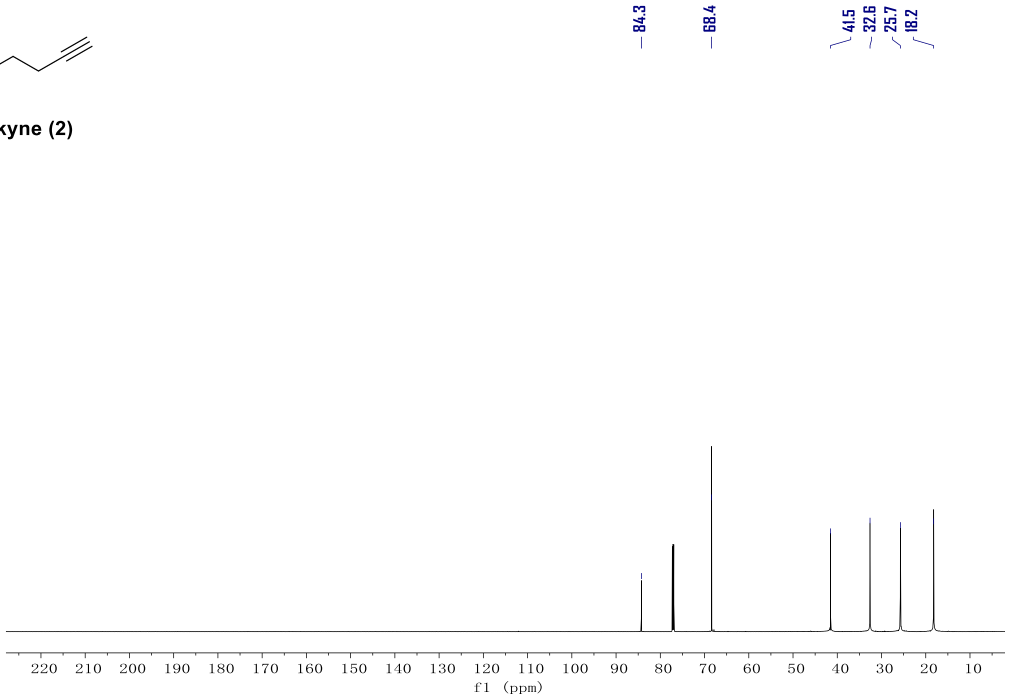


$^{13}$C NMR (126 MHz) spectrum of Am-alkyne in $CDCl_3$.

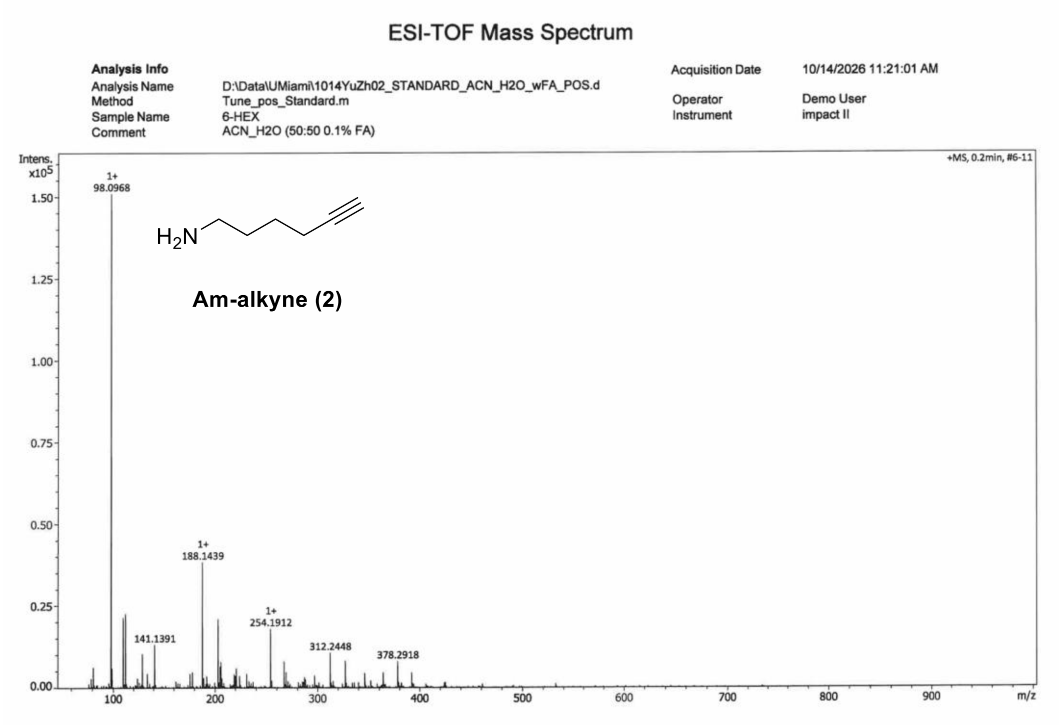


HR-MS spectrum of Am-alkyne.

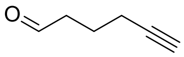


**Ald-alkyne (5)**

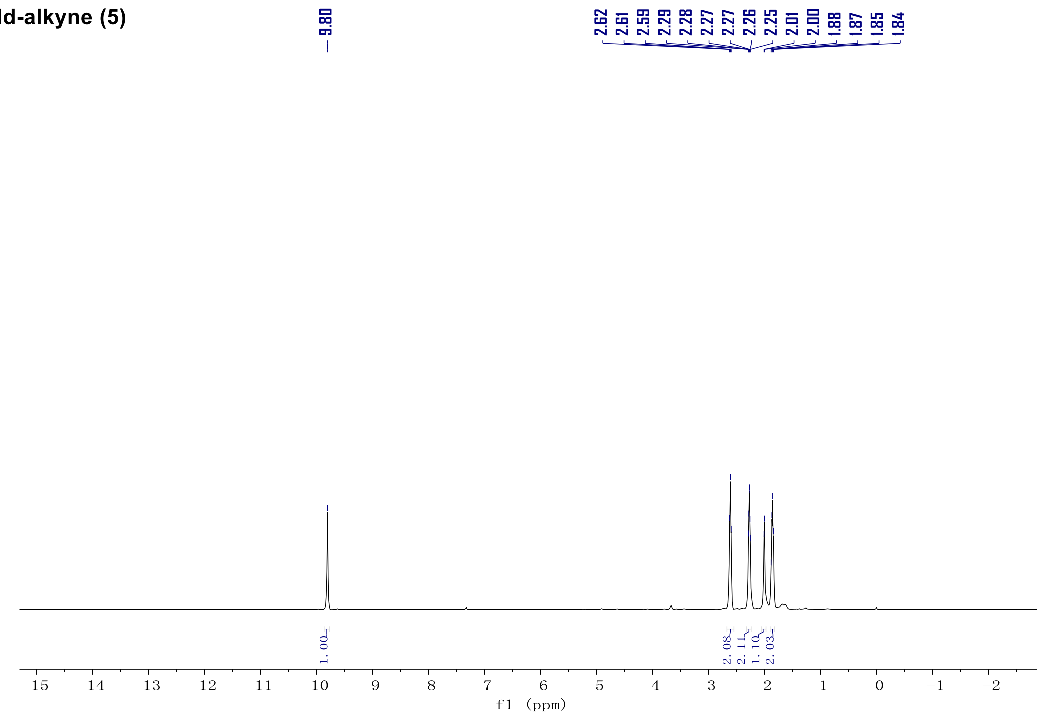


$^1$H NMR (500 MHz) spectrum of Ald-alkyne in $CDCl_3$.

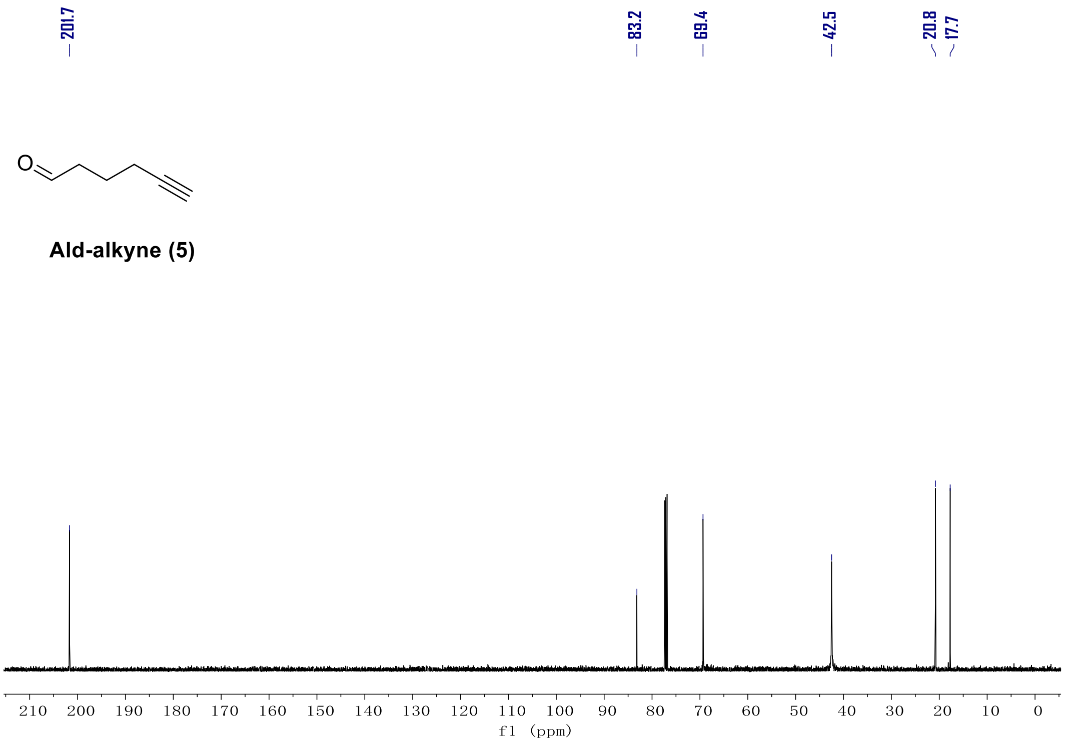


$^{13}C$ NMR (126 MHz) spectrum of Ald-alkyne in $CDCl_3$.

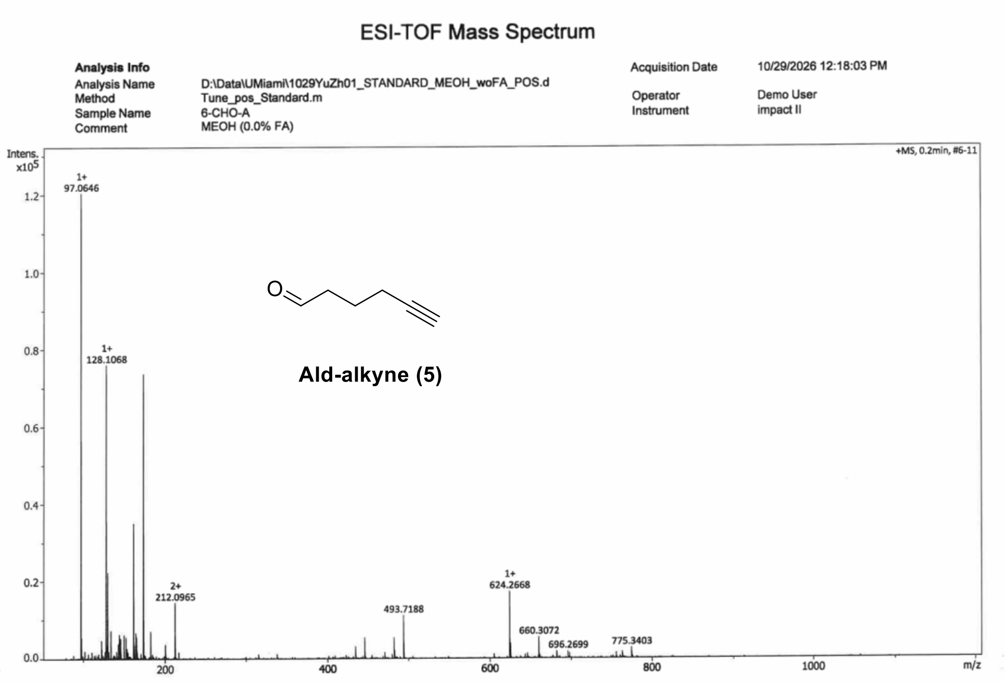


HR-MS spectrum of Ald-alkyne.

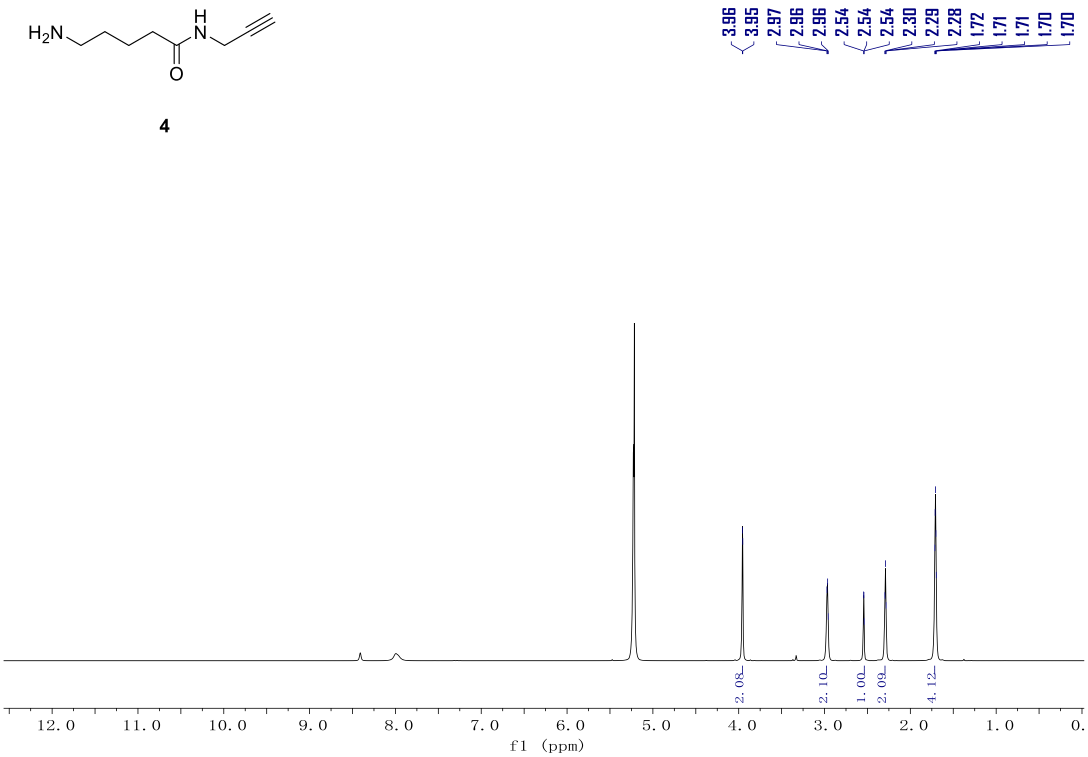


$^{1}H$ NMR (500 MHz) spectrum of **4** in $CD_3OD$.

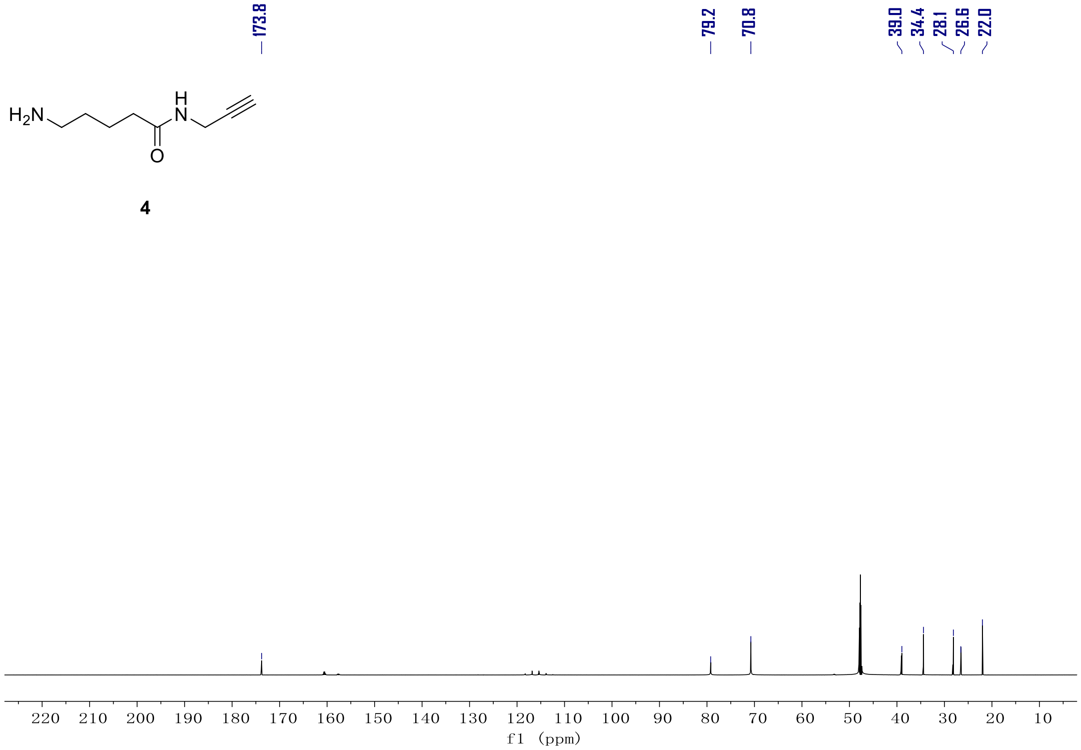


$^{13}C$ NMR (126 MHz) spectrum of **4** in $CD_3OD$.

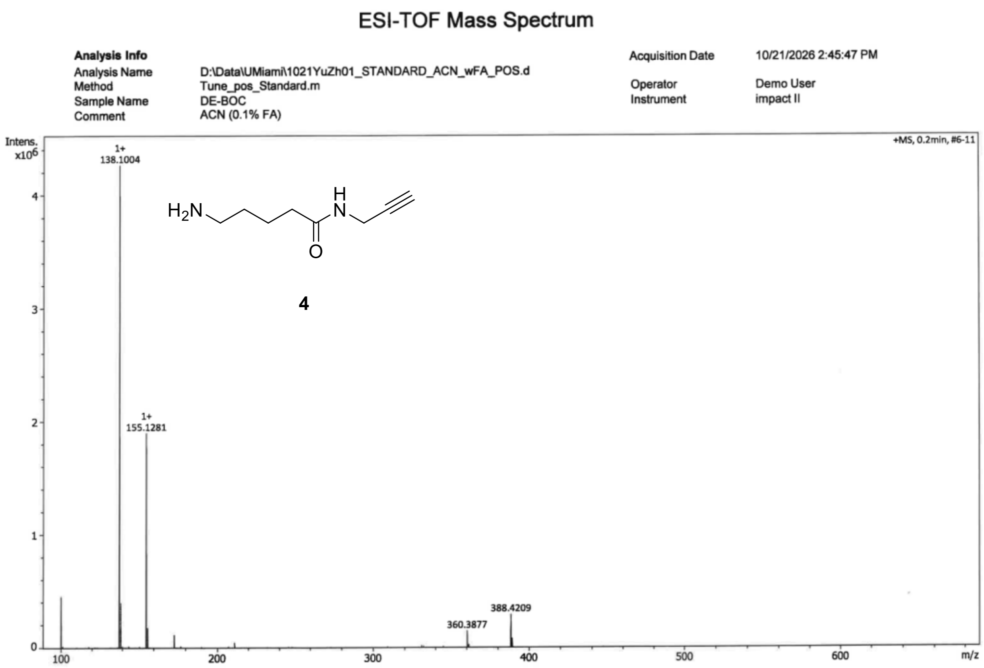


HR-MS spectrum of **4**.

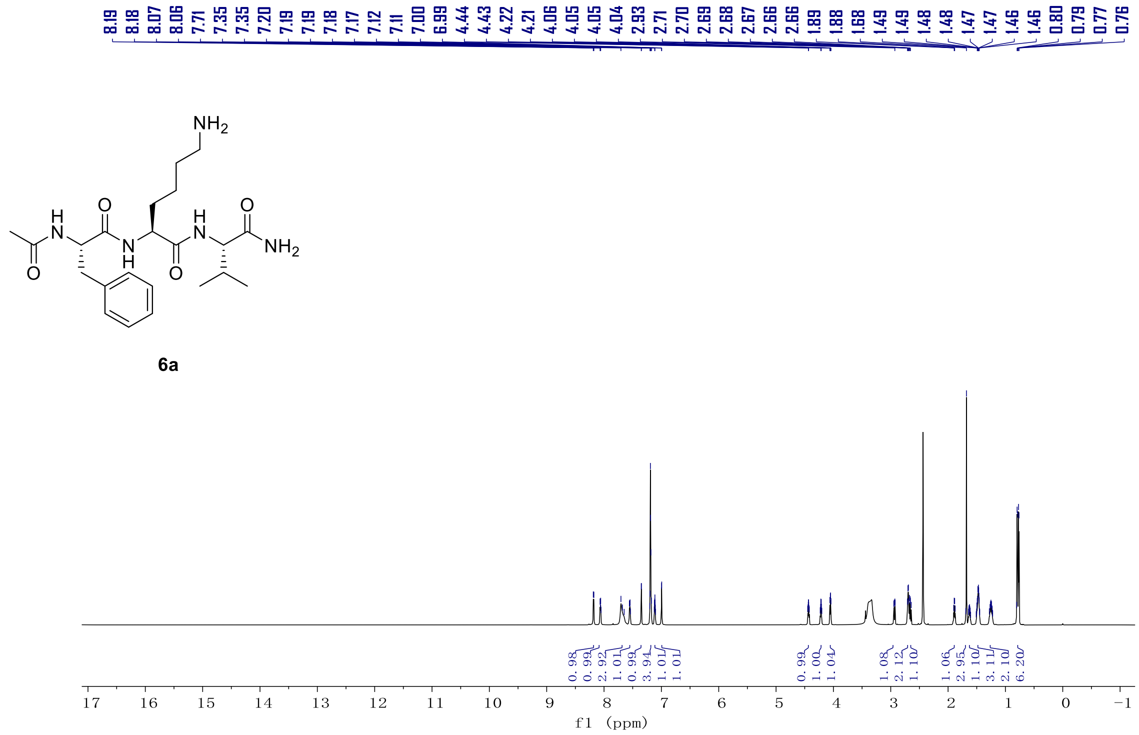


$^1$H NMR (800 MHz) spectrum of **6a** in DMSO-$d_6$.

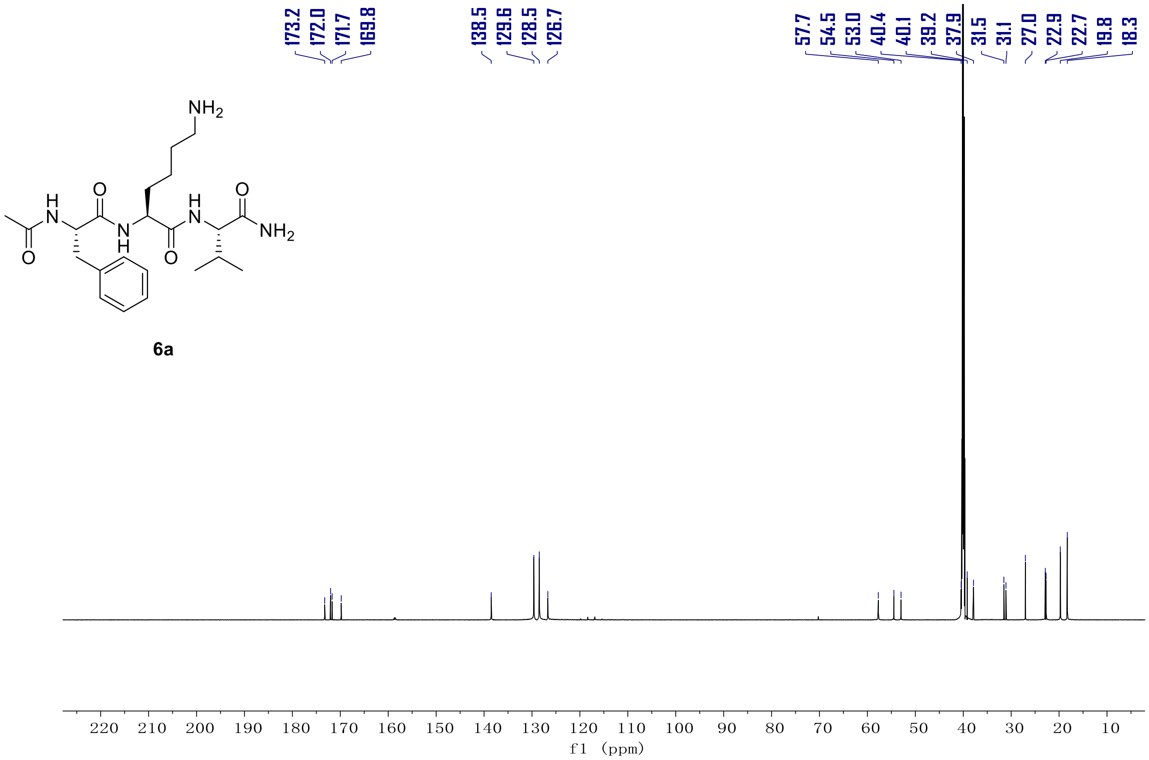


$^{13}C$ NMR (201 MHz) spectrum of **6a** in DMSO-d$_6$.

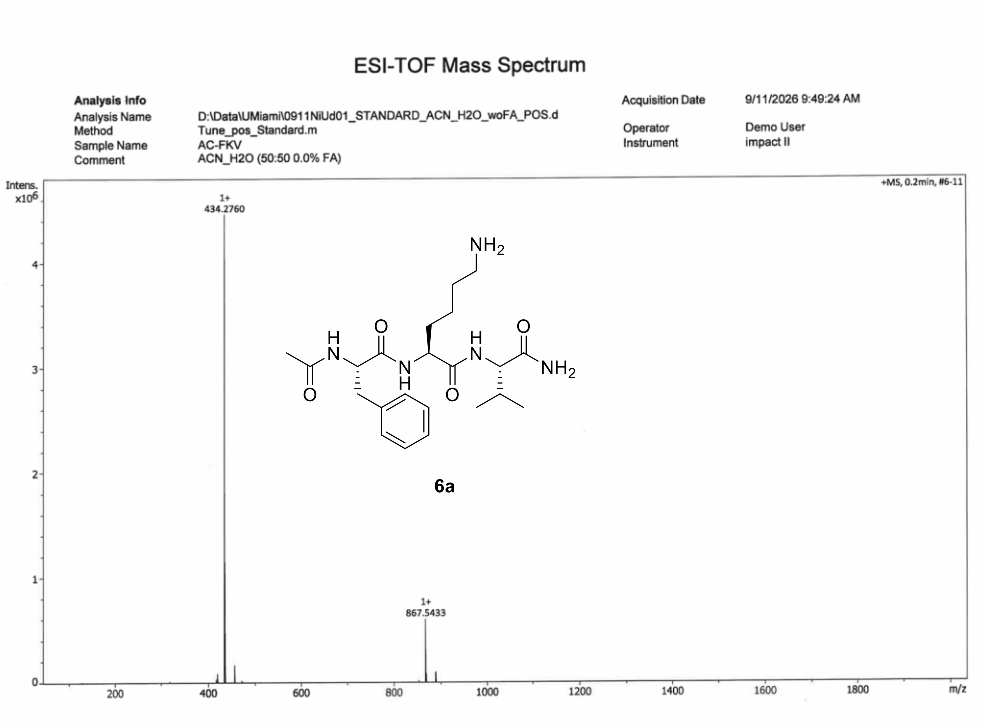


HR-MS spectrum of **6a**.

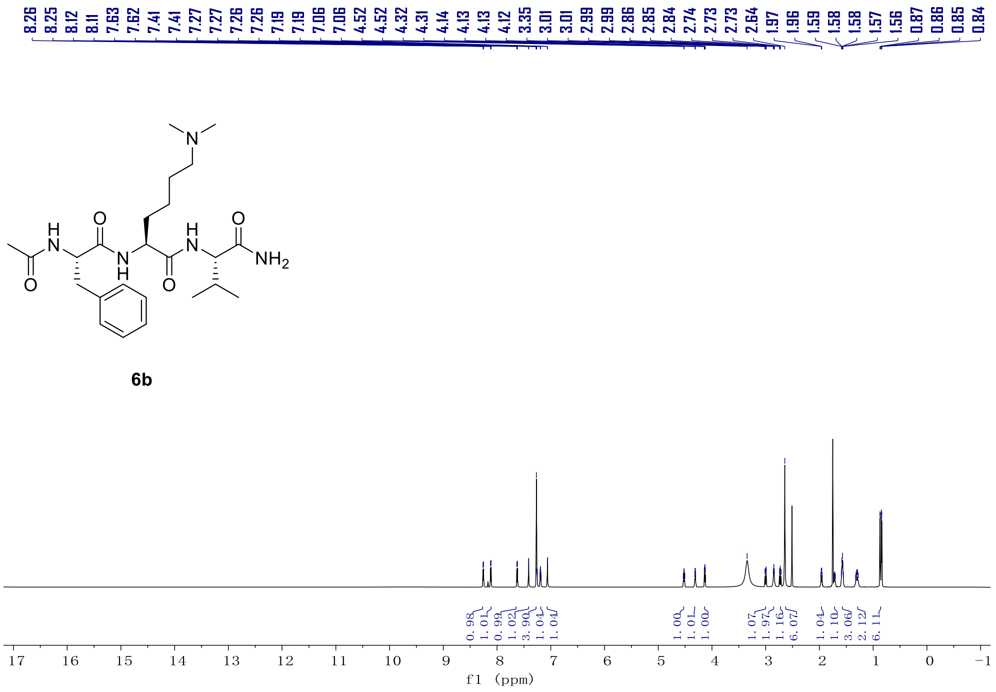


$^{1}$H NMR (800 MHz) spectrum of **6b** in DMSO-d$_6$.

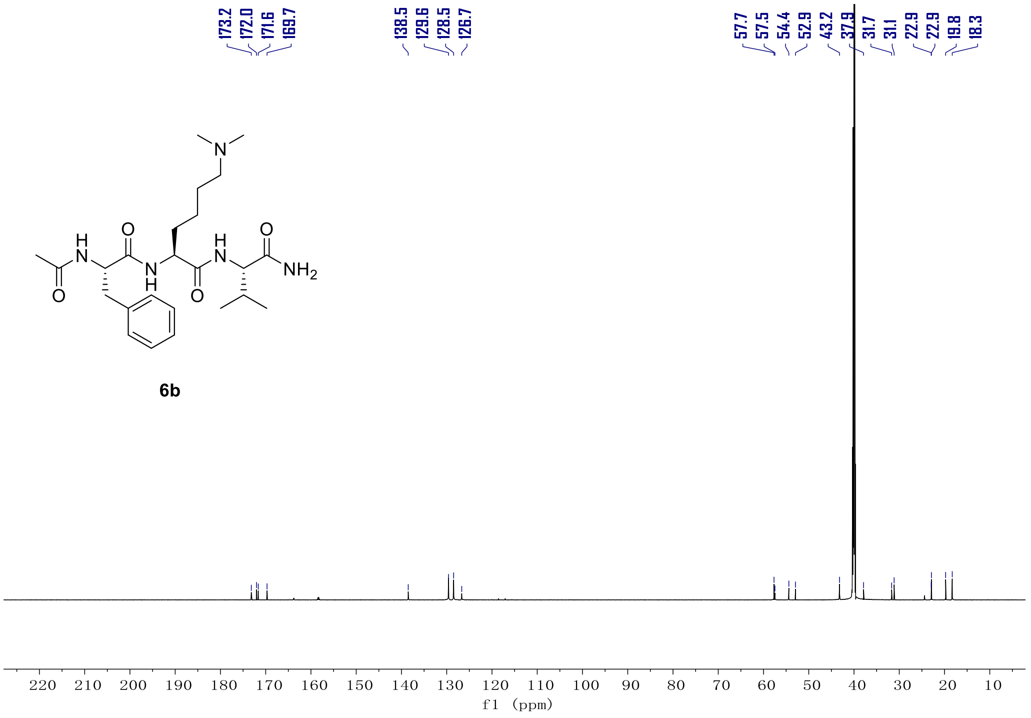


$^{13}$C NMR (201 MHz) spectrum of **6b** in DMSO-d$_6$.

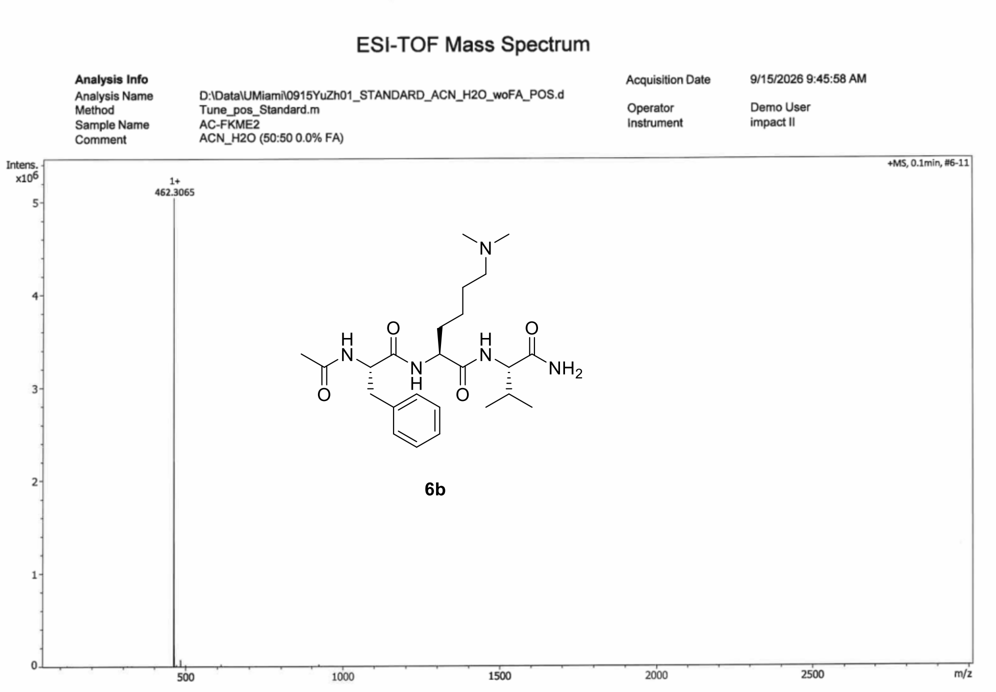


HR-MS spectrum of **6b**.

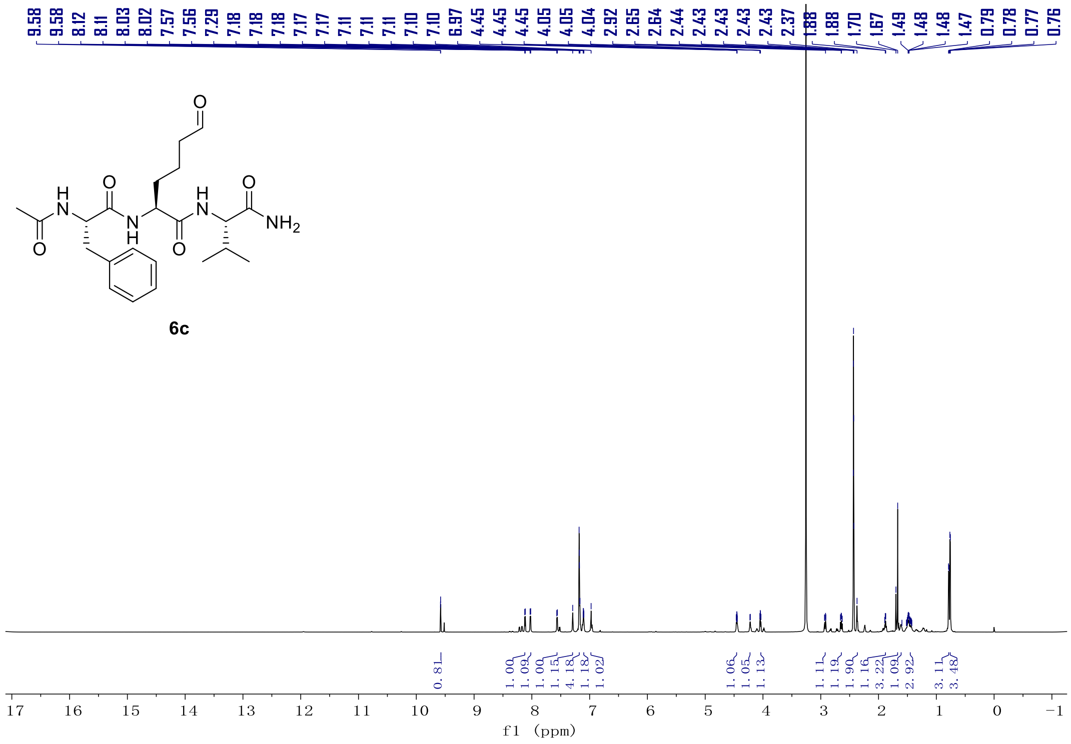


[1]H NMR (800 MHz) spectrum of **6c** in DMSO-$d_6$.

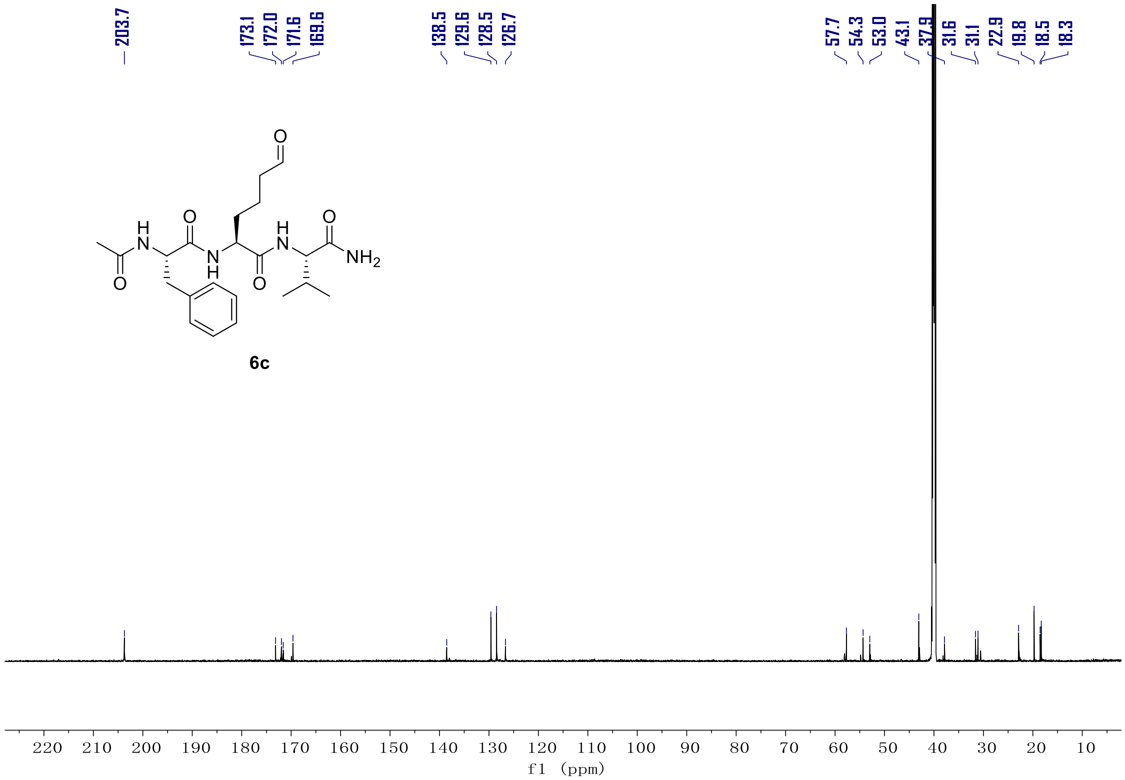


$^{13}C$ NMR (201 MHz) spectrum of **6c** in DMSO-d6.

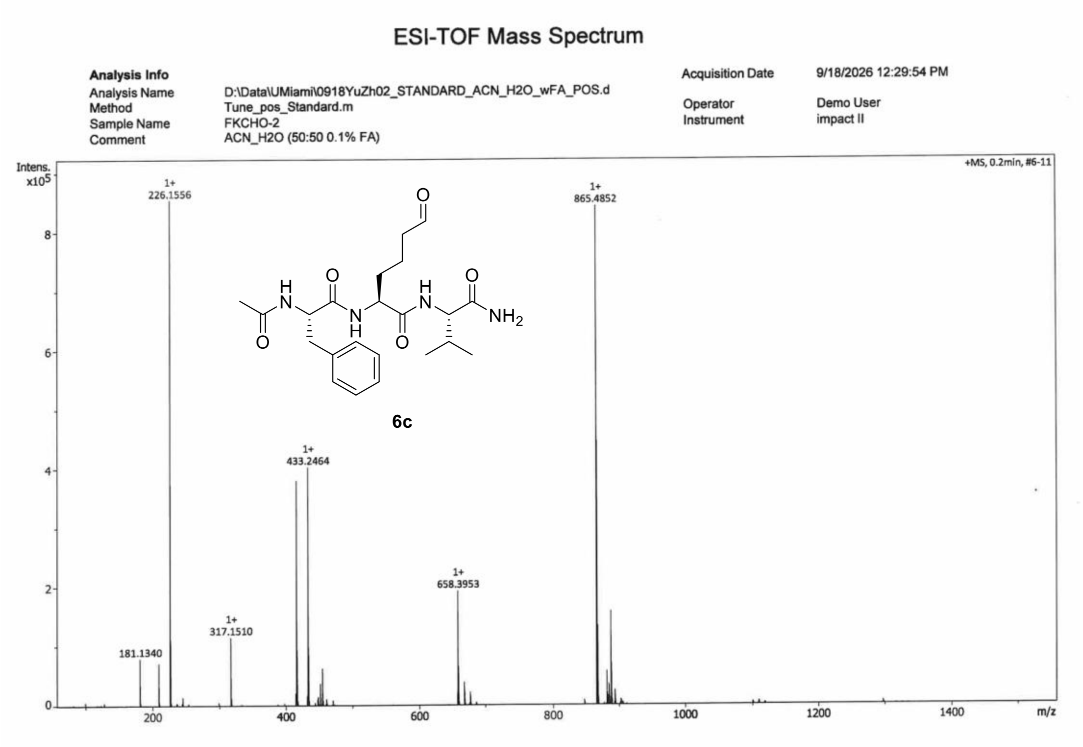


HR-MS spectrum of **6c**.

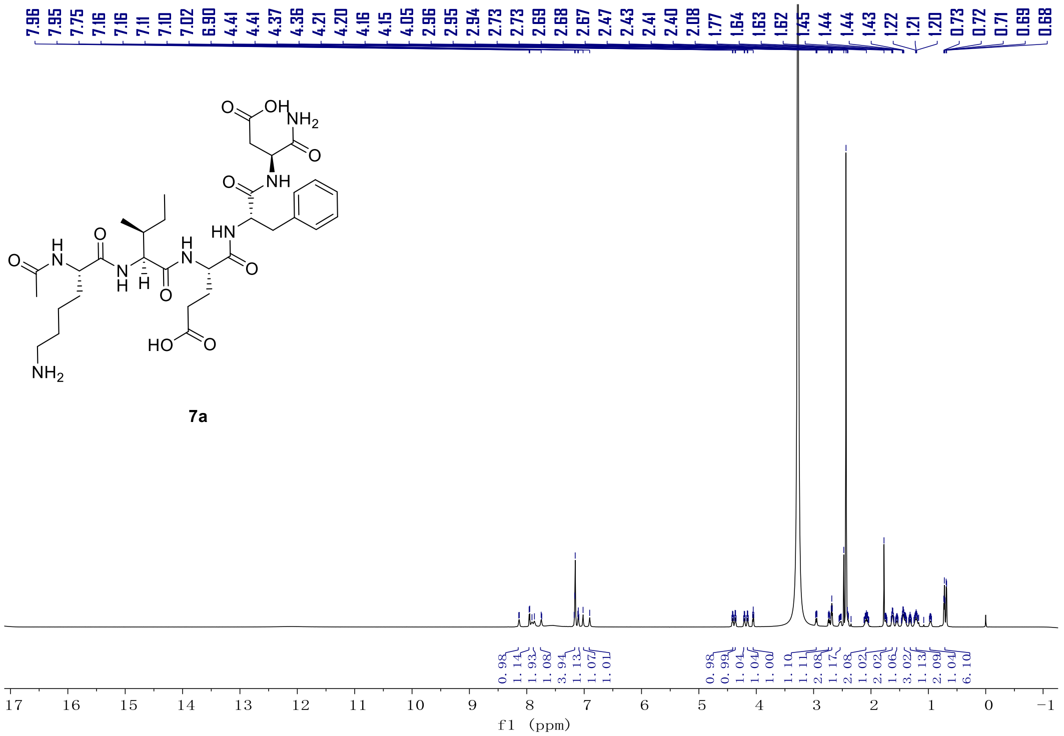


$^{1}$H NMR (800 MHz) spectrum of **7a** in DMSO-d6.

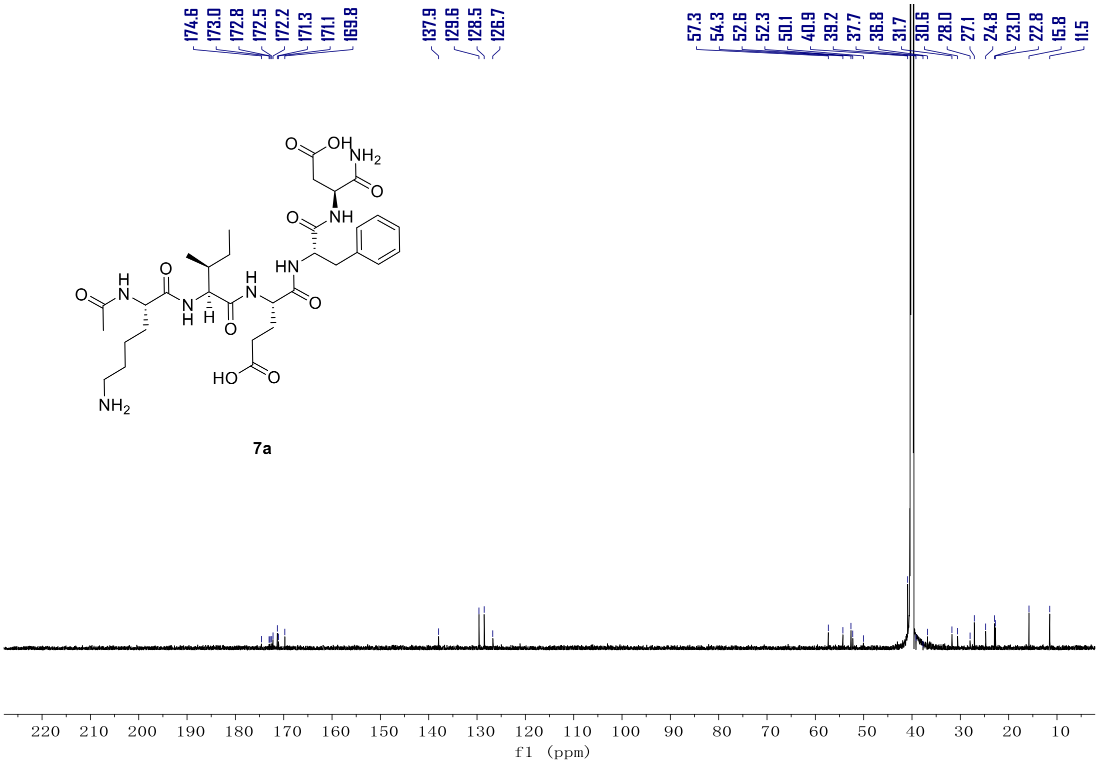


$^{13}$C NMR (201 MHz) spectrum of **7a** in DMSO-d6.

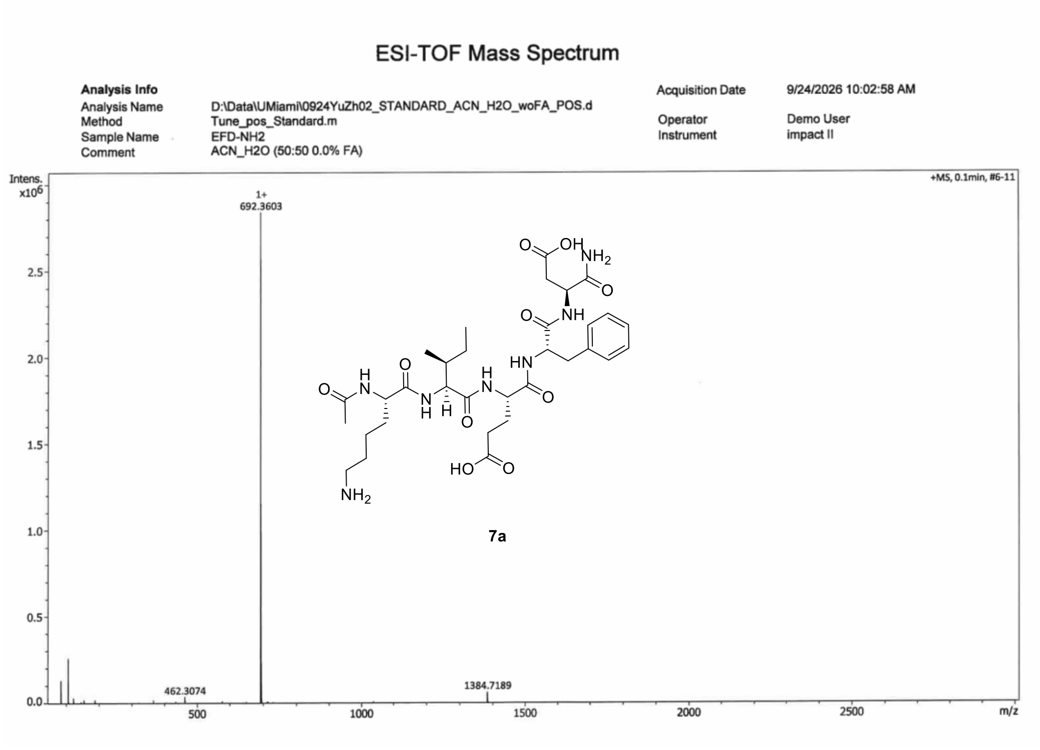


HR-MS spectrum of **7a**.

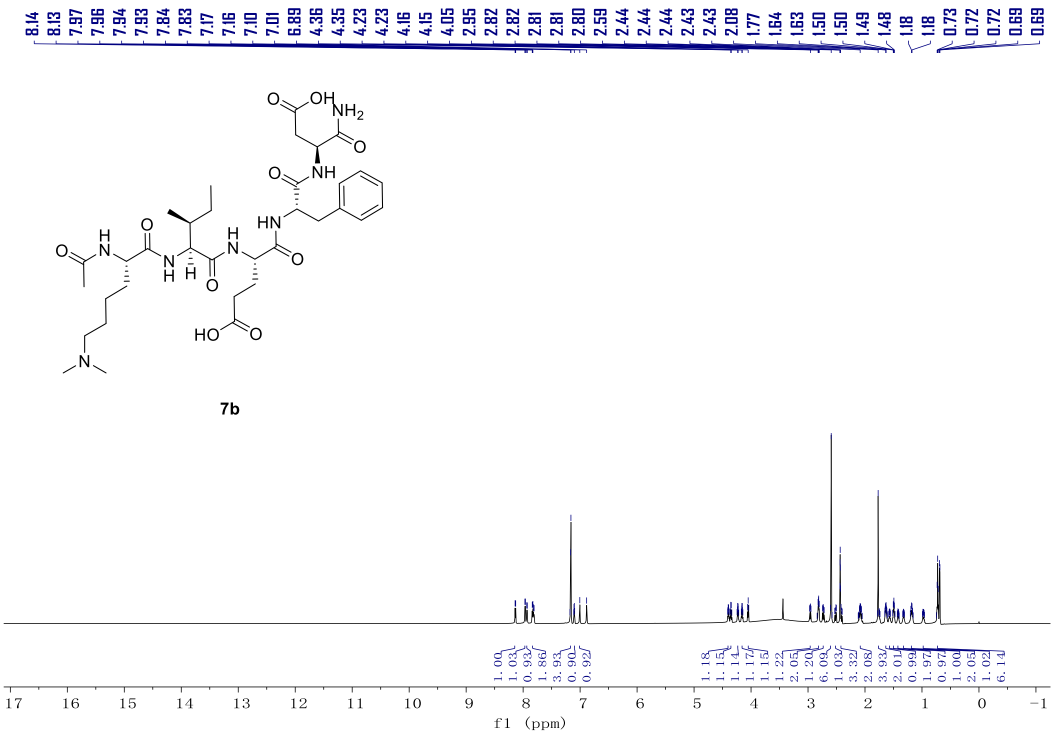


[1]H NMR (800 MHz) spectrum of **7b** in DMSO-d6.

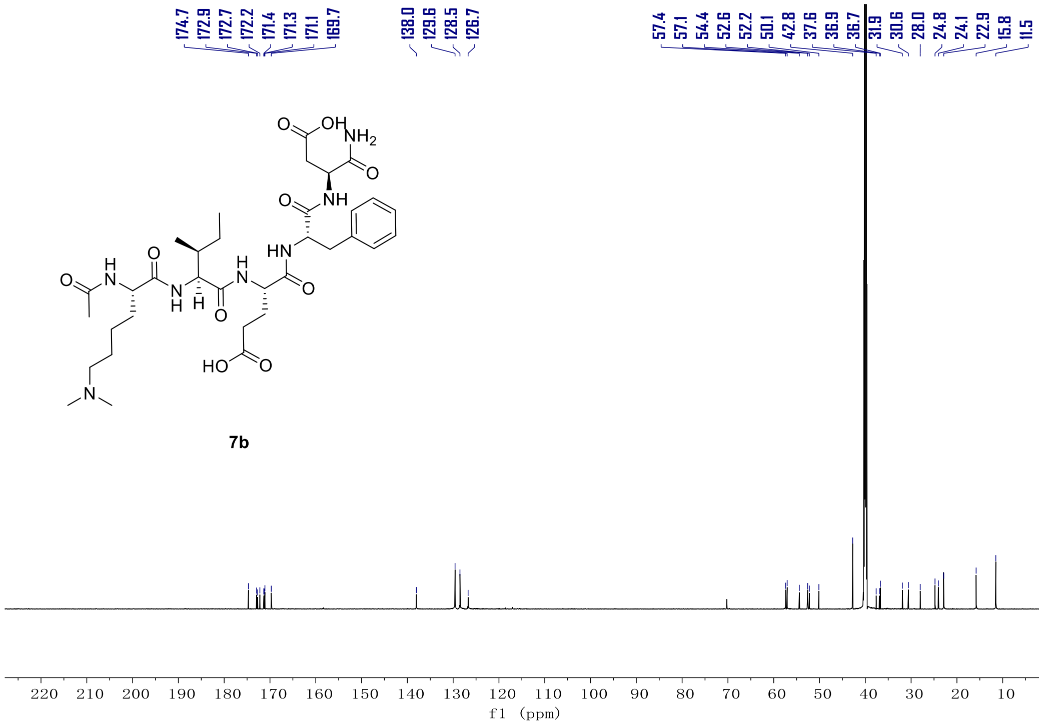


$^{13}$C NMR (201 MHz) spectrum of **7b** in DMSO-$d_6$.

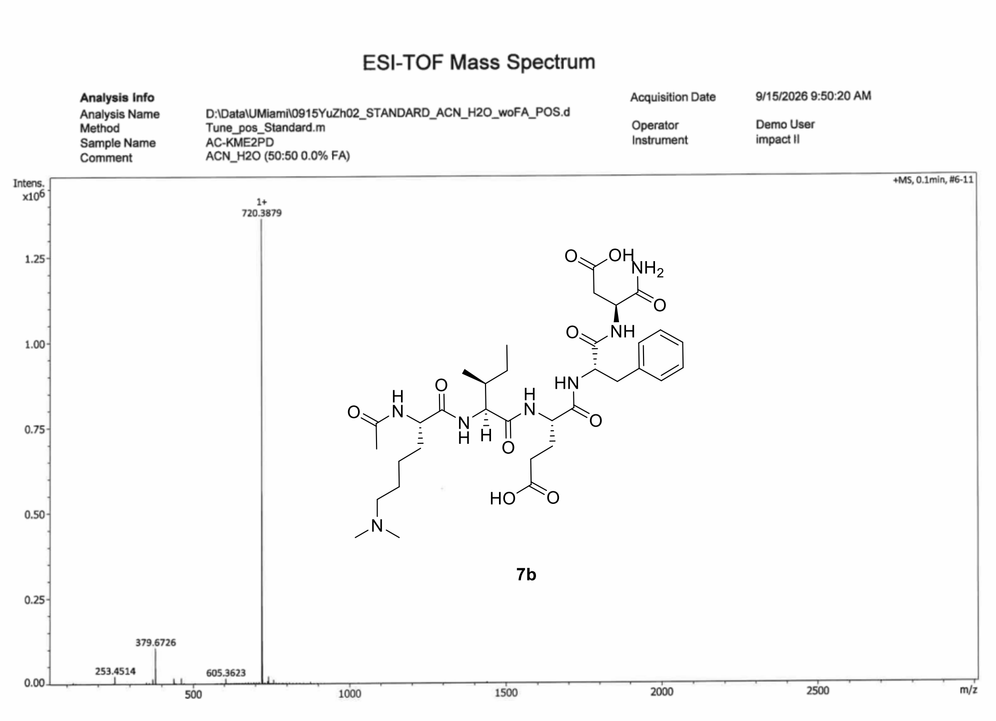


HR-MS spectrum of **7b**

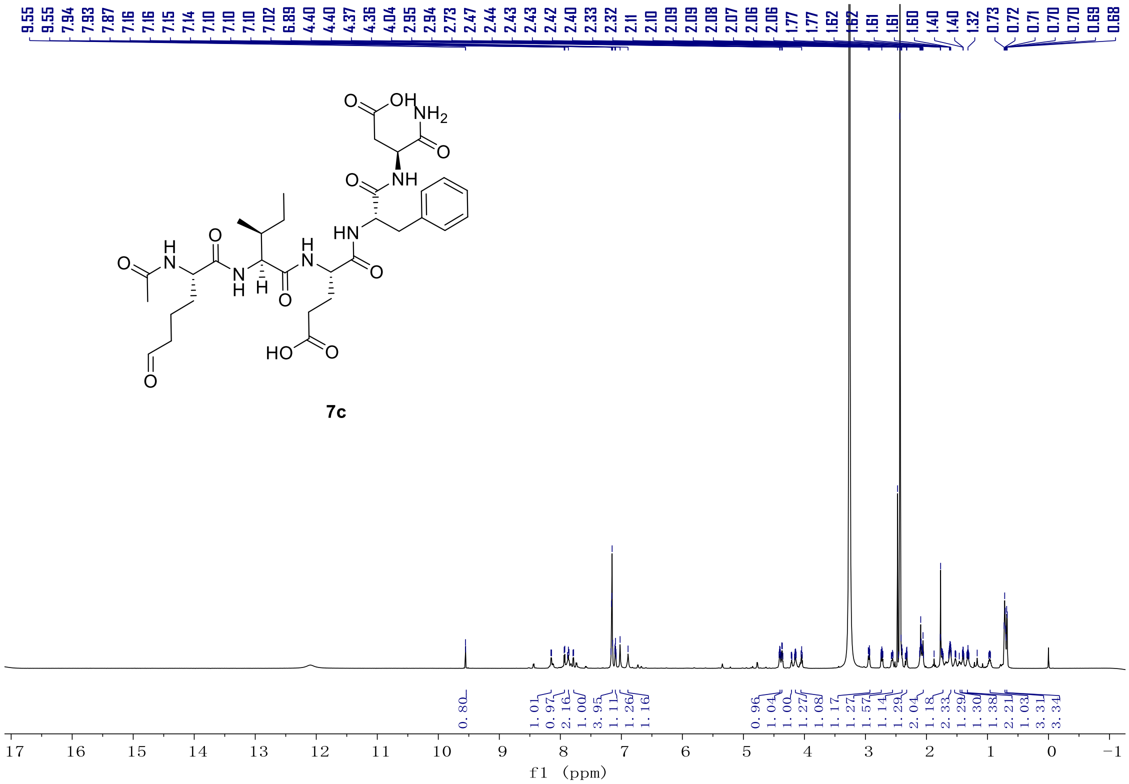

$^1H$ NMR (800 MHz) spectrum of **7c** in DMSO-d6.

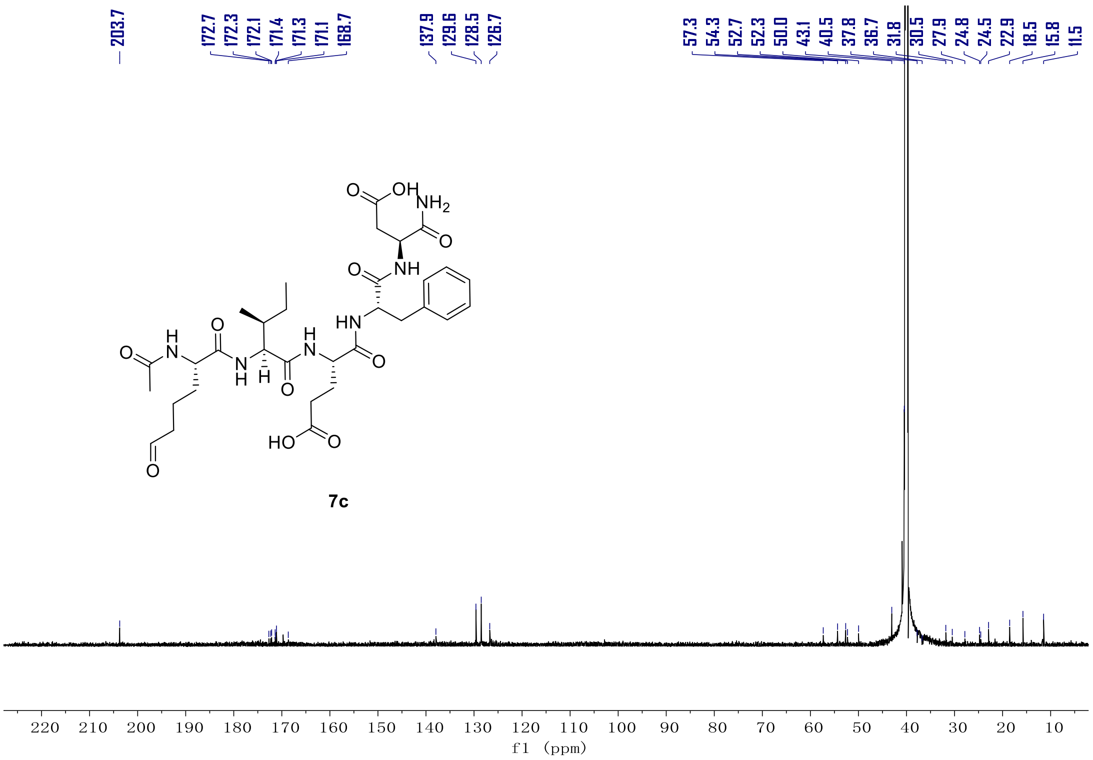

$^{13}C$ NMR (201 MHz) spectrum of **7c** in DMSO-d6.

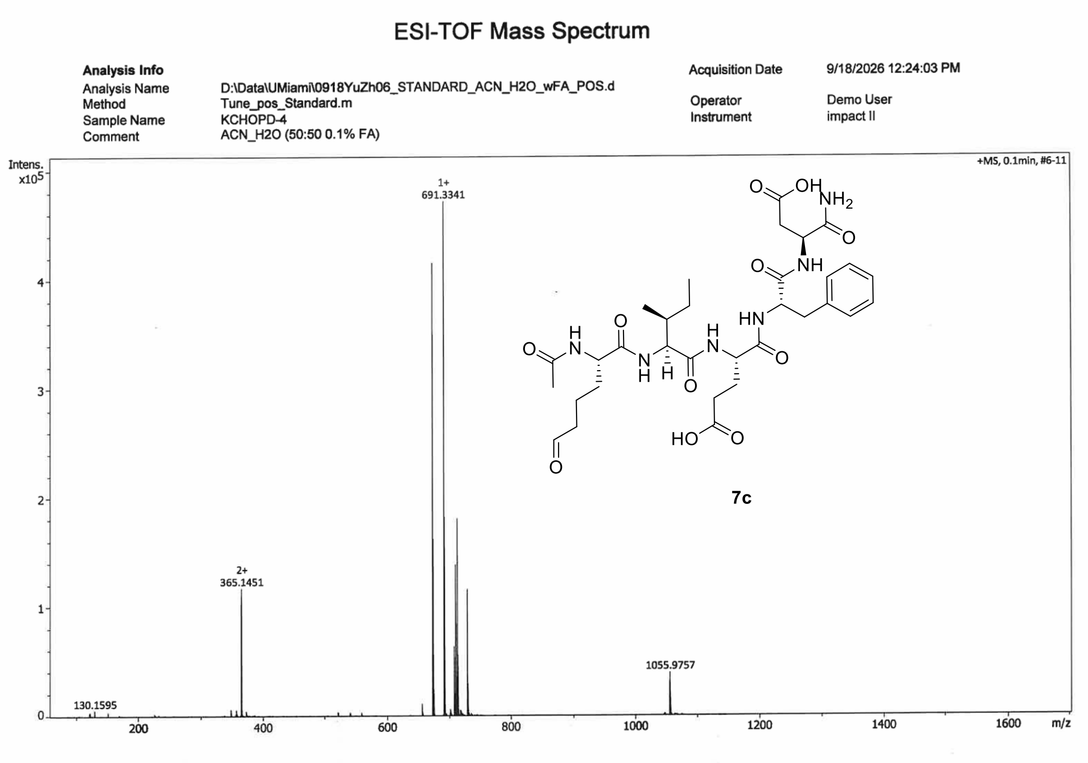


HR-MS spectrum of **7c**

#### 8. References

1. Kong, A. T.; Leprevost, F. V.; Avtonomov, D. M.; Mellacheruvu, D.; Nesvizhskii, A. I., MSFragger: ultrafast and comprehensive peptide identification in mass spectrometry-based proteomics. Nat. Methods 2017, 14, 513-520.
2. da Veiga Leprevost, F.; Haynes, S. E.; Avtonomov, D. M.; Chang, H. Y.; Shanmugam, A. K.; Mellacheruvu, D.; Kong, A. T.; Nesvizhskii, A. I., Philosopher: a versatile toolkit for shotgun proteomics data analysis. Nat. Methods 2020, 17, 869-870.
3. Yu, F.; Haynes, S. E.; Teo, G. C.; Avtonomov, D. M.; Polasky, D. A.; Nesvizhskii, A. I., Fast Quantitative Analysis of timsTOF PASEF Data with MSFragger and IonQuant. Mol. Cell Proteomics 2020, 19, 1575-1585.
4. Chen, T.; Ma, J.; Liu, Y.; Chen, Z.; Xiao, N.; Lu, Y.; Fu, Y.; Yang, C.; Li, M.; Wu, S.; Wang, X.; Li, D.; He, F.; Hermjakob, H.; Zhu, Y., iProX in 2021: connecting proteomics data sharing with big data. Nucleic Acids Res. 2022, 50, D1522-D1527.
5. Ma, J.; Chen, T.; Wu, S.; Yang, C.; Bai, M.; Shu, K.; Li, K.; Zhang, G.; Jin, Z.; He, F.; Hermjakob, H.; Zhu, Y., iProX: an integrated proteome resource. Nucleic Acids Res. 2019, 47, D1211-D1217.
